# Estimation of Speciation Times Under the Multispecies Coalescent

**DOI:** 10.1101/681023

**Authors:** Jing Peng, David L. Swofford, Laura Kubatko

## Abstract

**Motivation:** The multispecies coalescent model is now widely accepted as an effective model for incorporating variation in the evolutionary histories of individual genes into methods for phylogenetic inference from genome-scale data. However, because model-based analysis under the coalescent can be computationally expensive for large data sets, a variety of inferential frameworks and corresponding algorithms have been proposed for estimation of species-level phylogenies and associated parameters, including speciation times and effective population sizes.

**Results:** We consider the problem of estimating the timing of speciation events along a phylogeny in a coalescent framework. We propose a maximum *a posteriori* estimator based on composite likelihood (*MAP*_CL_) for inferring these speciation times under a model of DNA sequence evolution for which exact site pattern probabilities can be computed under the assumption of a constant *θ* throughout the species tree. We demonstrate that the *MAP*_CL_ estimates are statistically consistent and asymptotically normally distributed, and we show how this result can be used to estimate their asymptotic variance. We also provide a more computationally efficient estimator of the asymptotic variance based on the nonparametric bootstrap. We evaluate the performance of our method using simulation and by application to an empirical dataset for gibbons.

**Availability and implementation:** The method has been implemented in the *PAUP\** program, freely available at https://paup.phylosolutions.com for Macintosh, Windows, and Linux operating systems.

**Supplementary information:** Supplementary data are available at *Bioinformatics* online.

## 1 Introduction

Though numerous methods have recently been developed for estimating species tree topologies, methods for estimating the associated speciation (divergence) times are less well-developed. Traditionally, divergence times have been obtained using maximum likelihood (ML) estimates of branch lengths from a concatenated alignment, but this approach has been shown to produce systematic errors because it fails to account for variation in gene genealogies and their associated gene divergence times. As a result, some node ages are overestimated while others are underestimated (Ogilvie *et al*., 2017).

In contrast to concatenation, coalescent-based methods explicitly model variation in individual gene genealogies under the multispecies coalescent (MSC) model (Hudson, 1983; Rannala and Yang, 2003). Several widely used implementations provide estimates of either speciation times or internal branch lengths in addition to estimating the species tree topology. Of the methods that infer species trees from multilocus data using estimated gene trees (“summary statistic methods” or “summary methods”), *ASTRAL* (Sayyari and Mirarab, 2016) and *MP-EST* (Liu *et al*., 2010) can also provide estimates of internal branch lengths in coalescent units. Branch-length estimates from both of these methods are statistically consistent (Liu *et al*., 2010; Sayyari and Mirarab, 2016), but consistency has only been shown to hold when the input data consist of an unbiased sample of true gene trees. Even if input gene trees are estimated using a statistically consistent method (e.g., ML), a proof of consistency for branch length estimation would either need to allow gene lengths to go to infinity (violating the MSC assumption of no intralocus recombination) or demonstrate the absence of any small sample bias in topology estimation, even though such a bias is known to exist for ML (Swofford *et al*., 2001; Roch *et al*., 2019). In fact, both *ASTRAL* and *MP-EST* have been shown to underestimate internal branch lengths when gene tree estimation error increases (Sayyari and Mirarab, 2016). In addition, Yang (2002) showed that phylogenetic errors inflate the probability of incongruent gene trees and lead to biased estimation of internal branch lengths.

An alternative to summary methods is a fully Bayesian approach that jointly estimates the species tree topology, speciation times, and effective population sizes using the complete sequence data without first estimating gene trees for each locus. These methods are implemented in *\*BEAST* /*StarBEAST2* (Heled and Drummond, 2010; Ogilvie *et al*., 2017) and *BPP* (Yang and Rannala, 2014; Rannala and Yang, 2017) for multilocus sequence data, and *SNAPP* (Bryant *et al*., 2012) for biallelic SNP data. *StarBEAST2* and *BPP* differ in the prior distributions assumed for the species tree, the range of evolutionary models supported, and details of the Markov chain Monte Carlo (MCMC) strategies used to sample from the posterior distribution. Bayesian methods have the advantage of using all of the data, but due to reliance on MCMC, they can be very slow for data sets with a large number of species and/or genes.

A third class of methods infers species trees directly from the sequence data without requiring prior estimation of gene trees for each locus. The most widely used example of this class, SVDQuartets (Chifman and Kubatko, 2014), is much faster than fully Bayesian approaches, but it can only estimate the topology of the species tree. Here we use some of the theory underlying SVDQuartets (Chifman and Kubatko, 2015) to derive an estimator for node ages under the MSC model and the JC69 DNA substitution model (Jukes and Cantor, 1969), assuming a molecular clock. Similar approaches have been used to obtain parameter estimates for two (Andersen *et al*., 2014) or three (Zhu and Yang, 2021) species for fixed phylogenies. Our estimator is not directly connected to SVDQuartets, apart from being a quartet-based method that operates under the MSC assumptions. As such, it can be used to estimate speciation times on trees obtained using any method, although it is especially relevant for SVDQuartets, which does not intrinsically provide estimates of node ages or branch lengths.

Our proposed node-age estimator differs from the two-step summary methods described above by eliminating the step of estimating gene trees. Instead, it uses the sequence data directly to compute composite likelihoods based on the fit of observed site pattern probabilities to their expectations under the MSC model. It thus captures variability due to both the coalescent process and the mutation process in a way that the two-step summary methods do not. However, our method diverges from a full likelihood method in that site pattern frequencies are calculated by pooling sites across all loci. Consequently, when multilocus data are used as input, the coalescent variation among gene trees and the mutational variance among sites of the same locus are confounded. On the other hand, by avoiding MCMC, our method gains a strong computational advantage over current fully Bayesian methods. Here, we prove that this estimator is statistically consistent and argue that it is asymptotically normally distributed. Though the uncertainty in the estimator can be quantified by the theoretical asymptotic variance predicted by our normality result, we find that use of the nonparametric bootstrap provides a more computationally efficient estimate of the variance of the estimates. The performance and computational cost associated with our method of speciation time estimation is compared with *BPP* using simulated datasets. We use a genome-scale dataset for gibbons (Carbone *et al*., 2014; Veeramah *et al*., 2015; Shi and Yang, 2018) to demonstrate the performance of our estimator for empirical data.

## 2 Methods

### 2.1 Time scales

Speciation times (node ages) represent the amount of time elapsed between each ancestral node and the present. The amount of time between any pair of nodes is typically measured in either “coalescent units” (number of generations scaled by 2*N*_*e*_, where *N*_*e*_ is the effective population size) or “mutation units” (expected time to accumulate one mutation per site assuming a mutation rate *µ*, defined as the expected number of mutations per site per generation). For speciation times *τ*, these units can be interconverted using *τ*_coal_=(2*/θ*)*τ*_mut_ or *τ*_mut_=(*θ/*2)*τ*_coal_, where *θ* = 4*N*_*e*_*µ*. When we assume that a single value of *θ* applies to the entire tree, age estimates in either units satisfy the molecular clock. However, if *θ* is allowed to vary over the tree (e.g., a separate *θ* parameter for each branch), ages measured in coalescent units are no longer proportional to time in any sense, and mutation units are more appropriate. For generality, we prefer to use *τ*_mut_, but there are situations where *τ*_coal_ is more convenient. In order to simplify our notation below, we drop the subscript on *τ*, and make it clear in the text which units are being used.

### 2.2 Site pattern probabilities

In a 4-taxon species tree, there are 4^4^=256 possible site patterns. Chifman and Kubatko (2015) show that for the JC69 (Jukes and Cantor, 1969) model and a 4-leaf species tree containing species *a, b, c*, and *d*, each site pattern probability 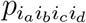 for a specific observation *i*_*a*_*i*_*b*_*i*_*c*_*i*_*d*_, *i*_*j*_∈{A,C,G,T}, can be written as a function of the mutation-scaled population size (*θ*) parameter and the node ages (***τ***) in the tree (in coalescent units).

Under this model as well as the molecular clock assumption, the rooted symmetric 4-leaf species tree ((*a, b*), (*c, d*)) has 9 distinct site pattern probabilities: *p*_1_=*p*_*xxxx*_, *p*_2_=*p*_*xxxy*_=*p*_*xxyx*_, *p*_3_=*p*_*xyxx*_=*p*_*yxxx*_, *p*_4_=*p*_*xyxy*_ =*p*_*yxxy*_, *p*_5_=*p*_*xxyy*_, *p*_6_=*p*_*xyxz*_ =*p*_*yxxz*_ =*p*_*xyzx*_=*p*_*yxzx*_, *p*_7_=*p*_*xxyz*_, *p*_8_=*p*_*yzxx*_, and *p*_9_=*p*_*xyzw*_, where *x, y, z* and *w* denote different nucleotides. For example, *p*_*xxxx*_ includes the site patterns *p*_AAAA_, *p*_CCCC_, *p*_GGGG_ and *p*_TTTT_, which have identical probabilities under the model, and *p*_xxxy_ includes the site patterns *p*_AAAC_, *p*_AAAG_, *p*_AAAT_, *p*_CCCA_, etc. These expressions provide probabilities for individual sites in a nucleotide sequence alignment given a species tree under the MSC model. Using the same notation, the rooted asymmetric 4-leaf species tree (*a*, (*b*, (*c, d*))) has 11 distinct site pattern probabilities: *p*_1_=*p*_*xxxx*_, *p*_2_=*p*_*xxxy*_ =*p*_*xxyx*_, *p*_3_=*p*_*xyxx*_, *p*_4_=*p*_*yxxx*_, *p*_5_=*p*_*xyxy*_ =*p*_*yxxy*_, *p*_6_=*p*_*xxyy*_, *p*_7_=*p*_*xyxz*_ =*p*_*xyzx*_, *p*_8_=*p*_*yxxz*_ =*p*_*yxzx*_, *p*_9_=*p*_*xxyz*_, *p*_10_=*p*_*yzxx*_, and *p*_11_=*p*_*xyzw*_.

We use 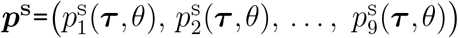 to denote the 9 different site pattern probabilities arising from the symmetric 4-taxon species tree, augmenting the notation above to indicate the dependence of the site pattern probabilities on the quantities *θ* and ***τ***. Likewise, 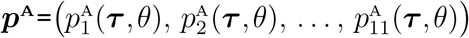 denotes the 11 distinct site pattern probabilities from the asymmetric 4-taxon species tree. In an alignment of length *M*, the site pattern frequencies for these classes can be modeled as a multinomial random variable under the assumption that the observed sites are independent, conditional on the species tree:

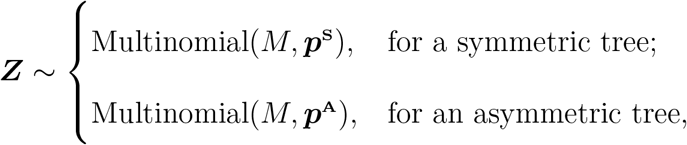

where ***Z*** is the vector of site pattern counts for the 9 or 11 distinct classes.

### 2.3 Maximum *a posteriori* estimation based on composite likelihood

We can split a tree of arbitrary size into the subtrees induced by each quartet of four leaves, and write the likelihood of the observed site pattern frequencies for each quartet. For example, in the 5-leaf species tree in Figure 1, we can consider all sets of 4 tips, to get 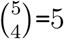 different quartets. For any quartet *i*, each site in an alignment of length *M* can be classified into one of *n*_*i*_ distinct site patterns, where *n*_*i*_ equals 11 if the quartet induces an asymmetric subtree of the full tree (*i*∈{1, 2} in this case) or 9 if it induces a symmetric subtree (*i*∈{3, 4, 5}). Our classification of each site into one of the possible site patterns assumes that the sites are unlinked observations from the species tree under the MSC model. We refer to data that satisfy this assumption as Coalescent Independent Sites (CIS) data. Multilocus data will not satisfy this assumption, as sites sampled within the same gene are not independent observations from the species tree. However, we will show that our proposed methodology performs well for multilocus data as well as for CIS data. Some justification for this claim is given in Wascher and Kubatko (2021).

**Fig. 1.**
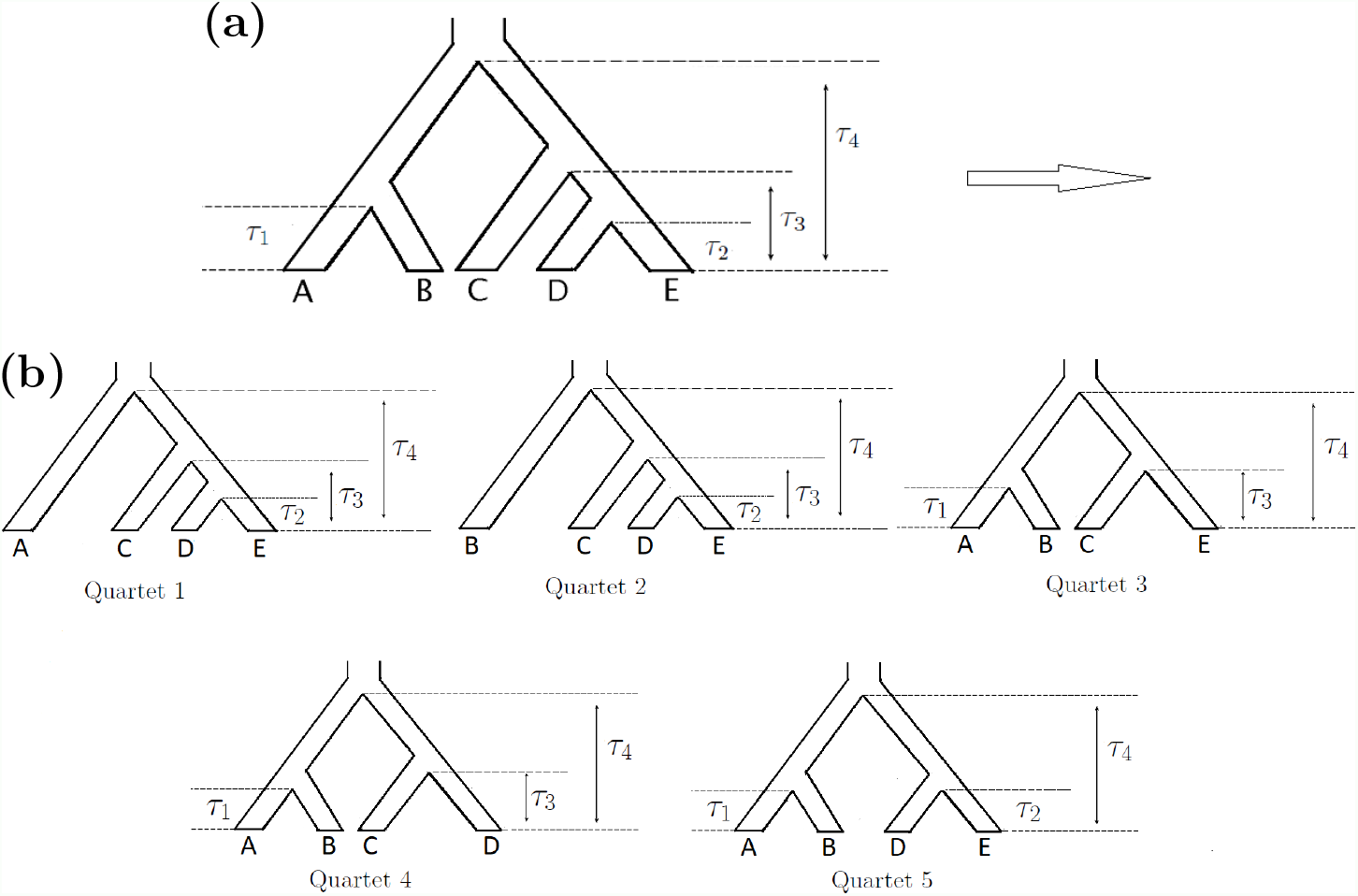
The 5-leaf species tree (a) can be split into the 5 different 4-leaf subtrees (b), shown with speciation times marked.

For each site *m, m*=1, 2, …, *M*, and each quartet *i, i*=1, 2, …, 5, define 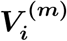 to be the random vector of length *n*_*i*_ that contains a 1 in the *j*^*th*^ entry if site pattern *j* is observed at that site and 0 in all other entries, and 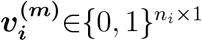 represent the corresponding observed data. Let 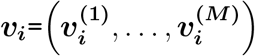 denote the observed data across all *M* sites, and let (*u*_*i*_)_*j*_ be the *j*^*th*^ entry of the vector 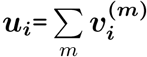, which counts the number of times site pattern *j* is observed. Letting *f*_*i*_(***v***_*i*_|***τ***, *θ*)=Pr(***v***_*i*_ | ***τ***, *θ*), the likelihood for quartet *i* can then be expressed as a function of *θ* and ***τ*** :

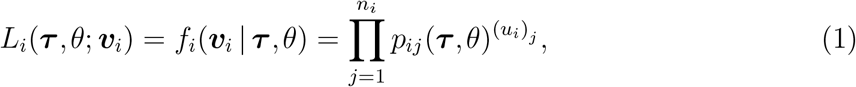

where *p*_*ij*_ is the *j*^*th*^ entry in either ***p***^**S**^ or ***p***^**A**^ for quartet *i*, depending on whether the subtree induced by this quartet is symmetric or asymmetric, respectively.

Importantly, the subtrees induced by different quartets are not independent, and computing a true likelihood would require accounting for the correlation structure among quartets. Therefore, we instead use *composite likelihood* —the product of the individual likelihoods for all possible quartets despite their non-independence. Note that composite likelihood is also often referred to in the statistical and biological literature as *pseudolikelihood* or *approximate likelihood* (see Varin *et al*., 2011, for a review of the history of composite likelihood methods).

A maximum composite likelihood estimator (MCLE) based on (1) would optimize the function

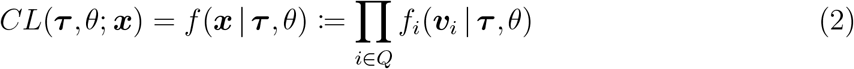

where *Q* is the set of all possible quartets. The vector ***x***=(***x***^(1)^, …, ***x***^(*M*)^) is defined similarly to the ***v***_***i***_, but for the entire tree; its dimension depends on the number of possible distinct site patterns on a 5-leaf tree. Specifically, each vector ***x***^(*m*)^ records which of the possible distinct site patterns on a tree of 5 tips is observed at site *m*, while the corresponding 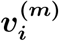 stores the indicators of the *n*_*i*_ site patterns for the *i*^*th*^ quartet of this tree at site *m*.

Instead of using the MCLE, however, we prefer to estimate ***τ*** and *θ* via Bayesian maximum *a posteriori* (MAP) estimation (e.g., Bassett and Deride, 2019). MAP estimation has two advantages. First, it allows incorporation of prior knowledge into the estimate as for the fully Bayesian methods discussed above. Perhaps more importantly, weighting the likelihood by the priors improves the computational efficiency and stability of the optimization algorithms by reducing the flatness of the optimality surface in regions of the parameter space that have very low likelihood.

With inclusion of the priors, the (unnormalized) posterior density function becomes

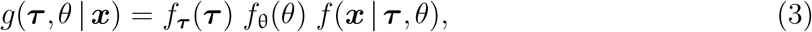

where *f*_***τ***_ and *f*_θ_ are the prior density functions for the vector of node ages ***τ*** and the shared *θ* parameter, respectively. We follow *BPP* in using inverse-gamma priors for the root age (= *τ*_R_) and *θ* parameters. The rank-ordered non-root node ages are assigned a uniform Dirichlet prior, which simply incorporates a constant scaling factor into the joint prior on ***τ***. By maximizing the log posterior density

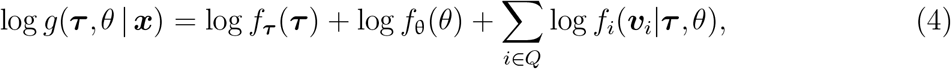

we obtain our maximum *a posteriori* estimator *MAP*_CL_:

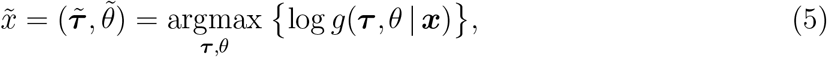

with the “CL” subscript signifying that a composite-likelihood term is used in (2) rather than a true likelihood.

Using results from Miller (2019) and Arnold and Strauss (1991), we can prove that the *MAP*_CL_ estimator is statistically consistent and we argue further that it is also asymptotically normally distributed (detailed proofs can be found in Section 1 of the Supplemental Material):

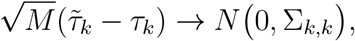

where Σ_*k,k*_ is the cell in the *k*^*th*^ row and column of the variance-covariance matrix

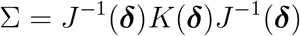

and

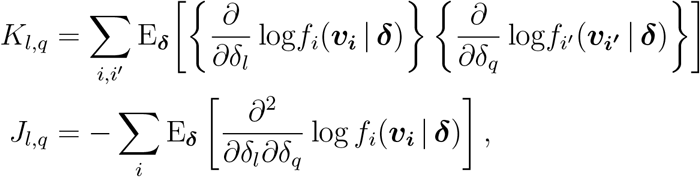

where 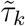 is the *k*^*th*^ component of the *MAP*_CL_ estimator 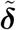, and *f*_*i*_(***v***_***i***_ | ***δ***) is the density function for indicator variable ***v***_***i***_ conditional on the parameters ***δ***. Furthermore, we can use the observed data to approximate *J* and *K*:

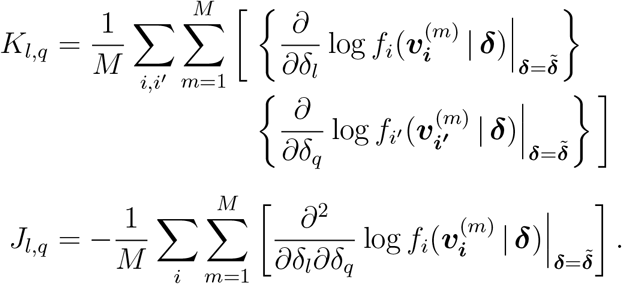

Computation of the asymptotic variance above requires inversion of a matrix that becomes large as the number of parameters increases, which may become problematic. As an alternative, we can use a bootstrap estimator to measure the variance of the *MAP*_CL_ estimator. In this approach, a bootstrap replicate is obtained by resampling the columns, i.e., site patterns in the original DNA sequences, using the following steps:

1. Obtain a bootstrap sample by randomly selecting *M* columns (with replacement) from the original sequence alignment, creating a data set of the same size as the original data;
2. Repeat step 1 *B* times to get a full set of bootstrap samples;
3. For each of the bootstrap samples created in steps 1 and 2, redo the analysis to compute the estimates 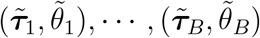, and calculate the sample variance of the estimates, 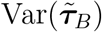 and 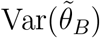.

All of the methods described herein are implemented in the *PAUP\** program written by DLS (https://paup.phylosolutions.com), where they are accessed using the `qage` command (type `help qage;` at the command prompt for a description of the available options). A detailed explanation of the implementation, including parameterizations, mathematical details for likelihood and gradient evaluations, optimization strategies, and validation, is provided in the “Implementation of qAge in *PAUP\**” document contained in the Supplemental Material.

### 2.4 Computer simulation study

We first use simulation to assess the statistical consistency and asymptotic normality of the *MAP*_CL_ estimator and to compare the two methods of measuring uncertainty (calculation of the theoretical asymptotic variance *vs*. bootstrapping). While many methods for inferring species-level phylogenies are based on multilocus data, the theory in the previous section applies specifically to CIS data (unlinked sites arising from the MSC model). For CIS data, the site patterns in the sequences constitute independent draws from the distribution characterized by the MSC and nucleotide substitution models (Chifman and Kubatko, 2015), conditional on the species tree, whereas for multilocus data, all sites at a locus are assumed to have evolved on the same genealogy and are not independent of other sites at the same locus. However, a straightforward argument, similar to that of Wascher and Kubatko (2021) for the SVDQuartets method, can be made that methods developed for CIS data are also valid for multilocus data, and we therefore consider both data types here. Although CIS data are not ordinarily collected in practice, it useful to examine the performance of the method when data are simulated directly from its underlying model.

To examine the properties of the *MAP*_CL_ estimator, we thus simulated two types of data: (1) unlinked CIS data (each site evolves on its own own tree drawn randomly from the distribution of gene trees expected for the true simulation parameters under the MSC model), and (2) multilocus data (a sequence of length *l* is simulated for each locus on an underlying gene tree drawn randomly from the expected gene tree distribution). The simulations were performed as follows:

1. Generate gene tree samples under the MSC model based on a specified input species tree;
2. Generate DNA sequences of length *l* for each gene tree under the JC69 model (*l* = 1 for CIS data);
3. Choose prior distributions for the parameters;
4. Compute the site pattern frequencies for all possible quartets and maximize the log posterior density to obtain node age estimates using the *MAP*_CL_ estimator and estimate their theoretical asymptotic variances;
5. Resample the simulated sequences to get *B* bootstrap replicates, and compute the sample variance of the estimates via bootstrapping, as described in the previous section (for the multilocus datasets, a two-level bootstrap is conducted where we first take a bootstrap sample of genes followed by independent bootstrap resampling of sites within each gene);
6. Repeat steps 1–4 *D* times to obtain node age estimates and estimate variances using both theoretical asymptotic calculations and bootstrapping.

All steps in the simulations were performed using the simulation module and qage command in *PAUP\**. In step 1, two different model species trees were defined: a 5-leaf tree and a 6-leaf tree (Figure 2). Population-size and mutation-rate parameters were set so that *θ*=0.002 (constant throughout the tree). Speciation times were assigned (in mutation units) as (*τ*_1_, *τ*_2_, *τ*_3_, *τ*_4_) = *b* · (0.0005, 0.0005, 0.001, 0.0015) for the 5-tip model tree and (*τ*_1_, *τ*_2_, *τ*_3_, *τ*_4_, *τ*_5_) = *b* · (0.0005, 0.0005, 0.001, 0.0015, 0.002) for the 6-tip model tree. Setting *b* ≠ 1 stretches or shrinks the tree while still satisfying the molecular clock assumption; we used *b* = 1, 2, and 4 for our simulations. In step 2, the gene length *l* was set to 1 for CIS data (i.e., we simulated 100,000 genealogies with one DNA site for each). For multilocus data, we simulated 10,000 genes, each of length *l*=100. We note that this simulation condition results in relatively large data sets, which are useful for comparing the performance of our method to the theoretical predictions derived above. In step 3, we assigned diffuse inverse-gamma priors, parameterized as (mean, standard deviation): *IG*(0.003, 1.0) for *θ* and *IG*(1.0, 1.0) for the age of root. In the parameterization used by BPP, these are *IG*(*α*=3, *β*=0.006) for *θ* and *IG*(*α*=3, *β*=2) for the age of root. In steps 5 and 6, we chose *B*=100 and *D*=100, respectively.

**Fig. 2.**
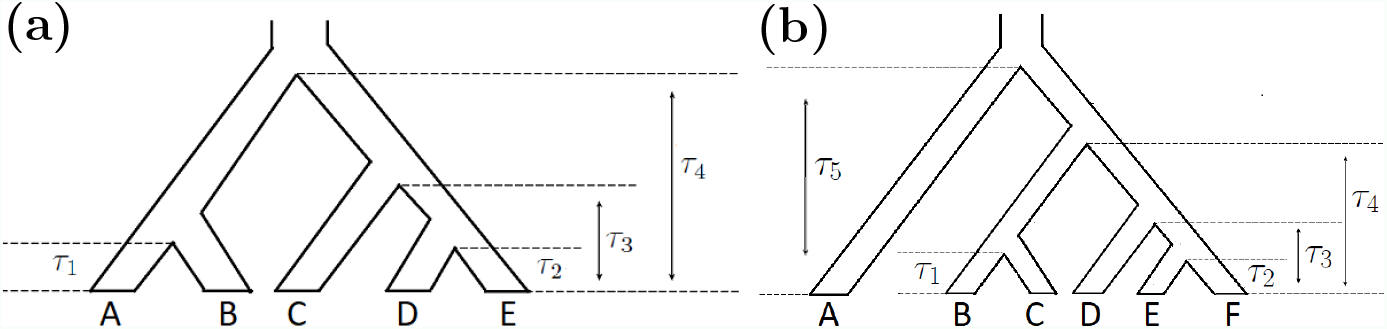
Two different model species trees with speciation times used as parameters for the simulation process: (a) 5-species tree. (b) 6-species tree.

The *MAP*_CL_ estimator is also applicable when multiple lineages are sampled for each tip species (see Supplemental Material: “Implementation”). To evaluate the performance of this option, *PAUP\** was instructed to generate gene tree samples using the same 5-leaf model species tree and parameter settings as above, but with two lineages for both species D and species E. We then used *PAUP\** to simulate and analyze the multilocus data to estimate parameter values and their variances.

We carried out an additional simulation to compare the performance of the *MAP*_CL_ estimator in *qAge* with *BPP*, again using the simulation module in *PAUP\** (which provides an interface to invoke *BPP* from a Nexus file). We simulated multilocus data with 2,000 genes each of length 100, for trees with *K* tips (*K*=7, 8, …, 15, 20), with *θ* set to 0.002. The trees used for simulation are included in Figures S16 and S17.

We ran *BPP* and *qAge* analyses with inverse-gamma priors *IG*(*µ, σ*) for *θ* and the age of the root node *R* (*τ*_R_). Specifically, we used *σ*=1.0 for a diffuse prior and adjusted *µ* such that the prior mean was equal to 5*θ, θ* or *θ/*5 for the *θ* parameter, and 5*h, h*, or *h/*5 for the root age *τ*_R_, where the tree height *h* is defined as the maximum number of branches connecting the tips and the root. For example, in Figure 2, the height of the tree is 3 in (a), and 4 in (b). The details of prior-distribution combinations can be found in Figure 6. We then investigated the impact of the 3 × 3 combinations of the priors on the performance of *BPP* and *MAP*_CL_. To make the comparison in a computationally feasible way, for smaller trees (K=7, 8, 9, 10), we discarded the first 1,000 samples as burnin, and sampled every 50^*th*^ observation. For larger trees with more than 10 tips, we ran the *BPP* analysis 1,000 times longer than *qAge*, and discarded the first 10% of the samples as burnin. 500 observations were sampled equally frequently and used to compute estimates in all cases.

The detailed MCMC configurations and running time can be found in Table S1 and Section S3. In all of the analyses, *BPP* estimates a different *θ* parameter for each branch, whereas the current *qAge* implementation assumes and estimates a single *θ* that applies to the entire tree. After performing analyses with *BPP* and *qAge* for 100 replicates, we quantified the deviation of estimated node ages in mutation units from the true values of the simulation model using the root-mean-square error (RMSE) and mean absolute error (MAE). We calculated the proportion of 95% confidence/credible intervals that included the true parameter value. Finally, for *BPP* analyses, we summarized the percentage of ESS values *>* 200.

### 2.5 Application to gibbon data

We explored the performance of our *MAP*_CL_ estimator in inferring speciation times for empirical data by applying it to a genome-scale dataset previously analyzed by Shi and Yang (2018) for five species of gibbons: *Hylobates moloch* (Hm), *Hylobates pileatus* (Hp), *Nomascus leucogenys* (N), *Hoolock leuconedys* (B), and *Symphalangus syndactylus* (S) (Carbone *et al*., 2014; Veeramah *et al*., 2015). The dataset consists of 11,323 coding loci, each of length 200 bp. Except for the outgroup (O), multiple lineages are included for each species: two for Hm and Hp, and four for N, B, and S. Here, we reanalyze these data with *qAge* (for *MAP*_CL_) and *BPP*.

For both analyses, we fixed the species tree to be that shown in Figure 3. In this example, both programs estimate 5 node age parameters, but *BPP* estimates 10 *θ* parameters while *qAge* estimates a single *θ* value. As recommended by the *BPP* authors (Flouri *et al*., 2018), we use inverse-gamma prior distributions with the *α* parameter set to 3 for both *θ* and for the root age, *τ*_R_. We then chose the value of *β* to match the mean of the distribution used by Shi and Yang (2018), although they assumed gamma rather than inverse-gamma prior distributions in an earlier version of *BPP*. To study sensitivity to the prior, we also conducted analyses with prior means that were five times larger and five times smaller than these values, and looked at all combinations of these prior settings for each parameter, leading to a total of nine prior combinations which we label Settings 1-9 in Table S2 of Section S4 (see also Figure 8). Setting 5 corresponds most closely to the priors used in Shi and Yang (2018): *θ*∼*IG*(0.001, 1.0) and *τ*_R_∼*IG*(0.01, 1.0). For each choice of prior distribution, we repeated the analysis twice, with each replicate run for two weeks, and we sampled every 100^*th*^ observation. All prior settings reached at least 10,000 samples during this time (which corresponds to 10,000 × 100 = 1 million iterations of the algorithm), except for replicate 1 in setting 9, for which 9,205 samples were obtained. After discarding the first 2,000 samples as burnin, samples 2,000-10,000 from both replicates were combined to compute estimates (for setting 9, replicate 1, samples 2,000 - 9,205 were used).

**Fig. 3.**
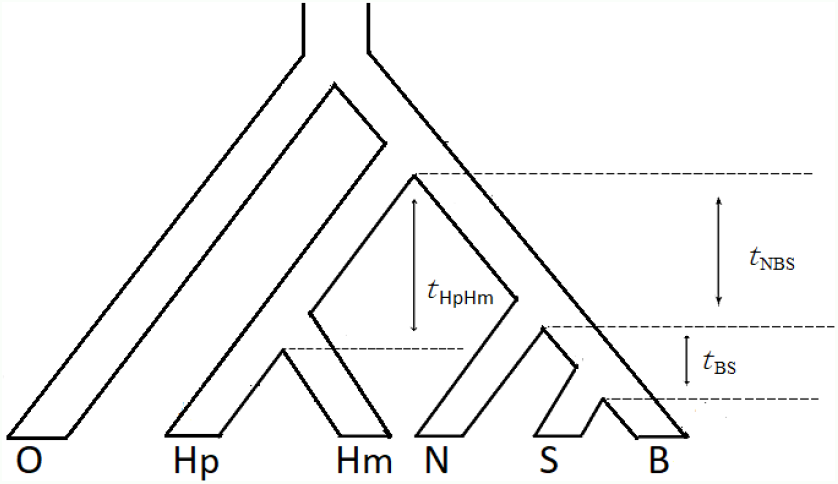
The species tree for the five gibbon species and the outgroup (O=human) with branch length parameters labeled by *T*_*i*_ : *i* = labels for all species descending from the lower node incident to the branch.

For *MAP*_CL_, we enumerated all possible quartets by selecting one lineage per tip species, resulting in 752 quartet likelihoods used to calculate the composite likelihood. Using the same priors as for *BPP*, we estimated the internal branch lengths *t*_*BS*_, *t*_*NBS*_, and *t*_*HpHm*_ (see Figure 3), and the single *θ* parameter; variances were estimated using the bootstrap. Note that although we used the difference between speciation times in this case (i.e., branch lengths rather than node ages), statistical consistency and asymptotic normality can still be shown to hold.

As discussed above, an important assumption for *MAP*_CL_ estimation is that the mutation-scaled population size *θ* is constant throughout the tree. Since this assumption is likely to be violated in practice, we used simulation to check the impact of variable *θ*s on estimation accuracy. To make our simulation realistic, we used the species tree with node ages set to those inferred by *BPP* for the gibbon data of Shi and Yang (2018) (which matches the topology of Figure 2b) and simulated 11,323 loci with 200 bp for each. We conducted simulations as follows:

1. Sample 11 values of the *θ* parameter from an exponential distribution with mean 0.0053; these serve as the “true” *θ*s for the 11 ancestral and extant populations for the gibbon phylogeny of Shi and Yang (2018);
2. Generate 100 replicates of DNA sequences with 11,323 loci, 200 bp for each under the species tree in Figure 2(b). Compute 100 *MAP*_CL_ estimates and compute the mean square errors and absolute errors for the five node ages.
3. Repeat steps 1-2 100 times and compute normalized RMSEs (RMSE/truth*100%) and RMSEs for 100 combinations of *θ* values. We normalize RMSEs to the true parameter values to get a better idea of the amount of error since some branches are very short.

## 3 Results

### 3.1 Simulation study

To assess the statistical properties of the *MAP*_CL_ estimator, we plotted histograms of the 100 *MAP*_CL_ estimates for node ages in the three 5-taxon and the three 6-taxon model trees (Supplemental Material, Section S2 contains figures for all of the simulation settings). As a representative example, Figure 4 shows histograms of the 100 *MAP*_CL_ estimates of node age *τ*_1_ for the three 5-leaf model trees under our simulation conditions. From these plots, we see that the estimates are approximately normal and distributed around the true value, thus supporting our theoretical finding of statistical consistency. Moreover, when we include multiple lineages per tip or analyze multilocus data in the same way, consistency and asymptotic normality still appear to hold. When we increase the number of sites, we see these results even more clearly (see Section S2).

**Fig. 4.**
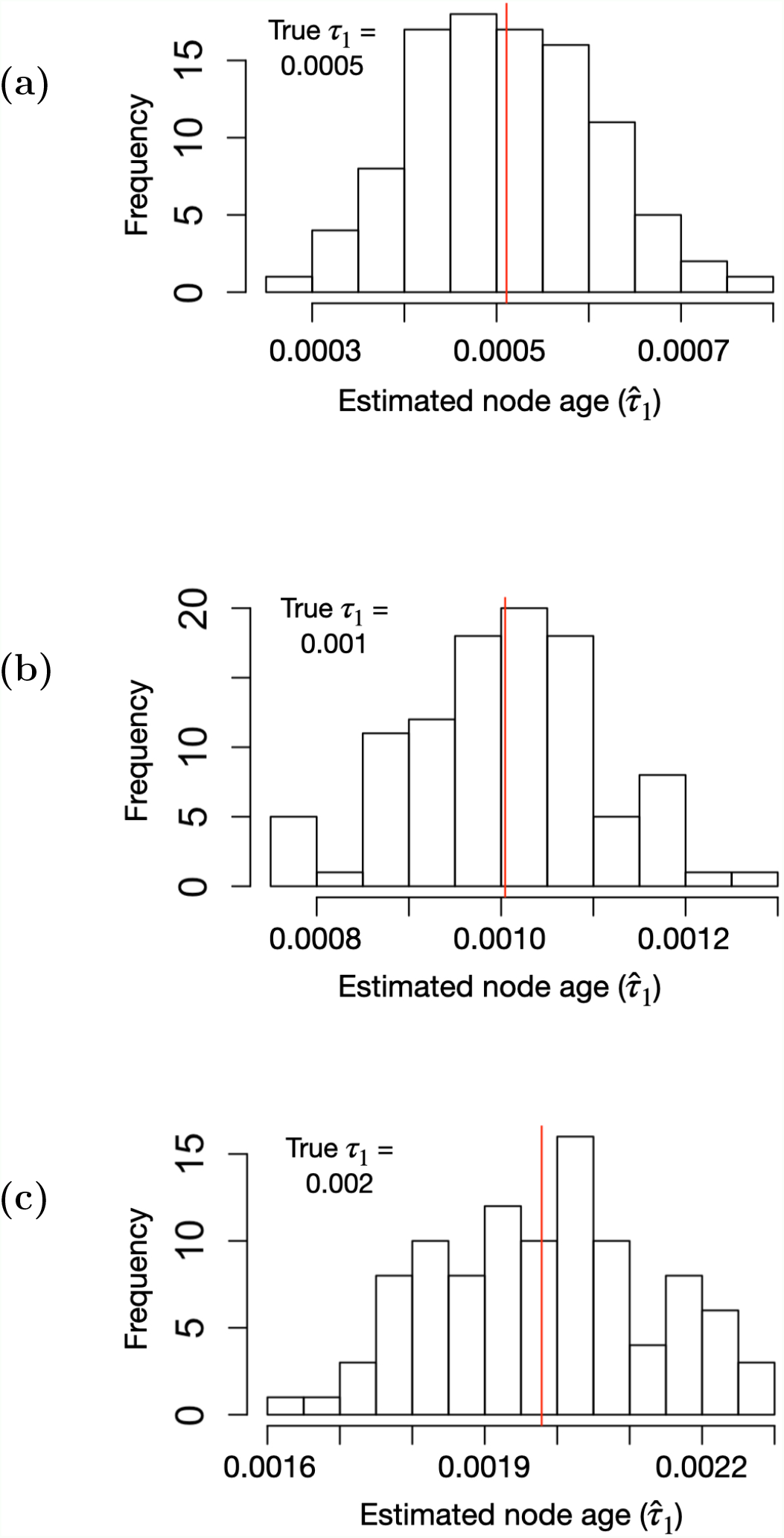
Histograms of 100 *MAP*_CL_ estimates for node age *τ*_1_ (in mutation units) using 100,000 unlinked CIS from the 5-leaf model trees with a single lineage per tip. The red line in each histogram is the sample mean of the 100 *MAP*_CL_ estimates. (a) *τ*_1_ = 0.0005; (b) *τ*_1_ = 0.001; (c) *τ*_1_ = 0.002.

To assess the performance of our method in estimating the uncertainty of the *MAP*_CL_ estimator, Figure 5 shows plots of the 100 variance estimates of the *MAP*_CL_ estimates of node age *τ*_1_ for the three 5-leaf model trees under our simulation conditions. In the unlinked-CIS, single-lineage-per-tip setting, it is immediately clear that in all cases both the bootstrap and asymptotic variance estimates perform similarly and the values are scattered evenly around the sample variance. This approximation can be improved as the number of sites increases (Supplemental Material, section S2.1). These results show both the bootstrap and asymptotic variance estimators are theoretically valid and provide unbiased uncertainty measurements. The bootstrap variance estimator slightly overestimates the uncertainty for multilocus data, while the asymptotic variance estimator shows better performance and less bias for this data type. We also see this tendency toward overestimation in cases under multiple lineages per tip in the 5-taxon and 6-taxon model trees (see Section S2).

**Fig. 5.**
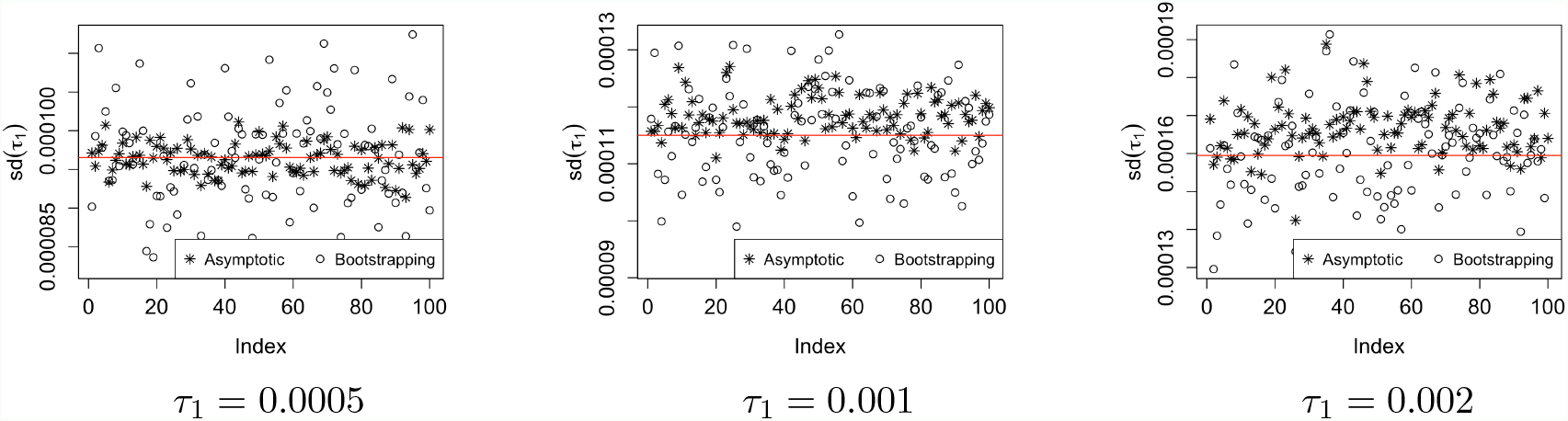
Plots of 100 standard deviation estimates in coalescent units for node age *τ*_1_ using 100,000 unlinked CIS from the 5-leaf model trees with a single lineage per tip. Points denoted by ○ are obtained by bootstrapping. The x-axis is an index for the simulated samples. The red line in the plots is the sample standard deviation of the 100 *MAP*_CL_ estimates.

**Fig. 6.**
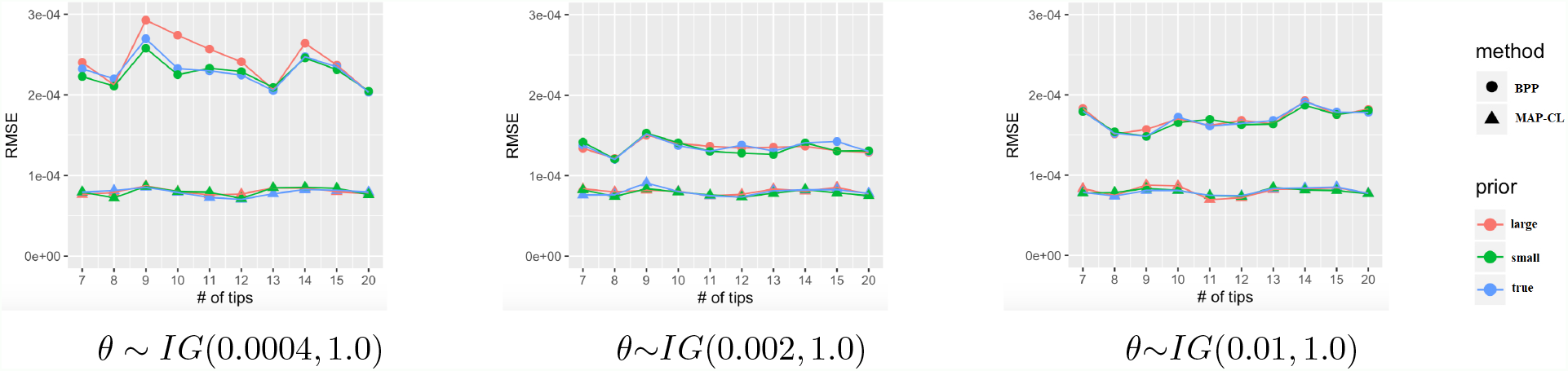
Plots of the RMSE of the node age estimates (in mutation units) for trees with varying numbers of tips. The x-axis shows the number of tips in the tree. Analysis based on two methods (circles – *BPP*, triangles – *MAP*_CL_) is conducted with different priors. In each plot, the “large” prior for the root age, *τ*_R_ ∼ IG(5*h*,1.0), is shown in red; the “small” prior, *τ*_R_ ∼ IG(*h*/5,1.0), is shown in green; and the prior centered at the true value, *τ*_R_ ∼ IG(*h*,1.0), is shown in blue (*h* is the tree height). Panels show the results of analyses using priors centered on different values of *θ*, with the middle panel centered on the true *θ* used for simulation.

We now compare our method with *BPP* and examine the estimation accuracy of both methods. Figure 6 summarizes the RMSE of the node age estimates on trees with different sizes. We find that *MAP*_CL_ estimates speciation times with smaller error than *BPP* and is quite robust to different prior combinations. The estimation error from *BPP* may be partly due to convergence difficulties for some runs, which can be seen from the ESS values (see Figure S18). (*BPP* convergence was better when the prior for *θ* was chosen to have a large mean.) Also, when the data are simulated with a constant *θ* parameter, the fact that *BPP* is estimating a different *θ* parameter for each branch gives an advantage to *MAP*_CL_, which assumes and estimates a single *θ* for the whole tree. Overall, we conclude that after running *BPP* 1,000 times longer than *qAge*, our estimates are comparable or more accurate than those from *BPP* over a wide range of conditions. The results of MAE are similar to those for RMSE (Figure S19).

Additionally, Figure 7 shows the proportion of 95% confidence/credible intervals that include the true parameter value in 100 simulation replicates. Again we note that the performance of *BPP* depends on the choice of prior for *θ*, especially when we compare the coverage probabilities for large trees (which generally did not shown indications of lack of convergence in terms of ESS values) from (a) to (c) in Figure 7. On the other hand, the confidence intervals from *MAP*_CL_ include the true parameter values nearly 95% of the time, which highlights our finding that the bootstrapping produces an consistent estimator of the uncertainty when the number of genes is large enough (Supplemental Material, Figures S7-S15 and Section S2).

**Fig. 7.**
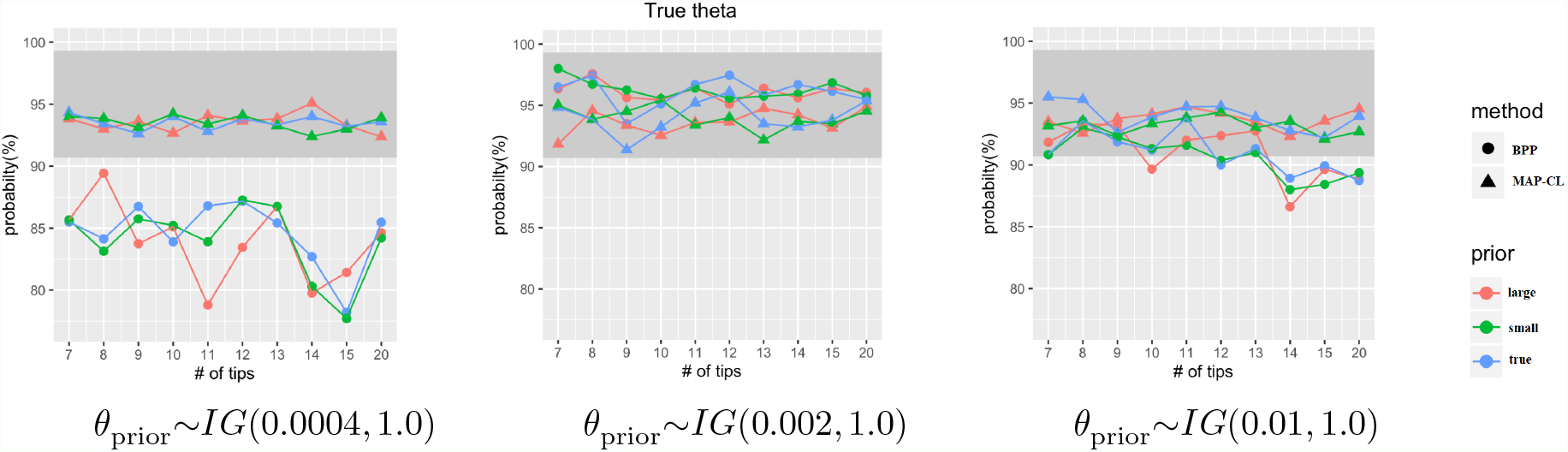
Plots of the percentage of 95% confidence/credible intervals that include the true parameter value. The x-axis gives the tree size (number of tips). Points with different colors give values obtained using the 9 prior combinations for *θ* and *τ*_R_ (see Figure 6). The shaded area gives the expected acceptance region of the coverage proportions in 100 simulation replicates. All summaries were computed using mutation units.

### 3.2 Application to gibbon data

Results (in coalescent units) from *BPP* and *MAP*_CL_ for the choice of prior distributions corresponding most closely to those used by Shi and Yang (2018) are shown in Table 1. We use coalescent units here because they allow a more direct comparison of methods when population sizes and/or mutation rates are allowed to vary across the tree. In addition, Shi and Yang (2018, p. 167) reported that internal branch lengths were more accurately estimated with coalescent units when *θ* was allowed to vary across branches, presumably due to these parameters being poorly estimated in short branches due to the influence of the prior. We have included a comparison of node ages in mutation units in Table S3 for completeness. In Table 1, we see that for *t*_*NBS*_ and *t*_*BS*_, the estimates from *MAP*_CL_ and *BPP* are similar, with wider confidence intervals for *MAP*_CL_ that cover the intervals given by *BPP*, as in the simulation studies. For *t*_*HmHp*_, however, the intervals given by *MAP*_CL_ and *BPP* do not overlap (though the values estimated are similar) and both are similar in width.

**Table 1.** Means and 95% credible (*BPP*) or confidence (*MAP*_CL_) intervals in coalescent units for three internal branch lengths of interest for the gibbon dataset.^a^

| Parameter | BPP | | $MAP_{CL}$ | |
| --- | --- | --- | --- | --- |
|  | mean | 95% HPD interval | mean | 95% CI |
| $t_{NBS}$ | 0.075 | (0.052, 0.096) | 0.097 | (0.065, 0.119) |
| $t_{BS}$ | 0.052 | (0.039, 0.069) | 0.051 | (0.023, 0.076) |
| $t_{HmHp}$ | 2.054 | (1.972, 2.140) | 1.898 | (1.840, 1.945) |

<sup>a</sup> Using the prior distributions matching those used by Shi and Yang (2018) most closely: $\theta \sim IG(0.001, 1.0)$ and $\tau_R \sim IG(0.01, 1.0)$ (our setting 5). For *BPP*, the conversion to coalescent units was carried out as in Shi and Yang (2018) (their Table 5); i.e., we used $2\Delta\tau/\theta$ , where $\tau$ is the relevant node age in mutation units and $\theta$ is the value estimated by *BPP* for that branch. Confidence intervals were determined using the bootstrap.

To examine the sensitivity of these estimators to the prior distribution, we evaluated both estimators under nine different prior settings (Figure 8). Estimates obtained using *MAP*_CL_ are robust to the choice of prior distribution, with little variation across the range of values selected. Conversely, *BPP* is sometimes strongly affected by the choice of priors, most notably for estimation of *t*_*NBS*_ for settings 2 and 3. To examine this more carefully, we made trace plots of all parameters for all replicates and prior choices (see Supplemental Material, section S4). These trace plots show some cases in which the two replicates within a prior setting sampled different values for the entire run (see, e.g., the results for *θ*_*B*_ in Figure S36, setting 4, or for *θ*_*ONBSHmHp*_ in Figure S59, noting the difference in the y-axis values for setting 2). It is also clear that for some settings, *BPP* experienced some difficulty converging, making clear that long runs may be required, even for the relatively straightforward problem of inferring node ages on a fixed 6-taxon species tree with a large data set. In contrast, *qAge* quickly produces stable *MAP*_CL_ estimates that are robust to the choice of prior distribution—it took only 17 seconds to produce an estimate and confidence interval for this data set on a current-generation laptop computer.

**Fig. 8.**
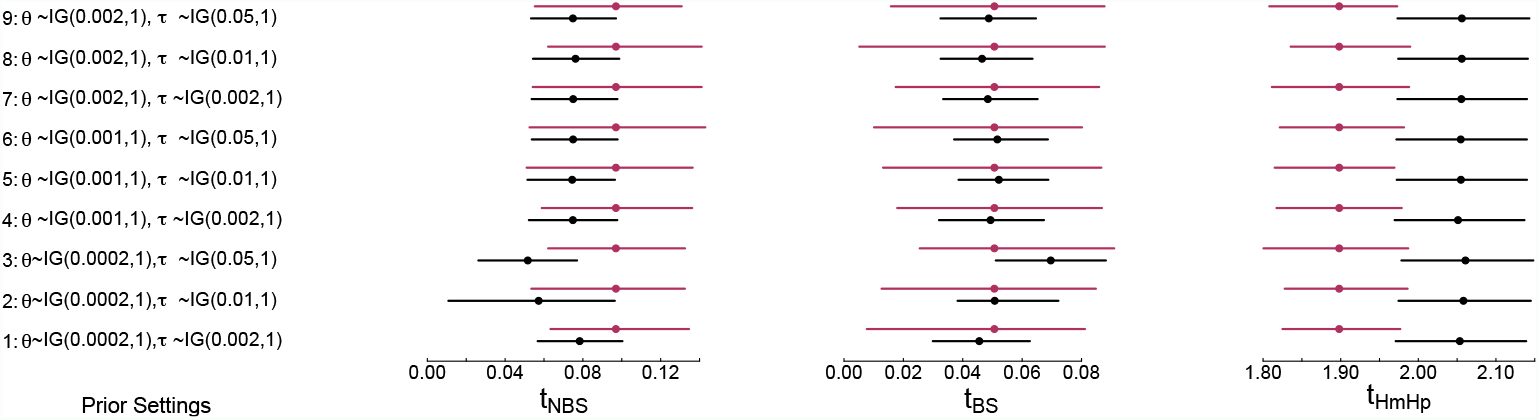
95% credible (*BPP*; black) and confidence (*MAP*_CL_; maroon) intervals for the gibbon data for the 9 prior choices considered here (left panel). Setting 5 (*θ*∼*IG*(0.001, 1.0) and *τ*_R_∼*IG*(0.01, 1.0)) is the closest match to the priors used by Shi and Yang (2018).

Figure 9 shows the error distribution resulting from our simulations to check the robustness of the *MAP*_CL_ estimator to variable *θ* parameters. It is evident from both plots that estimation accuracy does not change significantly when data are generated using different combinations of *θ*s. In the worst case, the estimates for the smallest speciation time between species S and B deviate from the truth by 15% on average. As this branch is quite short (0.000885), the RMSE of 0.00014 indicates that the *MAP*_CL_ estimator performs well even for this case.

**Fig. 9.**
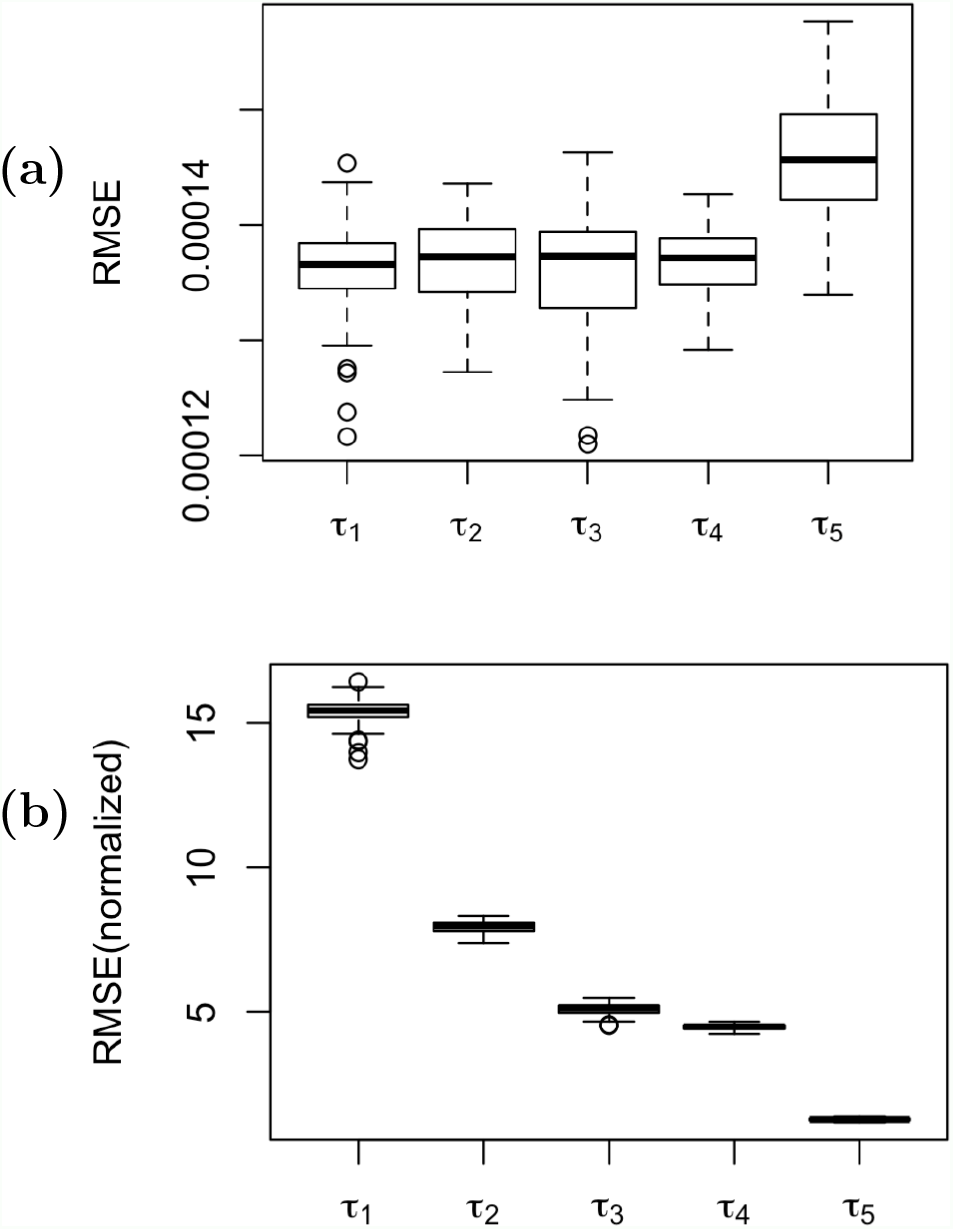
Boxplots of RMSEs for node age estimates (in mutation units) for the gibbon tree with varying *θ*s. Each box in the plot represents the distribution of (a) RMSEs or (b) normalized RMSEs for 100 combinations of *θ* values. Node ages (*τ* s) correspond to Figure 2(b).

## 4 Discussion

### 4.1 Is our method Bayesian or frequentist?

An earlier version of our method used a simpler composite likelihood estimator, maximizing log *CL*(***τ***, *θ*; ***x***) in (2) rather than incorporating prior terms as in (4). We switched to MAP estimation as a means of dealing with instability in the optimization due to the flatness of the likelihood surface in regions of parameter space that have low likelihood. The incorporation of vague prior terms increases the gradient of the optimality surface in these regions, mitigating problems with premature termination (due to floating-point roundoff error) and/or overshooting.

A reviewer suggested that the change from a “frequentist” to a “Bayesian” perspective constituted a fundamental change in methodology requiring additional theoretical validation. Although we have addressed technical aspects of this change in the Supplemental Material, the distinction between these perspectives is not as stark as it might seem. The technique of regularization—intentionally increasing the bias of an estimator in order to decrease the overall estimation error—is commonly used by frequentists. For example, “penalty” or “shrinkage” terms are added to the loss function in the lasso and ridge regression methods (Tibshirani, 1996), but these methods can be equivalently treated as Bayesian MAP estimates where a specific prior distribution corresponds to the regularization term (e.g., Tibshirani, 1996, section 5). Because we use confidence intervals rather than Bayesian credible intervals to assess the uncertainty of our estimates, we accept that our method is well characterized as a frequentist penalized likelihood method. However, because of the Bayesian motivation for our penalty term, it is also formally a Bayesian method as well, while differing from more typical fully Bayesian methods in (1) using a composite posterior density rather than a posterior density that incorporates the true likelihood, and (2) obtaining point estimates from a mode, rather than the mean, of the posterior distribution). Consequently, the choice between classifying it as frequentist or a Bayesian method is largely a matter of personal taste; the result is the same regardless of which perspective is adopted. Of course, a standard composite likelihood method is obtained if uniform priors are used for *τ*_*R*_ and *θ*, and although not recommended, this option is available in *PAUP\**.

### 4.2 Computational efficiency of the *MAP*_CL_ estimator

To obtain good *MAP*_CL_ estimates of the node ages on a species tree, we need to be able to do two things well: compute the composite likelihood, and search the parameter space for values that optimize the posterior probability density. The former can be done very efficiently for trees of arbitrary size. The number of individual likelihoods for all possible quartets (2) grows as the 4^*th*^ power of the tree size, but the amount of work required per quartet is light, so that the total likelihood can be computed quickly even for a large tree.

Despite running far more quickly than fully Bayesian methods including *BPP* and *StarBEAST2*, the second task becomes more difficult as the dimension of the parameter space increases. Fortunately, the gradient (first partial derivatives of the posterior density function with respect to each parameter) can be calculated quickly for any point in the parameter space, allowing the use of quasi-Newton optimizers that typically need fewer function evaluations to converge to an optimum than derivative-free methods. In addition, the bootstrapping procedure used for measuring uncertainty could easily be parallelized, although we have not yet done so.

### 4.3 Assumptions and performance of the *MAP*_CL_ estimator

The assumptions that (1) nucleotide sites evolve according to the JC69 substitution model, and (2) effective population sizes are constant throughout the tree, permit the use of formulas in Chifman and Kubatko (2015) for computing the site pattern probabilities used in equation (1). Without these closed-form expressions, exact calculation of site pattern probabilities would involve an intractable multidimensional integration over gene trees and their associated branch lengths. Empirical data, however, may evolve under a nucleotide substitution model more complex than JC69, and preliminary simulations indicate that our method is not always robust when the nucleotide substitution model is misspecified and divergence between species is high (results not shown). However, for closely related species like gibbons, Shi and Yang (2018) argue that the JC69 model should be adequate for *BPP* and *ASTRAL*, and we note that *BPP* currently also assumes the JC69 model. Our simulations do indicate that our method is robust to the violation of the assumption of constant effective population sizes across all populations, although preliminary simulations indicate that in some cases, the combined impact of a misspecified substitution model and constant *θ* parameter can be substantial.

We are investigating plausible approaches for extending our method to allow inference under more general models, such as the GTR model and its submodels. An obvious, but computationally expensive, strategy would be to estimate site pattern probabilities by Monte Carlo simulation of a large number of independent sites under the assumed model for each point in parameter space visited by the optimizer. We are exploring an alternative method that makes a deterministic estimate of the desired vector of site pattern probabilities using the expected lengths of the branches on each possible gene tree, conditional on a species-tree topology and the current set of *θ* and ***τ*** values. The latest versions of *PAUP\** support this method (to be described in a subsequent paper), allowing the choice between using exact site-pattern probabilities under the JC model, or an approximation of these probabilities under more complex models.

In summary, our *MAP*_CL_ estimator of speciation times has several qualities that set it apart from existing methods. It is fast enough to be used for data sets that are too large for existing fully Bayesian methods. It does not appear to be overly sensitive to the location of weakly informative inverse-gamma priors (*BPP* can exhibit slow convergence when the locations of prior and posterior distributions are very different). Even if the ultimate goal is to conduct *BPP* or *StarBEAST2* analyses, our method may be useful in parameterizing prior distributions in an empirical Bayes setting, or in choosing starting values for MCMC iterations in a fully Bayesian analysis. Unlike other fast methods, including *ASTRAL* and *MP-EST*, it does not require prior estimation of gene trees. It is both statistically consistent and asymptotically normal, ensuring good statistical properties as the amount of data increases. It can handle both CIS and multilocus data and can accommodate the sampling of multiple individuals per species. We anticipate that it will be a useful addition to the collection of methods available for inferring speciation times from genome-scale data under the multispecies coalescent model.

## Supporting information

Implementation of qAge in PAUP*

## Acknowledgments

We thank Jeff Thorne for helpful discussion regarding statistical issues, as well as the three anonymous reviewers who made important suggestions that strengthened the quality of the paper. We also acknowledge University of Florida Research Computing (http://rc.ufl.edu) and The Ohio State University College of Arts and Sciences (http://go.osu.edu/unitycompute) for providing computational resources.

