## Supplementary material for "Estimation of Speciation Times Under the Multispecies Coalescent": Implementation of qAge in PAUP*

David L. Swofford

16 December 2020

PAUP* now estimates divergence times for a species tree assuming a multispecies coalescent process, using the quartet-based method of Peng *et al.* (2021) under the Jukes-Cantor substitution model. This method, called **qAge**, relies on functions for computing expected site pattern probabilities given in Supplement A to Chifman and Kubatko (2015). Peng *et al.* (2021) provide mathematical and statistical background; this document supplies additional mathematical and algorithmic details for the code in PAUP* that may be useful to others interested in understanding or implementing the method.

Numerical optimization of equation (3) in Peng *et al.* (2021) is a nonlinear optimization problem that must satisfy constraints that the $\boldsymbol{\tau}$ and $\theta$ parameters be nonnegative, and that the age of a node cannot exceed that of its parent. Although "active-set" methods using Lagrange multipliers are available for solving these problems (e.g., Nocedal and Wright, 2006), this approach is mathematically and algorithmically complex and requires an artificial increase in the dimensionality of the problem. Instead, the PAUP* implementation uses "black-box" optimization routines and enforces the required constraints through variable transformations.

(*Note:* The following initials are used to refer to individuals mentioned in this document: JC = Julia Chifman; LK = Laura Kubatko; JP = Jing Peng; DLS = David Swofford)

### 1 Parameterization of node ages

The species tree is a rooted, binary tree with $S$ tip species. Internal nodes are indexed by the integers 1 through $S-1$ in the order nodes are visited in a postorder traversal, with node ages $\tau_j$ for each node $j$. In the remainder of this document, the term "node" will be assumed to refer to an internal node unless otherwise indicated. The root of the tree has index $R = S-1$; otherwise we denote the index of a node $j$'s parent by $P(j)$. The free parameters being optimized are $\tau_1, \tau_2, \ldots, \tau_R$ as well as (optionally) the $\theta$ parameter. In order to recast the inequality constraints $\tau_j \le \tau_{P(j)}$ to simple bound constraints, we reparameterize node ages using age ratios $\tau_j/\tau_{P(j)}$ for nodes $1, 2, \ldots, R-1$, which are restricted to the interval $[0,1]$. The root age and $\theta$ parameters must both be nonnegative. Entries in the full parameter vector $\boldsymbol{y}$ are then defined as

$$y_j = \begin{cases} \tau_j/\tau_{P(j)}, & \text{if } 1 \le j < R; \\ \tau_j, & \text{if } j = R; \\ \theta, & \text{otherwise } (j = S). \end{cases} \tag{1}$$

For each point $\boldsymbol{y}$ evaluated during the optimality search, we convert the $y_j$ corresponding to node ages back to $\tau_j$ prior to evaluating the likelihood function:

$$\tau_j = \begin{cases} y_j, & \text{if } j = R; \\ y_j \tau_{P(j)}, & \text{if } j < R, \end{cases} \tag{2}$$

where nodes are visited in a preorder traversal of tree so that $\tau_{P(j)}$ will be updated prior to using it for calculation of $\tau_j$.

### 2 Optimization details

The estimator given by Peng *et al.*'s (2021) equation (4) is obtained by minimizing the negative log posterior density function

$$G(\boldsymbol{\tau}, \theta) = -\left(\log f_\theta(\theta) + \log f_\text{h}(\tau_R) + \sum_{i=1}^{Q} \ell_i(\boldsymbol{\tau}, \theta | \boldsymbol{v}_i)\right), \tag{3}$$

where $f_\theta(\theta)$ is the prior density of $\theta$, $f_\text{h}(\tau_R)$ is the prior density of the tree height (= root age) $\tau_R$, $Q$ is the number of possible quartets $= \binom{S}{4}$, and $\ell_i$ is a function that returns $\log L_i$, with $L_i$ equal to the likelihood of the $\theta$ and $\boldsymbol{\tau}$ parameters for quartet $i$. ($L_i$ is the probability of observing the vector of site pattern occurrences $\boldsymbol{v}_i$ for this quartet given the values of $\theta$ and $\boldsymbol{\tau}$; see Peng *et al.*, 2021, for details.) The $\log(f_\theta)$ and $\log(f_\text{h})$ functions are of the form (4), but use different shape and scale parameters for the inverse gamma distribution. Note that $G(\boldsymbol{\tau}, \theta)$ can be used for composite likelihood rather than MAP estimation simply by omitting the two log prior terms (equivalent to assuming uniform priors on $\theta$ and $\boldsymbol{\tau}$).

We perform a multidimensional optimization simultaneously for all parameters. PAUP* supports two optimization algorithms: (1) the "Praxis" refinement of Powell's (1970) method of conjugate search directions (Brent, 1973), and (2) the quasi-Newton limited-memory BFGS method (L-BFGS-B version 3.0; Byrd *et al.*, 1995; Zhu *et al.*, 1997; Morales and Nocedal, 2011). The Praxis implementation in PAUP* is a hand translation of the Algol code in Brent (1973) to C by DLS. For L-BFGS-B, the Fortran code of Zhu *et al.* (1997) is called from a wrapper written in C.

Both Praxis and L-BFGS typically converge to the same solution, but L-BFGS is almost always faster, and all analyses reported in Peng *et al.* (2021) were performed using this method. Note that the L-BFGS-B code (Morales and Nocedal, 2011) allows enforcement of simple lower and/or upper bound constraints on the variables. However, we have found that convergence is both more reliable and faster using variable transformations (see section 2.2.1) that permit unconstrained optimization.

#### 2.1 Optimization of the posterior density function

To optimize model parameters using L-BFGS, we need to compute the gradient of $G(\boldsymbol{\tau}, \theta)$ evaluated at a point $\{\tau_1, \tau_2, \ldots, \tau_R, \theta\}$. For a tree with exactly four tip species, we can write the likelihood function corresponding to equations 1 and 2 of Peng *et al.* (2021) as $L_i\left(\tau_1, \tau_2, \tau_3, \theta | \boldsymbol{v}_i\right))$, where $\tau_1$, $\tau_2$, and $\tau_3$ are the ages of the three internal nodes. LK, JC, and DLS developed formulas for computing the partial derivatives of $\ell$ with respect to the $\boldsymbol{\tau}$ and $\theta$ parameters: $\partial\ell_i/\partial\tau_1$, $\partial\ell_i/\partial\tau_2$, $\partial\ell_i/\partial\tau_3$, and $\partial\ell_i/\partial\theta$ (documented elsewhere).

Extending these derivatives to the gradient used for optimizing $G(\boldsymbol{\tau}, \theta)$ in (3) requires dealing with four additional issues: (1) incorporation of the prior terms $f_\theta(\theta)$ and $f_\text{h}(\tau_R)$, (2) parameterization of node ages as proportions in order to enforce the constraint that node $j$ cannot be older than its parent $P(j)$; (3) calculation of partial derivatives of $G(\boldsymbol{\tau}, \theta)$ with respect to each parameter when the species tree contains more than four tips; and (4) handling of the arcsine square root and log transformations used to eliminate lower and upper bounds, allowing unconstrained optimization of all parameters.

##### 2.1.1 Log prior densities

Following Flouri *et al.* (2018), we use the inverse gamma distribution as a prior on the root age and $\theta$ parameters, which has probability density equal to

$$IG(x; \alpha, \beta) = \frac{\beta^\alpha}{\Gamma(\alpha)} x^{-\alpha-1} \exp\left(-\frac{\beta}{x}\right),$$

where $\alpha$ and $\beta$ are the shape and scale parameters, respectively. To evaluate the log prior densities in (3), we use

$$\log f(x) = \log IG(x; \alpha, \beta) = \alpha \log(\beta) - \log(\Gamma(\alpha)) - (\alpha+1)\log(x) - \frac{\beta}{x}, \tag{4}$$

with $\log \Gamma(\alpha)$ evaluated using the C library function *lgamma*. The partial derivative of (4) with respect to $x \in \{\theta, \tau_R\}$, needed for computing the gradient of the log posterior density function, is given by

$$\frac{\partial}{\partial x} \log f(x) = \frac{\beta}{x^2} - \frac{(\alpha+1)}{x}. \tag{5}$$

In PAUP*, parameters of the inverse gamma distribution are specified using the mean $\mu$ and coefficient of variation $c_\text{v} = \sigma/\mu$ rather than $\alpha$ and $\beta$, where

$$\mu = \frac{\beta}{\alpha - 1}, \qquad \sigma = \frac{1}{\sqrt{\alpha - 2}}, \qquad \alpha > 2.$$

Before evaluating (4), $\mu$ and $c_\text{v}$ are converted to $\alpha$ and $\beta$ using

$$\alpha = c_\text{v}^{-2} + 2, \qquad \beta = \mu\left(c_\text{v}^{-2} + 1\right).$$

##### 2.1.2 Likelihood function and gradient evaluation for individual quartets

The $i$th quartet (containing four tip species) induces a subtree $\mathcal{T}_i$ of the full species tree; we can compute partial derivatives for nodes of the full tree by combining terms from individual quartets. The induced subtree, which can be either symmetric or asymmetric, contains three internal nodes from the full tree. We designate these three nodes $u_i$, $v_i$, and $w_i$, again in order of a postorder traversal.

Define the functions $h_1, h_2, \ldots, h_Q$ in terms of $\boldsymbol{y}$ as

$$h_i(y_{u_i}, y_{v_i}, y_{w_i}, \theta) = \ell_i(\tau_{u_i}, \tau_{v_i}, \tau_{w_i}, \theta), \tag{6}$$

where $y_{u_i}$, $y_{v_i}$, and $y_{w_i}$ are the reparameterized node ages corresponding to $\tau_{u_i}$, $\tau_{v_i}$, and $\tau_{w_i}$ from (1) and $\ell_i$ is the log likelihood function for quartet $i$ as in (3). The $h_i$ functions in (6) are never evaluated directly; instead, the $y_i$ are backtransformed to $\tau_i$ parameters using (2) prior to evaluation of $\ell_i$. However, for L-BFGS-B optimization, we need the gradient of $h_i$ evaluated at $\boldsymbol{y}$:

$$\nabla h_i = \begin{bmatrix} \dfrac{\partial h_i}{\partial y_{u_i}} \\ \dfrac{\partial h_i}{\partial y_{v_i}} \\ \dfrac{\partial h_i}{\partial y_{w_i}} \\ \dfrac{\partial h_i}{\partial \theta} \end{bmatrix}. \tag{7}$$

Applying the chain rule for $j \in \{u_i, v_i, w_i\}$,

$$\frac{\partial h_i}{\partial y_j} = \frac{\partial h_i}{\partial \tau_{u_i}} \frac{\partial \tau_{u_i}}{\partial y_j} + \frac{\partial h_i}{\partial \tau_{v_i}} \frac{\partial \tau_{v_i}}{\partial y_j} + \frac{\partial h_i}{\partial \tau_{w_i}} \frac{\partial \tau_{w_i}}{\partial y_j}. \tag{8}$$

Form an ordered set $\mathcal{A}$ containing the indices of the $m$ nodes that lie in the path from a node $c$ to the root node $R$ (inclusive):

$$\mathcal{A} = (c{=}a_1,\ a_2,\ \ldots,\ a_{m-1},\ a_m{=}R)$$

By sequential application of (1), we can express the age of any node $c$ as

$$\tau_c = \prod_{j=1}^{m} y_{a_j}. \tag{9}$$

If node $d = a_k$ is an ancestor of node $c$ at location $k$ in $\mathcal{A}$, then we can write (9) as

$$\tau_c = \left(\prod_{j=1}^{k-1} y_{a_j}\right) y_d \left(\prod_{j=k+1}^{m} y_{a_j}\right),$$

with empty products set to 1, so that for any $c$ and $d$, we obtain the partial derivatives needed by (8) as

$$\frac{\partial \tau_c}{\partial y_d} = \begin{cases} \dfrac{1}{y_d} \displaystyle\prod_{j=1}^{m} y_{a_j}, & \text{if } d \in \mathcal{A}; \\ 0, & \text{otherwise.} \end{cases}$$

The final $\nabla h_i$ entry in (7) is simply

$$\frac{\partial h_i}{\partial \theta} = \frac{\partial \ell_i}{\partial \theta}. \tag{10}$$

#### 2.1.3 Posterior density and gradient evaluation for full tree

Extending the gradient formulas for a single quartet described above to the full tree is straightforward. Since $y_R = \tau_R$ and $y_S = \theta$, we can rewrite (3) as

$$H(\boldsymbol{y}) = G(\boldsymbol{\tau}, \theta) = -\big(\log f_\theta(y_S) + \log f_\mathrm{h}(y_R) + \mathcal{L}(\boldsymbol{y})\big)$$

where $\mathcal{L}(\boldsymbol{y})$ corresponds to the log composite likelihood term, but in terms of the reparameterized variables $\boldsymbol{y}$ rather than $\boldsymbol{\tau}$ and $\theta$:

$$\mathcal{L}(\boldsymbol{y}) = \sum_{i=1}^{Q} h_i(\boldsymbol{y}). \tag{11}$$

Hence, we need $\nabla H(\boldsymbol{y})$ corresponding to $\nabla G(\boldsymbol{\tau}, \theta)$:

$$\nabla G = \begin{bmatrix} \frac{\partial G}{\partial \tau_1} \\ \frac{\partial G}{\partial \tau_2} \\ \vdots \\ \frac{\partial G}{\partial \tau_R} \\ \frac{\partial G}{\partial \theta} \end{bmatrix}, \qquad \nabla H = \begin{bmatrix} \frac{\partial H}{\partial y_1} \\ \frac{\partial H}{\partial y_2} \\ \vdots \\ \frac{\partial H}{\partial y_R} \\ \frac{\partial H}{\partial y_S} \end{bmatrix}. \tag{12}$$

Using $\partial h_i/\partial y_j$ from (8), partial derivatives of $\mathcal{L}$ with respect to node ages are obtained as

$$\frac{\partial \mathcal{L}}{\partial y_j} = \sum_{i=1}^{Q} \frac{\partial h_i}{\partial y_j}, \qquad 1 \le j \le R, \tag{13}$$

where $\partial h_i/\partial y_j = 0$ if $j \notin \{u_i, v_i, w_i\}$. Likewise, partial derivatives of $\mathcal{L}$ with respect to the $\theta$ parameter are obtained using (10):

$$\frac{\partial \mathcal{L}}{\partial y_S} = \frac{\partial \mathcal{L}}{\partial \theta} = \sum_{i=1}^{Q} \frac{\partial h_i}{\partial \theta}. \tag{14}$$

Incorporating the prior terms in (3), and using (5) and (13-14), we fill the gradient vector

$$(\nabla H)_j = \begin{cases} -\frac{\partial \mathcal{L}}{\partial y_j}, & \text{for } 1 \le j < R; \\ -\frac{\partial \mathcal{L}}{\partial \tau_R} - \frac{\partial}{\partial y} \log f_{\mathrm{h}}(\tau_R), & \text{for } j = R; \\ -\frac{\partial \mathcal{L}}{\partial \theta} - \frac{\partial}{\partial y} \log f_{\theta}(\theta), & \text{for } j = S. \end{cases} \tag{15}$$

The derivatives in (15) are negated because we are minimizing the negative posterior density rather than maximizing the posterior density.

We can now perform gradient-based optimization of the function $H(\boldsymbol{y})$. The $MAP_{\mathrm{CL}}$ estimator for node ages $\boldsymbol{\tau}$ and $\theta$ is obtained by backtransforming

$$\tilde{\boldsymbol{y}} = \underset{\boldsymbol{y}}{\operatorname{argmin}}\, H(\boldsymbol{y})$$

using (2). The optimization routine must allow simple lower and upper bounds on the variables in order to satisfy the constraints:

$$
\begin{aligned}
0 \le y_j \le 1, &\qquad 1 \le j < R\,; \\
y_j > 0, &\qquad j = R \quad (y_j = \tau_R)\,; \\
y_j > 0. &\qquad j = S \quad (y_j = \theta)\,.
\end{aligned}
$$

PAUP* supports the quasi-Newton L-BFGS-B method (Zhu *et al.*, 1997; Morales and Nocedal, 2011), which can enforce these bounds.

### 2.2 Allowing unconstrained optimization

Although the approach described in section 2.1 works reasonably well with L-BFGS-B, we have found that converting the optimization problem to a fully unconstrained one runs much faster, due to fewer iterations being needed for convergence to a final solution. We can eliminate the bound constraints in (1) using suitable transformations (Box, 1966; Sisser, 1981). These transformations are mandatory when using the Praxis optimization method, which does not support bound constraints.

#### 2.2.1 Transformations

Arcsine square root transformations are used for age ratios; log transformations are used to enforce nonnegativity of the root age (= $\tau_R$) and $\theta$ parameters. After these transformations, the parameter vector $\boldsymbol{z}$ becomes

$$
z_j = \begin{cases} \sin^{-1}\left(\sqrt{\tau_j/\tau_{P(j)}}\right), & \text{if } 1 \le j < R; \\ \log \tau_j, & \text{if } j = R; \\ \log \theta, & \text{otherwise } (j = S). \end{cases} \tag{16}
$$

In order to evaluate the log likelihood function $\ell_i$, the current parameter values $\boldsymbol{z}$ are first backtransformed using

$$
\begin{aligned}
\tau_j &= \begin{cases} e^{z_j}, & \text{if } j = R; \\ \tau_{P(j)} \sin^2(z_j), & \text{if } 1 \le j < R; \end{cases} \\
\theta &= e^{z_S}.
\end{aligned} \tag{17}
$$

where nodes indexed by $j$ are visited in preorder so that $\tau_{P(j)}$ will be updated prior to using it for calculation of $\tau_j$.

#### 2.2.2 Likelihood function and gradient evaluation for individual quartets

Analogously to (6), define the functions $h'_1, h'_2, \ldots, h'_Q$ in terms of $\boldsymbol{z}$ as

$$
h'_i(z_{u_i}, z_{v_i}, z_{w_i}, z_S) = \ell_i(\tau_{u_i}, \tau_{v_i}, \tau_{w_i}, \theta),
$$

where $z_{u_i}$, $z_{v_i}$, and $z_{w_i}$ are the transformed node ages corresponding to $\tau_{u_i}$, $\tau_{v_i}$, and $\tau_{w_i}$ from (16), and $z_S = \log\theta$. The $h'_i$ functions are backtransformed to $\tau_i$ parameters using (17) prior to evaluation of $\ell_i$. For optimization using quasi-Newton methods, we need the gradient of $h'_i$ evaluated at $\boldsymbol{z}$:

$$\nabla h'_i = \begin{bmatrix} \dfrac{\partial h'_i}{\partial z_{u_i}} \\ \dfrac{\partial h'_i}{\partial z_{v_i}} \\ \dfrac{\partial h'_i}{\partial z_{w_i}} \\ \dfrac{\partial h'_i}{\partial z_S} \end{bmatrix} \tag{18}$$

As in (8), we obtain $h'_i$ using the chain rule for $j \in \{u_i, v_i, w_i\}$:

$$\frac{\partial h'_i}{\partial z_j} = \frac{\partial h'_i}{\partial \tau_{u_i}} \frac{\partial \tau_{u_i}}{\partial z_j} + \frac{\partial h'_i}{\partial \tau_{v_i}} \frac{\partial \tau_{v_i}}{\partial z_j} + \frac{\partial h'_i}{\partial \tau_{w_i}} \frac{\partial \tau_{w_i}}{\partial z_j}. \tag{19}$$

However, calculation of $\partial \tau_{u_i}/\partial z_j$, $\partial \tau_{v_i}/\partial z_j$, and $\partial \tau_{w_i}/\partial z_j$ is more complicated due to the transformations. First, backtransform all $z_j$ to obtain $\tau_j$ and $\theta$ using (17).

For any pair of nodes $(c, d)$ such that either $c = d$ or node $c$ is ancestral to node $d$, form an ordered set $\mathcal{B}$ containing the indices of the $m$ nodes that lie in the path starting at node $c$ and ending at node $d$ (inclusive):

$$\mathcal{B} = (c{=}b_1,\, b_2,\, \ldots,\, b_{m-1},\, b_m{=}d).$$

At the root node ($c = R$),

$$\frac{\partial \tau_c}{\partial z_c} = \frac{\partial}{\partial z_R} e^{z_R} = e^{z_R} = \tau_R\,;$$

otherwise ($c < R$),

$$\frac{\partial \tau_c}{\partial z_c} = \frac{\partial}{\partial z_R} \tau_{P(c)} \sin^2(z_c) = 2\, \tau_{P(c)} \sin(z_c) \cos(z_c).$$

Then for descendant nodes $d$ of node $c$:

$$\frac{\partial \tau_c}{\partial z_d} = \frac{\partial \tau_c}{\partial z_c} \prod_{j=2}^{m} \sin^2\!\left(z_{b_j}\right).$$

The final $\nabla h'_i$ entry in (18) is

$$\begin{aligned} \frac{\partial h'_i}{\partial z_S} &= \frac{\partial h'_i}{\partial \theta} \frac{\partial \theta}{\partial z_S} = \frac{\partial h'_i}{\partial \theta} \frac{\partial e^{z_S}}{\partial z_S} = \frac{\partial h'_i}{\partial \theta} e^{z_S} \\ &= \frac{\partial h'_i}{\partial \theta}\, \theta. \end{aligned} \tag{20}$$

#### 2.2.3 Posterior density and gradient evaluation for full tree

The formulas for extending gradients for a single quartet to the full tree are similar to the untransformed case (section 2.1.3). Since $z_R = \log \tau_R$ and $z_S = \log \theta$, we can rewrite (3) as

$$H'(\boldsymbol{z}) = G(\boldsymbol{\tau}, \theta) = -\big(\log f_\theta(e^{z_S}) + \log f_{\mathrm{h}}(e^{z_R}) + \mathcal{L}'(\boldsymbol{z})\big)$$

where $\mathcal{L}'(\boldsymbol{z})$ corresponds to the log composite likelihood term, but now in terms of the reparameterized variables $\boldsymbol{z}$ rather than $\boldsymbol{\tau}$ and $\theta$:

$$\mathcal{L}'(\boldsymbol{z}) = \sum_{i=1}^{Q} h_i'(\boldsymbol{z}). \tag{21}$$

Hence, we need $\nabla H'(\boldsymbol{z})$ corresponding to $\nabla G(\boldsymbol{\tau}, \theta)$ in (12):

$$\nabla H' = \begin{bmatrix} \frac{\partial H'}{\partial z_1} \\ \frac{\partial H'}{\partial z_2} \\ \vdots \\ \frac{\partial H'}{\partial z_R} \\ \frac{\partial H'}{\partial z_S} \end{bmatrix}.$$

Using $\partial h_i'/\partial z_j$ from (19), partial derivatives of $\mathcal{L}'$ with respect to node ages are obtained as

$$\frac{\partial \mathcal{L}'}{\partial z_j} = \sum_{i=1}^{Q} \frac{\partial h_i'}{\partial z_j}, \qquad 1 \leq j \leq R, \tag{22}$$

where $\partial h_i'/\partial z_j = 0$ if $j \notin \{u_i, v_i, w_i\}$. Partial derivatives of $\mathcal{L}'$ with respect to the transformed $\theta$ parameter are obtained using (20):

$$\frac{\partial \mathcal{L}'}{\partial z_S} = \sum_{i=1}^{Q} \frac{\partial h_i'}{\partial \theta}.$$

The partial derivative of (4) with respect to transformed $\theta$ and $\tau_R$ parameters $z = \log x$ for $x \in \{\theta, \tau_R\}$, needed for computing the gradient of the log posterior density function, is given by

$$\begin{aligned} \frac{\partial}{\partial z}\log f(e^z) &= \frac{\partial}{\partial x}\log f(x)\,\frac{\partial x}{\partial z} = \frac{\partial}{\partial x}\log f(x)\,\frac{\partial e^z}{\partial z} = \left(\frac{\partial}{\partial x}\log f(x)\right) e^z \\ &= x\,\frac{\partial}{\partial x}\log f(x) \end{aligned} \tag{23}$$

Incorporating these values for the prior terms in (3) and using (22-23), we fill the gradient vector

$$(\nabla H')_j = \begin{cases} -\dfrac{\partial \mathcal{L}'}{\partial z_j}, & \text{for } 1 \leq j < R; \\ -\dfrac{\partial \mathcal{L}'}{\partial z_j} - \tau_R \dfrac{\partial}{\partial z}\log f_{\mathrm{h}}(\tau_R), & \text{for } j = R; \\ -\dfrac{\partial \mathcal{L}'}{\partial z_j} - \theta \dfrac{\partial}{\partial z}\log f_{\theta}(\theta), & \text{for } j = S. \end{cases}$$

As in (15), these derivatives are negated because the optimization minimizes the negative posterior density.

We can now perform *unconstrained* gradient-based optimization of the function $H'(\boldsymbol{z})$. The $MAP_{\mathrm{CL}}$ estimator for node ages $\boldsymbol{\tau}$ and $\theta$ is obtained by backtransforming

$$\tilde{\boldsymbol{z}} = \underset{\boldsymbol{z}}{\operatorname{argmin}}\ H'(\boldsymbol{z})$$

using (17).

### 3 Obtaining a starting point for the optimization

It is helpful (and sometimes essential) to begin the optimization of $H'(\boldsymbol{z})$ (or $H(\boldsymbol{y})$) with reasonable starting values for the node ages and $\theta$.

#### 3.1 Estimation of node ages conditional on $\boldsymbol{\theta}$

For any four-tip species tree and a given value of $\theta$, we can use the method-of-moments estimator of Kubatko and Chifman (2020) to obtain estimates $\hat{\tau}_1$, $\hat{\tau}_2$, and $\hat{\tau}_3$ for the ages of the three internal nodes. Although Kubatko and Chifman's estimator is limited to four species, we can extend it to $S > 4$ species by averaging $\hat{\tau}_j$ over all quartets that include node $j$ in their induced subtrees. As in section 2.1.2, we designate the three internal nodes of the subtree induced by quartet $i$ as $u_i$, $v_i$, and $w_i$ in order of a postorder traversal.

The mean age estimate for any node $j$ of the full tree over all quartets is then

$$\tau_j^* = \overline{\hat{\tau}_j} = \frac{\sum_i \{\hat{\tau}_{u_i}(\mathbb{1}_{j=u_i}) + \hat{\tau}_{v_i}(\mathbb{1}_{j=v_i}) + \hat{\tau}_{w_i}(\mathbb{1}_{j=w_i})\}}{\sum_i \mathbb{1}_{j\in\{u_i,v_i,w_i\}}}$$

where $\mathbb{1}_{condition}$ is an indicator function equal to 1 when *condition* is true and 0 otherwise, with summations taken over all possible quartets (the denominator is just the number of quartets for which node $j$ is included in the subtree induced by the $i$th quartet).

Note that the Kubatko-Chifman estimator can return negative $\tau$ values for one or more nodes with increasing probability as $\theta$ increases in magnitude; we say that the $\theta$ value is *infeasible* when this occurs for any quartet.

#### 3.2 Starting values for joint optimization of node ages and $\boldsymbol{\theta}$

Using good starting values for $\boldsymbol{\tau}$ and $\theta$ substantially reduces the number of function evaluations needed for multidimensional parameter optimization. We obtain a starting value for $\theta$ by optimizing a posterior-density (or likelihood) function corresponding to (3):

$$g(\theta) = -\left(\log f_\theta(\theta) + \log f_{\mathrm{h}}(\tau_R^*) + \sum_{i=1}^{Q} \ell_i(\boldsymbol{\tau}^*, \theta|\boldsymbol{v}_i)\right),$$

but where for any given value of $\theta$, the node ages $\boldsymbol{\tau}^*$ are computed using the (very fast) method described in section 3.1. Minimization of $g(\theta)$ is performed using a one-dimensional, derivative-free method, with the initial $\theta$ value chosen as

$$\theta_0 = \underset{\theta}{\text{argmin}}\ g(\theta).$$

Unfortunately, $g(\theta)$ is somewhat difficult to optimize. Due to correlation between $\theta$ and $\boldsymbol{\tau}^*$, the density is relatively, if not extremely, flat for values of $\theta$ less than the optimal value, sometimes causing a black-box optimization routine to terminate prematurely. The function is much steeper on the other side, however, suggesting an *ad hoc* method for finding a bracketing interval $[a, b]$ that contains $\theta_0$. We start by choosing a tiny lower bound $a$ (currently $10^{-5}$) assumed to be less than $\theta_0$. The upper bound $b$ is then iteratively increased by a factor of 10, starting with $b{=}10a$, until $\theta{=}b$ becomes infeasible, yielding an initial interval $[a, b]$ with $a \leq \theta_0 \leq b$.

Obviously, this bound is very loose, and we only use it to initiate a golden section search (e.g., Gill *et al.*, 1981) to move $b$ into the feasible region for the Kubatko-Chifman estimator and further tighten the bracket. The advantage of golden section search is that we do not need to compute the posterior density $g(\theta)$ when $\theta{=}b$ is outside this feasible region, which is undefined due to the negative node-age estimates. Instead, we can just set $g(\theta) = \infty$, ensuring that the $[a, b]$ interval will be shrunk by reducing $b$ rather than increasing $a$.

The *disadvantage* of golden section search is that it converges slowly, so we use a large stopping tolerance, terminating the search once $b - a < 0.01$. Starting with this $[a, b]$ bracket, the optimization is then completed using the *localmin* procedure of Brent (1973), which achieves superlinear convergence via a combination of golden section and parabolic interpolation steps.

After $\theta_0$ has been determined, we set the starting node ages $\boldsymbol{\tau}_0$ to the values of $\tau_j^*$ obtained using the method of section 3.1, with $\theta = \theta_0$.

A minor complication is that the *ad hoc* method for loose bracketing of $\theta_0$ before starting the golden section search ignores the priors on $\theta$ and $\tau_r$. Consequently, it is theoretically possible for the resulting bracket to exclude the optimal $\theta$ value for $G(\boldsymbol{\tau}, \theta)$ when a highly informative prior on $\theta$ is used that is centered far away from $\theta_0$. Because setting a tight prior on $\theta$ would be a questionable decision by the user, we don't worry about this possibility. In any case the only consequence would be slower convergence of the full multidimensional optimization due to a poor starting value for $\theta$.

### 4 Handling multiple individuals sampled per species

Up to this point, we have assumed that the data consist of a single sampled individual for each species. Multiple indiduals per species are also accommodated by including all quartets containing one individual from each of four different species in the set of possible quartets used to evaluate the composite likelihood terms in (11) and (21).

### 5 Validation

All derivations of partial derivatives in sections 2.1 and 2.2 were verified using numerical differentiation by finite differences. Although much slower to compute, the numerically approximated derivatives were very close to the corresponding analytical calculations.

Correctness of the overall implementation is supported by convergence of estimated parameter values to the true values used for simulations as the number of loci and sites/locus becomes large.
