## Supplemental Information for "Estimation of Speciation Times Under the Multispecies Coalescent"

### S1. Statistical consistency and asymptotic normality of the $\boldsymbol{MAP}_{\mathbf{CL}}$ estimator

Here we provide the details of our claim that the $MAP_{\text{CL}}$ estimator is statistically consistent and asymptotically normal. This is done in two steps. In Section S1.1, we prove that the maximum of the composite likelihood is a statistically consistent and asymptotically normal estimator by applying a result from Arnold and Strauss (1991), and we provide an expression for the asymptotic variance. In Section S1.2, we prove that the distribution of the posterior composite likelihood concentrates at the true parameter value, which implies consistency of the $MAP_{\text{CL}}$ estimator. We argue that the $MAP_{\text{CL}}$ estimator shares the same asymptotic distribution as the maximum composite likelihood estimator, though we note that our recommended estimators do not depend on this claim.

#### S1.1 Asymptotic distribution of the maximum composite likelihood estimator, $\boldsymbol{\delta_{MCLE}}$

Consider the composite likelihood given in equation (2) of the main text, where $Q$ is the set of quartets, $\boldsymbol{\delta} = (\boldsymbol{\tau}, \theta)$ is the parameter vector, and $L_i(\cdot)$ is the individual likelihood for the $i^{th}$ quartet. Denote the true parameter values as $\boldsymbol{\delta}_0$, and define

$$\boldsymbol{\delta_{MCLE}} = \underset{\boldsymbol{\delta}}{\operatorname{argmax}} \sum_{i \in Q} \log L_i(\boldsymbol{\delta}; \boldsymbol{v}_i),$$

i.e., $\boldsymbol{\delta_{MCLE}}$ is the maximum likelihood estimate of $\boldsymbol{\delta}_0$ based on the composite likelihood function in Equation (2) of the main text. In this section, we show that this estimator is statistically consistent and asymptotically normal. We begin by reviewing some results of Arnold and Strauss (1991) concerning estimators based on the composite likelihood.

Arnold and Strauss (1991) consider the case of $r$ independent random variables $\boldsymbol{X} = (X^{(1)}, \cdots, X^{(r)})$, each $p \geq 1$ dimensional, with common joint density $h(\boldsymbol{x}; \boldsymbol{\beta})$ that undergo $t$ transformations:

$$Z_i^{(j)} = g_i(X^{(j)}), \;\; i = 1, 2, \cdots, t; j = 1, 2, \cdots, r.$$

Rather than maximize the likelihood, Arnold and Strauss consider maximizing the following product of conditional densities of $\boldsymbol{Z_i}$'s:

$$\prod_{j=1}^{r}\left[\prod_{l}\prod_{l'} h^{\alpha_{ll'}}(\boldsymbol{z_l^{(j)}}|\boldsymbol{z_{l'}^{(j)}};\boldsymbol{\beta})\right] \tag{1}$$

where it is assumed that $l$ and $l'$ are such that the conditional densities are well defined, and $\alpha_{ll'} > 0$. According to Arnold and Strauss (1991), different choices of $l$ and $l'$ lead to estimators with differing efficiency. If we further define random variables $\boldsymbol{Y_i}$'s, where each observation $\boldsymbol{y_i}$ is of form $\boldsymbol{z_l}$ or $(\boldsymbol{z_l}, \boldsymbol{z_{l'}})$, the ratios of the unconditional densities of the random variables $\boldsymbol{Y_i}$'s can be used to express the objective function (1). After taking the logarithm, they define the pseudolikelihood (= composite likelihood) of the data by:

$$\log CL(\boldsymbol{\delta}, \boldsymbol{\beta}) = \sum_{i} \delta_i \sum_{j=1}^{r} \log(h(\boldsymbol{y_i^{(j)}};\boldsymbol{\beta})), \tag{2}$$

where $\delta_i$ is chosen to ensure that this composite likelihood arises from a product of positive likelihoods and conditional likelihoods. Again, the number of individual likelihoods on the right side of (2) depends on the choice of $l$ and $l'$. Under the above definition, the following theorem about statistical consistency is given by Arnold and Strauss (1991).

**Theorem** (Arnold and Strauss, 1991)**.** *Let $X^{(1)}, \ldots, X^{(r)}$ be independent and identically distributed, and suppose that the $Y_i^{(j)'}s$ are such that their densities $h(\boldsymbol{y_i}; \boldsymbol{\beta})$ are differentiable with respect to $\boldsymbol{\beta}$ for almost every $\boldsymbol{y_i}$. Then with probability tending to 1 as $r \to \infty$, the score equation of the log pseudolikelihood has a root $\tilde{\boldsymbol{\beta}}_{\boldsymbol{r}}$ such that $\tilde{\boldsymbol{\beta}}_{\boldsymbol{r}}$ converges in probability to the true parameter value.*

Next we just need to prove the composite likelihood can be written in the form required by the theorem above, and the consistency of the estimator $\boldsymbol{\delta_{MCLE}}$ can be shown. Specifically, we need to show that our composite likelihood can be defined in the same way as (2). In our case, random variable $X$ represents the site pattern counts from the entire tree, so $\boldsymbol{x^{(1)}}, \ldots, \boldsymbol{x^{(M)}}$ are the observations for each of the $M$ sites. If we define $Z_l$ as:

$$\boldsymbol{Z}_l = \begin{cases} 1 & \text{if } l = 1; \\ \boldsymbol{V}_{l-1} & \text{if } l = 2, \cdots, 6, \end{cases}$$

where $\boldsymbol{V}_{l-1} = (\boldsymbol{V}_{l-1}^{(1)}, \ldots, \boldsymbol{V}_{l-1}^{(M)})$ is the collection of random vectors of site pattern indicators at each site as defined in the previous section and $\alpha_{ll'} = 1$, the objective function (1) can be written as:

$$\prod_{j=1}^{M} \left[ \prod_{l=2}^{6} h(\boldsymbol{z_l^{(j)}} | \boldsymbol{z_1^{(j)}}; \boldsymbol{\beta}) \right] = \prod_{j=1}^{M} \left[ \prod_{k=1}^{5} f_k(\boldsymbol{v_k^{(j)}}; \boldsymbol{\tau}, \theta) \right],$$

where $\boldsymbol{v_k^{(1)}}, \cdots, \boldsymbol{v_k^{(M)}}$ are the observations of $\boldsymbol{V}_{l-1}$ for the $M$ sites from quartet $k$, defined as above. From Chifman and Kubatko (2015), the marginal density of the site pattern frequencies for the 4-leaf subtree (i.e., the $f_k(\cdot)$ terms above) can be expressed as

$$L_i(\boldsymbol{\tau}, \theta; \boldsymbol{u}_i) = \prod_{j=1}^{n_i} p_{ij}(\boldsymbol{\tau}, \theta)^{(u_i)_j},$$

where $p_{ij}$ is the $j^{th}$ entry in the appropriate vector of site pattern probabilities for quartet $i$ (either $\boldsymbol{p}^{\mathbf{S}}$ or $\boldsymbol{p}^{\mathbf{A}}$ for a symmetric or asymmetric quartet, respectively), $(u_i)_j$ are the corresponding observed counts, and $n_i$ is either 9 (symmetric quartet) or 11 (asymmetric quartet) (see Equation (1) in the main text).

If we further define random variable $\boldsymbol{Y_i}$, $i = 1, 2, \ldots, |Q|$, where each observation $\boldsymbol{y_i^{(j)}}$ is of the form $\boldsymbol{v_k^{(j)}}$, then the logarithm of the composite likelihood is of the form

$$\log CL(\boldsymbol{\theta}, \boldsymbol{\tau}; \boldsymbol{x}) = \sum_{i \in Q} \log L_i(\boldsymbol{\theta}, \boldsymbol{\tau}; \boldsymbol{y}_i)$$

i.e., our composite likelihood can be written in the form specified by Arnold and Strauss (1991). Using the theorem above, statistical consistency of the estimator $\boldsymbol{\delta_{MCLE}}$ is shown. Furthermore, the next theorem by Arnold and Strauss (1991) states that the $\boldsymbol{\delta_{MCLE}}$ estimator is asymptotically normal and gives an expression for its asymptotic variance.

**Theorem** (Arnold and Strauss, 1991)**.** *Suppose that $X^{(1)}, \cdots, X^{(r)}$ are independent and identically distributed. Assume that the standard regularity conditions hold. Then any consistent sequence $\tilde{\beta}_r = \beta_r(X^{(1)}, \cdots, X^{(r)})$ of roots of the score equation of the log pseudolikelihood satisfies:*

$$\sqrt{r}(\tilde{\beta}_r - \beta) \to N\left(0, \frac{K(\beta)}{J^2(\beta)}\right)$$

*where*

$$K(\beta)=\sum_{i,i'}\mathrm{E}_{\beta}\left[\left\{\frac{\partial}{\partial\beta}\log h(y_i;\beta)\right\}\left\{\frac{\partial}{\partial\beta}\log h(y_{i'};\beta)\right\}\right]$$

*and*

$$J(\beta)=-\sum_{i}\mathrm{E}_{\beta}\left\{\frac{\partial^2}{\partial\beta^2}\log h(y_i;\beta)\right\}.$$

Applying this result to our $MAP_{\mathrm{CL}}$ estimates of node ages, we have:

$$\sqrt{M}(\tilde{\tau}_k-\tau_k)\rightarrow N\big(0,\Sigma_{k,k}\big),$$

where $\Sigma_{k,k}$ is the cell in the $k^{th}$ column and $k^{th}$ row of the variance-covariance matrix $\Sigma$:

$$\Sigma=J^{-1}(\boldsymbol{\delta})K(\boldsymbol{\delta})J^{-1}(\boldsymbol{\delta})$$

and

$$K_{l,q}=\sum_{i,i'}\mathrm{E}_{\boldsymbol{\delta}}\left[\left\{\frac{\partial}{\partial\delta_l}\log f_i(\boldsymbol{v_i}|\boldsymbol{\delta})\right\}\left\{\frac{\partial}{\partial\delta_q}\log f_{i'}(\boldsymbol{v_{i'}}|\boldsymbol{\delta})\right\}\right]$$

$$J_{l,q}=-\sum_{i}\mathrm{E}_{\boldsymbol{\delta}}\left[\frac{\partial^2}{\partial\delta_l\partial\delta_q}\log f_i(\boldsymbol{v_i}|\boldsymbol{\delta})\right].$$

Thus we have shown that $\boldsymbol{\delta_{MCLE}}$ is consistent and asymptotically normal with variance-covariance matrix $\Sigma_{k,k}$.

### S1.2 Asymptotic equivalence of $\boldsymbol{\delta_{MCLE}}$ and $\tilde{\boldsymbol{\delta}}$

The $MAP_{\mathrm{CL}}$ estimator $\tilde{\boldsymbol{\delta}}$ that we propose in Equation (5) of the main text differs from $\boldsymbol{\delta_{MCLE}}$ in that prior distributions are placed on the parameters to form the posterior distribution,

$$\pi_M(\boldsymbol{\delta})=f(\boldsymbol{\theta})f(\boldsymbol{\tau})\prod_{i\in Q}L_i(\boldsymbol{\delta};\boldsymbol{v}_i),$$

where $f(\boldsymbol{\theta})$ and $f(\boldsymbol{\tau})$ are the prior density functions for the population size parameters and node ages, respectively, and $M$ is the number of sites. $\tilde{\boldsymbol{\delta}}$ is the value that maximizes this posterior distribution, and we wish to show that $\tilde{\boldsymbol{\delta}}$ and $\boldsymbol{\delta_{MCLE}}$ are asymptotically equivalent.

In the setting in which inference is based on the true likelihood, rather than on the composite likelihood, the well-known Bernstein-von Mises Theorem van der Vaart (1998) can be applied to show that the MAP estimator converges to the maximum likelihood estimator (MLE) under certain regularity conditions, and thus that the MAP estimator shares the asymptotic distribution of the MLE. The derivation of similar results in the setting of composite likelihood is a currently active research area (see, e.g., Bassett and Deride (2019); Miller (2021)). In particular, the results of Miller (2021) can be applied to support our claim of consistency of $\tilde{\boldsymbol{\delta}}$ and of asymptotic equivalence of $\boldsymbol{\delta_{MCLE}}$ and $\tilde{\boldsymbol{\delta}}$, as described below. We begin by stating the relevant result from Miller (2021).

**Theorem** (Miller, 2021)**.** *Let $\Theta \subseteq R^D$. Let $E \subseteq \Theta$ be open (in $R^D$), convex, and bounded. Let $\theta_0 \in E$ and let $\pi : \Theta \to R$ be a probability density with respect to Lebesgue measure such that $\pi$ is continuous at $\theta_0$ and $\pi(\theta_0) > 0$. Let $h_n : \Theta \to R$ have continuous third derivatives on $E$. Suppose $h_n \to h$ pointwise for some $h : \Theta \to R$, $h^{''}(\theta_0)$ is positive definite, and $h_n^{'''}$ is uniformly bounded on $E$. If either of the following two conditions is satisfied:*

1. *$h(\theta) > h(\theta_0)$ for all $\theta \in K \backslash \theta_0$ and $liminf_n \; inf_{\theta\in\Theta\backslash K} h_n(\theta) > h(\theta_0)$ for some compact $K \subseteq E$ with $\theta_0$ in the interior of $K$, or*
2. *each $h_n$ is convex and $h^{'}(\theta_0) = 0$,*

*then there is a sequence $\theta_n \to \theta_0$ such that $h_n^{'}(\theta_n) = 0$ for all $n$ sufficiently large, $h_n(\theta_n) \to h(\theta_0)$, $\pi_n$ concentrates at $\theta_0$, and $\sqrt{n}(\theta - \theta_n)$ converges to $N(0, H_0^{-1})$ in total variation, where the random variable $\theta \sim \pi_n$ and $H_0 = h^{''}(\theta_0)$.*

As above, denote the true parameter values as $\boldsymbol{\delta}_0$, and let $\boldsymbol{\delta} \sim \pi_M$. Then, since $\boldsymbol{\delta_{MCLE}}$ is statistically consistent (see Section S1.1) and the conditions of the above theorem are satisfied, the posterior composite likelihood $\pi_M$ concentrates at $\boldsymbol{\delta}_0$.

**Proof:** To see this, consider the continuous function:

$$f_M = \sum_{i\in Q} f_{Mi} = \sum_{i\in Q} \bigg\{ -\frac{1}{M} \sum_{m=1}^{M} \log f_i(\boldsymbol{v}_i^{(m)}|\boldsymbol{\delta}) \bigg\}, \tag{3}$$

and assume that the specified joint priors $\pi(\boldsymbol{\delta}) = f(\boldsymbol{\theta})f(\boldsymbol{\tau})$ are continuous at $\boldsymbol{\delta_0}$ and are such that $\pi(\boldsymbol{\delta}_0) > 0$. Furthermore, we note that each $f_{Mi}$ in equation (3) is the log likelihood for a multinomial sample for quartet $i$. Therefore,

$$f_{Mi} = -\frac{1}{M}\sum_{j=1}^{9} m_{ij}\log p_{ij}(\boldsymbol{\delta}) \quad \text{if the } i^{th} \text{ quartet is symmetric or}$$

$$f_{Mi} = -\frac{1}{M}\sum_{j=1}^{11} m_{ij}\log p_{ij}(\boldsymbol{\delta}) \quad \text{if the } i^{th} \text{ quartet is asymmetric,}$$

where $m_{ij}$ is the number of sites with pattern $j$ for quartet $i$, $\sum m_{ij} = M$, and the probability for each category $p_{ij}(\boldsymbol{\delta})$ is in the form shown in Supplement A of Chifman and Kubatko (2015). We can easily check that every $p_{ij}(\boldsymbol{\delta})$ has continuous derivatives of all order in the parameter space $\Theta$ for which all parameters satisfy the constraints of the phylogenetic setting, i.e., the effective population size parameter is positive and the node ages are ordered such that all branch lengths are positive. Next, we define

$$f(\boldsymbol{\delta}) = \sum_{i\in Q}\mathbb{E}(f_{Mi}) = -\sum_{i\in Q}\sum_{j} p_{ij}(\boldsymbol{\delta_0})\log p_{ij}(\boldsymbol{\delta}),$$

and note that since $\big\{\frac{m_{ij}}{M}|i = 1,\cdots,|Q|,\ j = 1,\cdots,9\ \mathit{or}\ 11\big\}$ are sufficient statistics for $\boldsymbol{\delta}$, and we have

$$\frac{m_{ij}}{M} \to p_{ij}(\boldsymbol{\delta_0}) \quad \text{almost surely.}$$

Thus, we have that with probability 1, for all $\boldsymbol{\delta} \in \Theta$, $f_M(\boldsymbol{\delta}) \to f(\boldsymbol{\delta})$. The second derivatives are:

$$\frac{\partial^2 f(\boldsymbol{\delta})}{\partial\boldsymbol{\delta}\partial\boldsymbol{\delta}^T} = \sum_{i\in Q}\frac{\partial^2}{\partial\boldsymbol{\delta}\partial\boldsymbol{\delta}^T}\mathbb{E}(f_{Mi}) = \sum_{i\in Q}\mathbb{E}\Big(\frac{\partial^2 f_{Mi}}{\partial\boldsymbol{\delta}\partial\boldsymbol{\delta}^T}\Big).$$

Since for each $f_{Mi}$, the score function satisfies

$$\mathbb{E}\bigg(-\frac{\partial M f_{Mi}(\boldsymbol{\delta})}{\partial\boldsymbol{\delta}}\bigg|_{\boldsymbol{\delta}=\boldsymbol{\delta_0}}\bigg) = 0, \tag{4}$$

the hessian matrix satisfies

$$\mathbb{E}\bigg(\frac{\partial^2 f_{Mi}}{\partial\boldsymbol{\delta}\partial\boldsymbol{\delta}^T}\bigg)\bigg|_{\boldsymbol{\delta}=\boldsymbol{\delta_0}} = M\mathbb{E}\bigg(\frac{\partial f_{Mi}(\boldsymbol{\delta})}{\partial\boldsymbol{\delta}}\Big(\frac{\partial f_{Mi}(\boldsymbol{\delta})}{\partial\boldsymbol{\delta}}\Big)^T\bigg)\bigg|_{\boldsymbol{\delta}=\boldsymbol{\delta_0}} \succ 0.$$

Therefore, $f^{''}(\boldsymbol{\delta_0})$ is positive definite because it is a sum of positive definite matrices. Furthermore, if we let $E$ be an open ball such that $\boldsymbol{\delta_0} \in E$ and $\bar{E} \subseteq \Theta$, since $f_M^{'''}$ is continuous, then it is also uniformly bounded on the compact set $\bar{E}$.

Having established that our formulation matches the criteria of the theorem, we now show that condition (1) is satisfied in our case. First, since we have shown above that $f^{''}(\boldsymbol{\delta_0}) \succ 0$ and $f^{'}(\boldsymbol{\delta_0}) = 0$ from equation (4), we know that $\boldsymbol{\delta_0}$ is a local minimum, which means that $f(\boldsymbol{\delta}) > f(\boldsymbol{\delta}_0)$ for all $\boldsymbol{\delta}$ in a compact set $K$. Also, from the above result, we have with probability 1, for all $\boldsymbol{\delta} \in \Theta$, $f_M \to f$:

$$\lim_{n\to\infty} Pr(|f_M - f| < \epsilon) = 1,$$

so with probability 1,

$$\liminf_M \inf_{\boldsymbol{\delta}\in\Theta} f_M(\boldsymbol{\delta}) = \liminf_M f_M(\boldsymbol{\delta}_{MCLE}) = f(\boldsymbol{\delta}_{MCLE}).$$

Since we know $\boldsymbol{\delta}_{MCLE}$ is a consistent estimator of $\boldsymbol{\delta}_0$ from Section S1.1, then with probability 1,

$$\liminf_M \inf_{\boldsymbol{\delta}\in\Theta\setminus K} f_M(\boldsymbol{\delta}) > \liminf_M \inf_{\boldsymbol{\delta}\in\Theta} f_M(\boldsymbol{\delta}) = f(\boldsymbol{\delta}_0).$$

Thus condition (1) of the theorem is satisfied, and we have therefore proved that $\sqrt{M}(\boldsymbol{\delta} - \boldsymbol{\delta}_{MCLE}) \to N(0, H_0^{-1})$ in total variation distance, where $H_0 = f''(\boldsymbol{\delta}_0)$, and that the posterior composite likelihood $\pi_M$ concentrates at $\boldsymbol{\delta_0}$ as $M \to \infty$. □

Thus, we know that the density of the posterior composite likelihood becomes concentrated on the true parameter value as the number of sites grows. From Section S1.1, we know that the maximum of the composite likelihood is a consistent estimator of the true parameter value that is asymptotically normally distributed. We thus conjecture that the distribution of the MAP estimate based on the posterior composite likelihood coincides with that of the maximum composite likelihood estimator. Our application of the asymptotic variance formula from Section S1.1 makes the assumption that this relationship holds, and our simulation results support this claim. Nonetheless, we ultimately recommend using the bootstrap for variance estimation, and thus the estimates reported in this paper do not depend on this claim. We note that the proof provided in this section does establish consistency of the $MAP_{\mathrm{CL}}$ estimator directly.

### S2. Simulation Study 1: Statistical Properties of $MAP_{\mathrm{CL}}$ Estimator

#### S2.1 Unlinked CIS data (one lineage per tip)

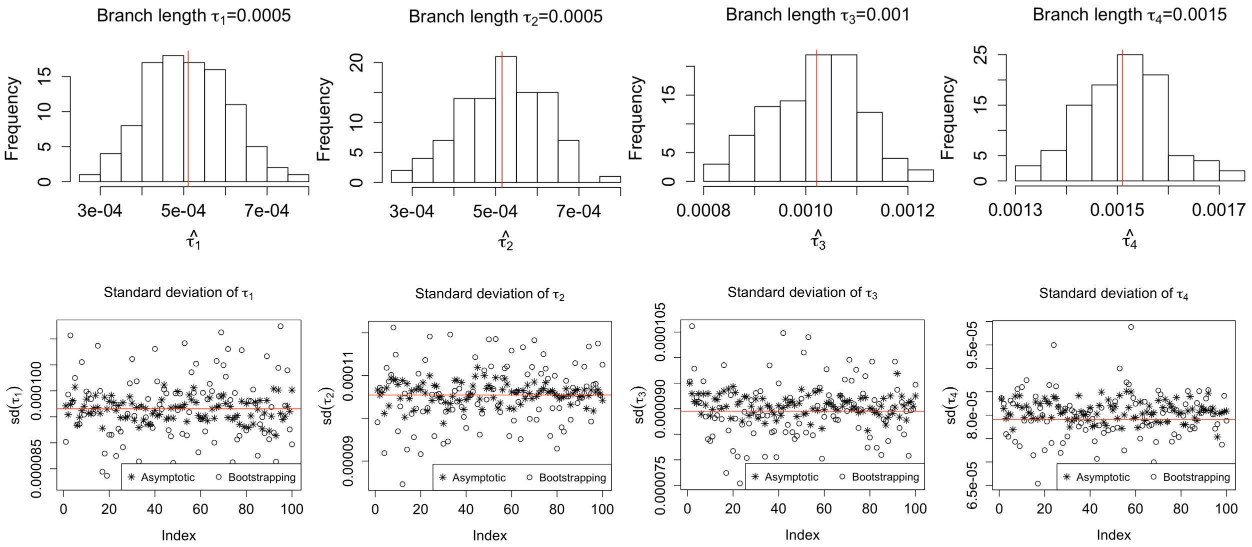


Figure S1: Plots of 100 $MAP_{\mathrm{CL}}$ estimates and their variance estimates. The 5-taxon model tree in Figure 2(a) with node ages ($\tau_1 = 0.5, \tau_2 = 0.5, \tau_3 = 1.0, \tau_4 = 1.5$) is prespecified to generate 100,000 unlinked CIS data for one lineage per tip. The histograms in the first row show the unbiasedness and asymptotically normal property. The red line in each histogram represents the sample mean of the 100 $MAP_{\mathrm{CL}}$ estimates. Points in variance plots (second row) denoted by $*$ are computed by asymptotic variance formula in (5), while points denoted by $o$ are obtained by bootstrapping. The x-axis is an index for the simulated samples. The red line in the plots is the sample variance of the 100 $MAP_{\mathrm{CL}}$ estimates.

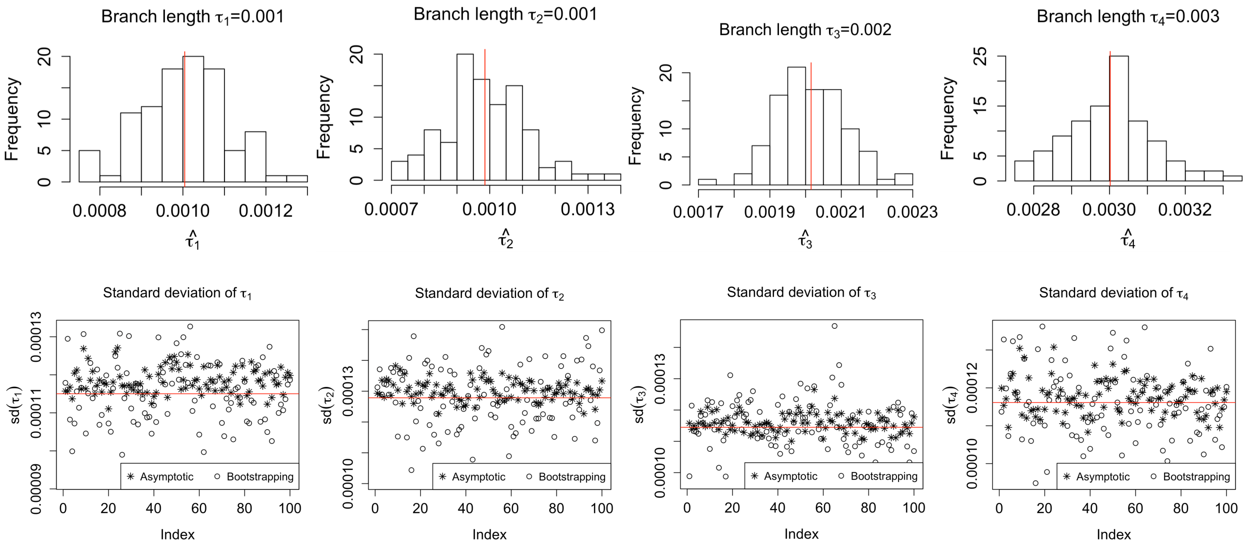


Figure S2: Plots of 100 $MAP_{\text{CL}}$ estimates and their variance estimates. The 5-taxon model tree in Figure 2(a) with node ages ($\tau_1 = 1.0, \tau_2 = 1.0, \tau_3 = 2.0, \tau_4 = 3.0$) is prespecified to generate 100,000 unlinked CIS data for one lineage per tip.

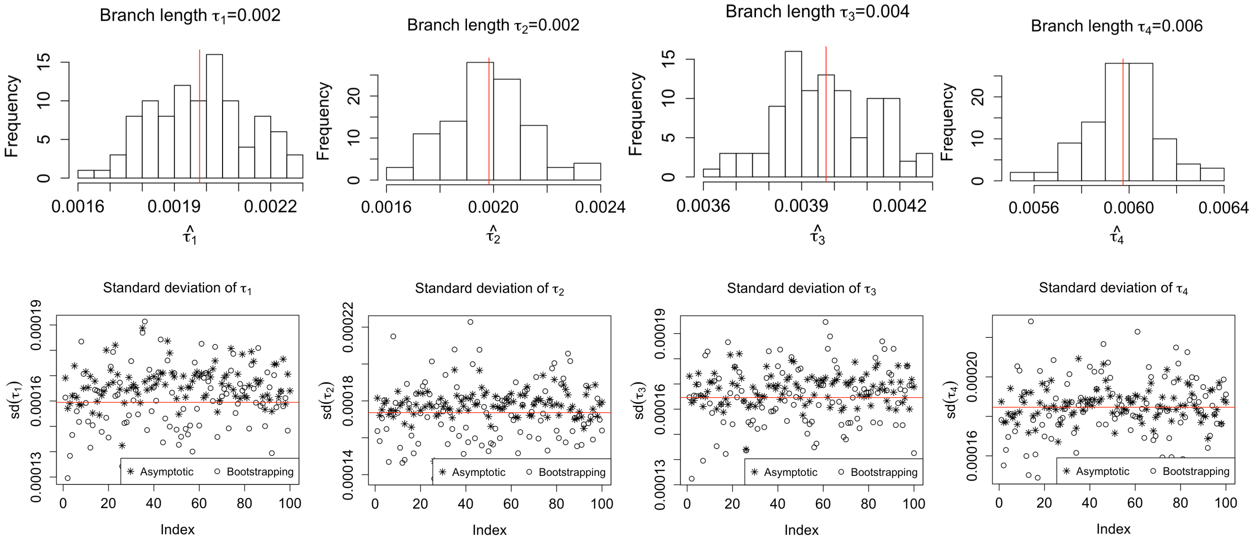


Figure S3: Plots of 100 $MAP_{\text{CL}}$ estimates and their variance estimates. The 5-taxon model tree in Figure 2(a) with node ages ($\tau_1 = 2.0, \tau_2 = 2.0, \tau_3 = 4.0, \tau_4 = 6.0$) is prespecified to generate 100,000 unlinked CIS data for one lineage per tip.

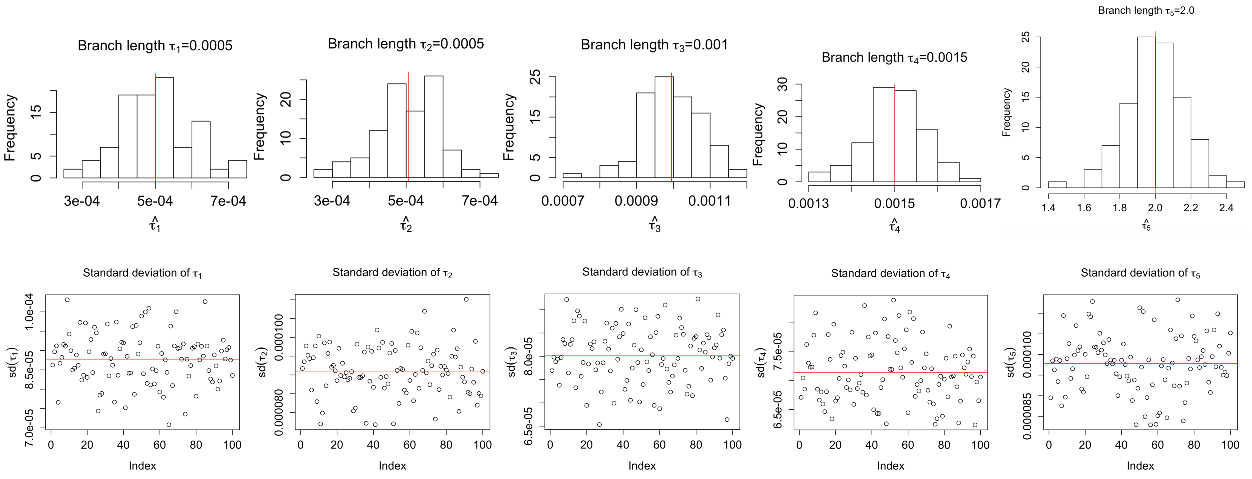


Figure S4: Plots of 100 $MAP_{\mathrm{CL}}$ estimates and their variance estimates. The 6-taxon model tree in Figure 2(b) with node ages ($\tau_1 = 0.5, \tau_2 = 0.5, \tau_3 = 1.0, \tau_4 = 1.5, \tau_5 = 2.0$) is prespecified to generate 100,000 unlinked CIS data for one lineage per tip.

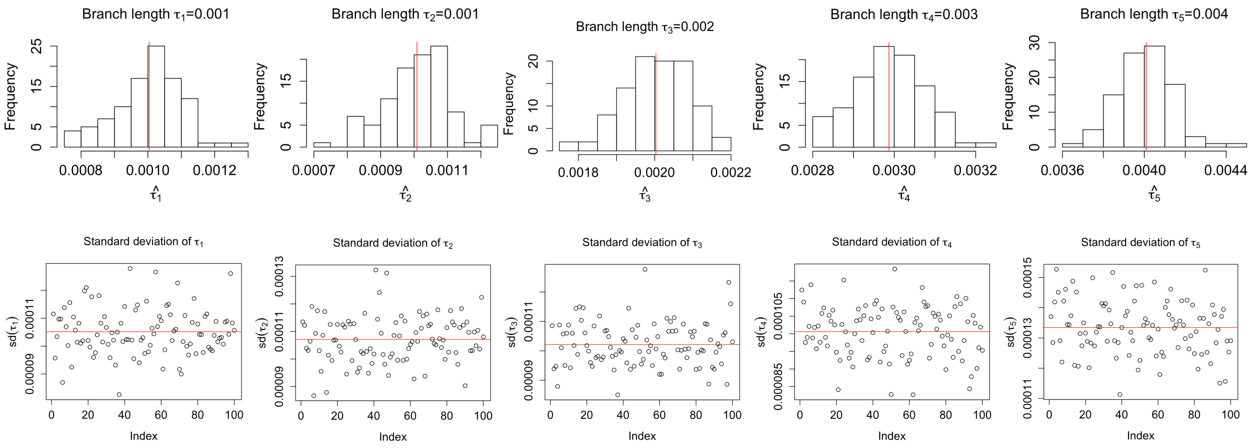


Figure S5: Plots of 100 $MAP_{\mathrm{CL}}$ estimates and their variance estimates. The 6-taxon model tree in Figure 2(b) with node ages ($\tau_1 = 1.0, \tau_2 = 1.0, \tau_3 = 2.0, \tau_4 = 3.0, \tau_5 = 4.0$) is prespecified to generate 100,000 unlinked CIS data for one lineage per tip.

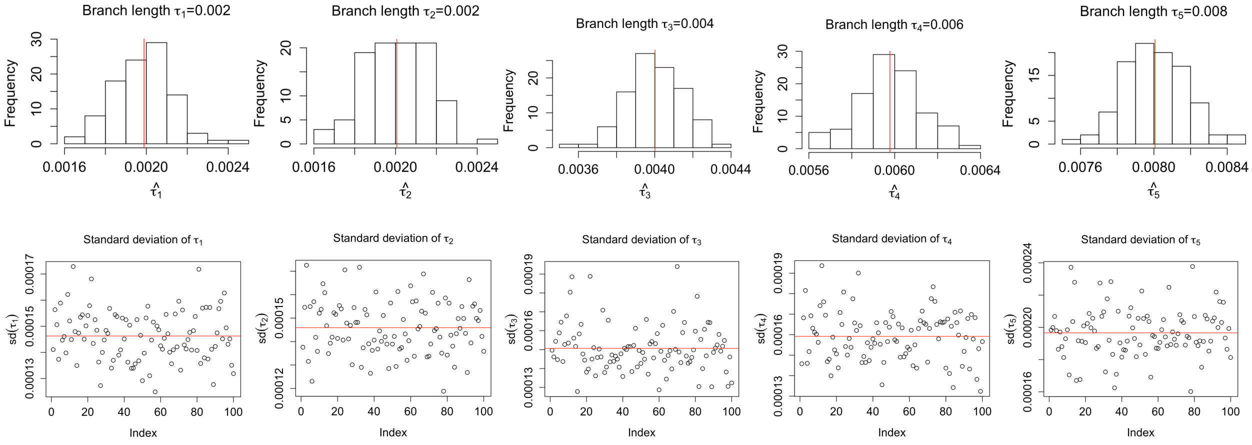


Figure S6: Plots of 100 $MAP_{\text{CL}}$ estimates and their variance estimates. The 6-taxon model tree in Figure 2(b) with node ages ($\tau_1 = 2.0, \tau_2 = 2.0, \tau_3 = 4.0, \tau_4 = 6.0, \tau_5 = 8.0$) is prespecified to generate 100,000 unlinked CIS data for one lineage per tip.

### S2.2 Multi-locus data

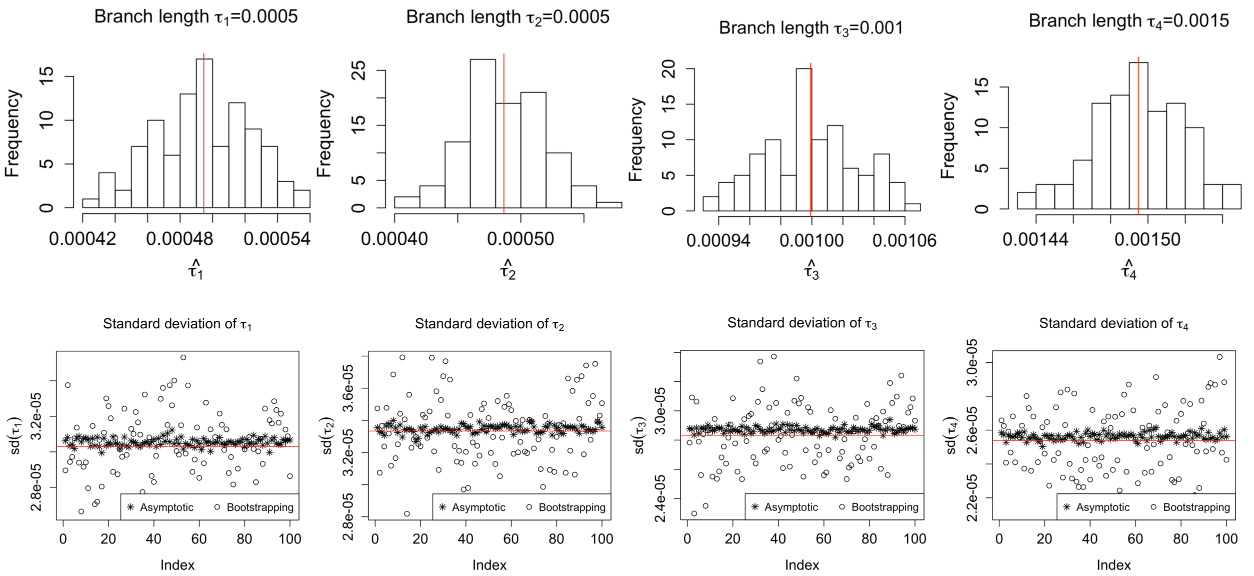


Figure S7: Plots of 100 $MAP_{\text{CL}}$ estimates and their variance estimates. The 5-taxon model tree in Figure 2(a) with node ages ($\tau_1 = 0.5, \tau_2 = 0.5, \tau_3 = 1.0, \tau_4 = 1.5$) is prespecified to generate 1,000,000 multi-locus data (10,000 genes, each of length 100).

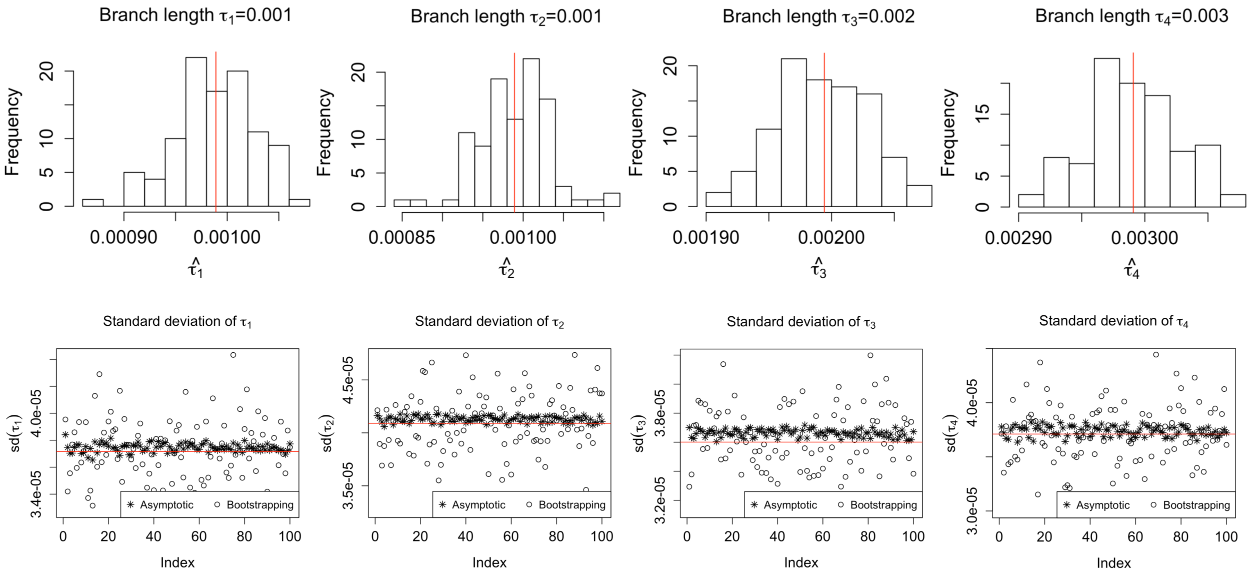


Figure S8: Plots of 100 $MAP_{\text{CL}}$ estimates and their variance estimates. The 5-taxon model tree in Figure 2(a) with node ages ($\tau_1 = 1.0, \tau_2 = 1.0, \tau_3 = 2.0, \tau_4 = 3.0$) is prespecified to generate 1,000,000 multi-locus data (10,000 genes, each of length 100).

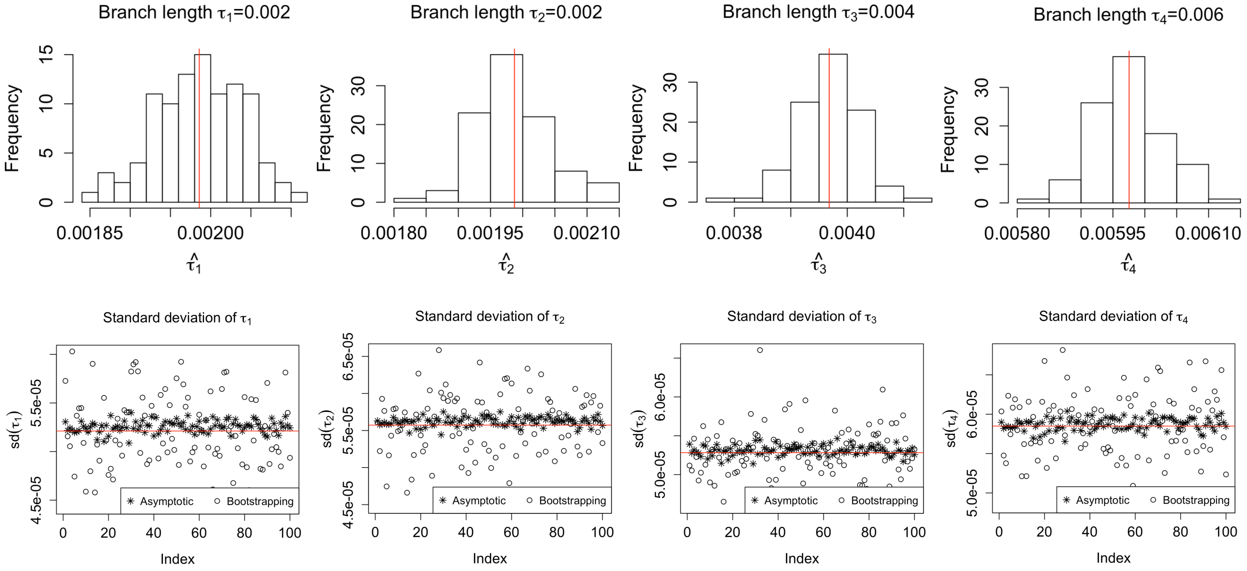

Figure S9: Plots of 100 $MAP_{\mathrm{CL}}$ estimates and their variance estimates. The 5-taxon model tree in Figure 2(a) with node ages ($\tau_1 = 2.0, \tau_2 = 2.0, \tau_3 = 4.0, \tau_4 = 6.0$) is prespecified to generate 1,000,000 multi-locus data (10,000 genes, each of length 100).

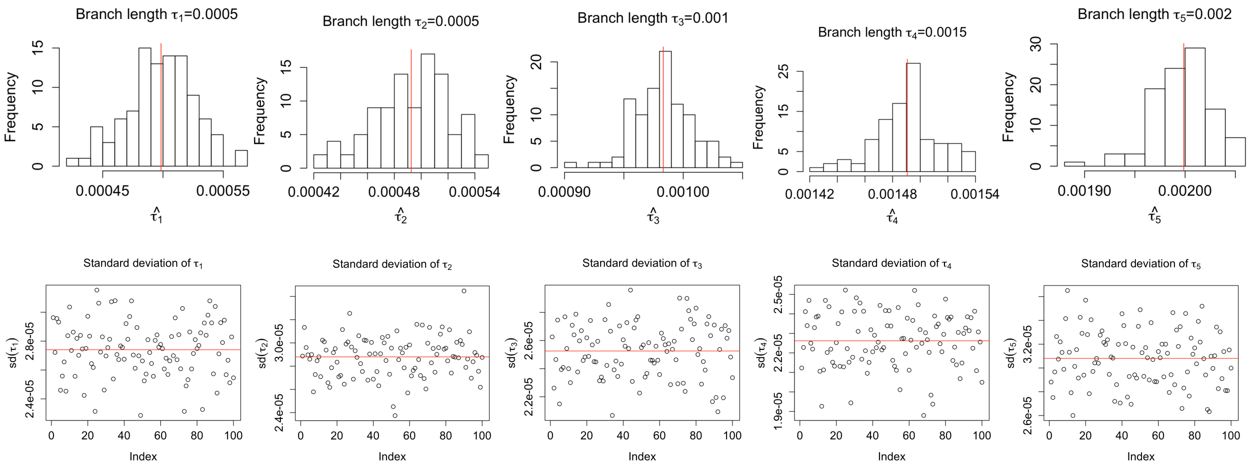

Figure S10: Plots of 100 $MAP_{\mathrm{CL}}$ estimates and their variance estimates. The 6-taxon model tree in Figure 2(b) with node ages ($\tau_1 = 0.5, \tau_2 = 0.5, \tau_3 = 1.0, \tau_4 = 1.5, \tau_5 = 2.0$) is prespecified to generate 1,000,000 multi-locus data (10,000 genes, each of length 100).

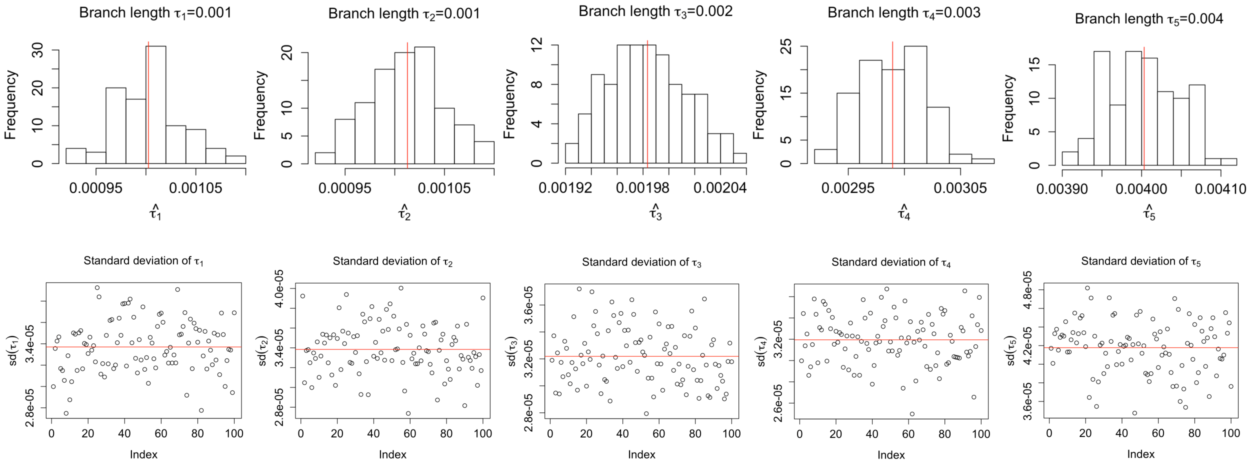


Figure S11: Plots of 100 $MAP_{\mathrm{CL}}$ estimates and their variance estimates. The 6-taxon model tree in Figure 2(b) with node ages ($\tau_1 = 1.0, \tau_2 = 1.0, \tau_3 = 2.0, \tau_4 = 3.0, \tau_5 = 4.0$) is prespecified to generate 1,000,000 multi-locus data (10,000 genes, each of length 100).

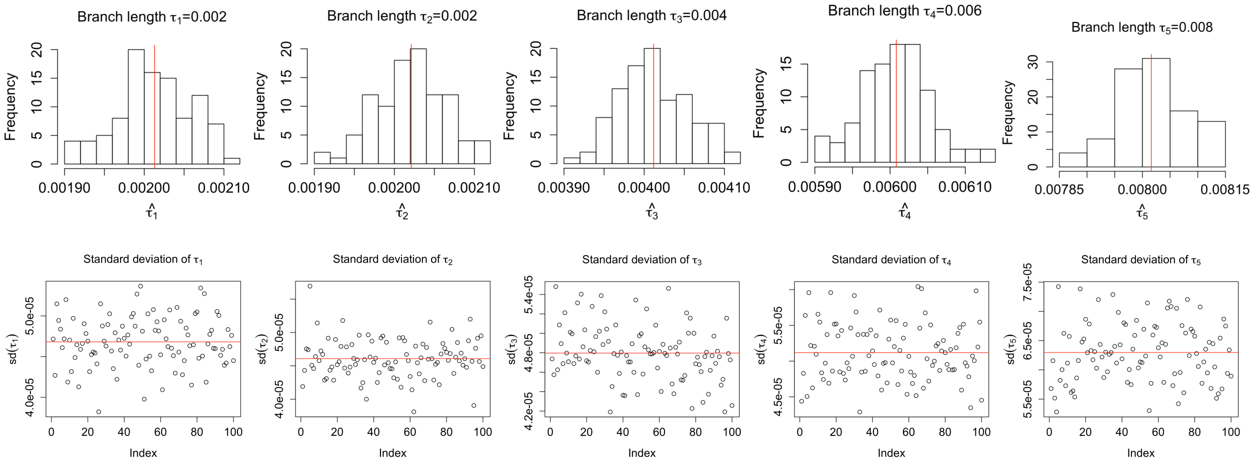


Figure S12: Plots of 100 $MAP_{\mathrm{CL}}$ estimates and their variance estimates. The 6-taxon model tree in Figure 2(b) with node ages ($\tau_1 = 2.0, \tau_2 = 2.0, \tau_3 = 4.0, \tau_4 = 6.0, \tau_5 = 8.0$) is prespecified to generate 1,000,000 multi-locus data (10,000 genes, each of length 100).

#### S2.3 Multi-locus data (multiple lineages per tip)

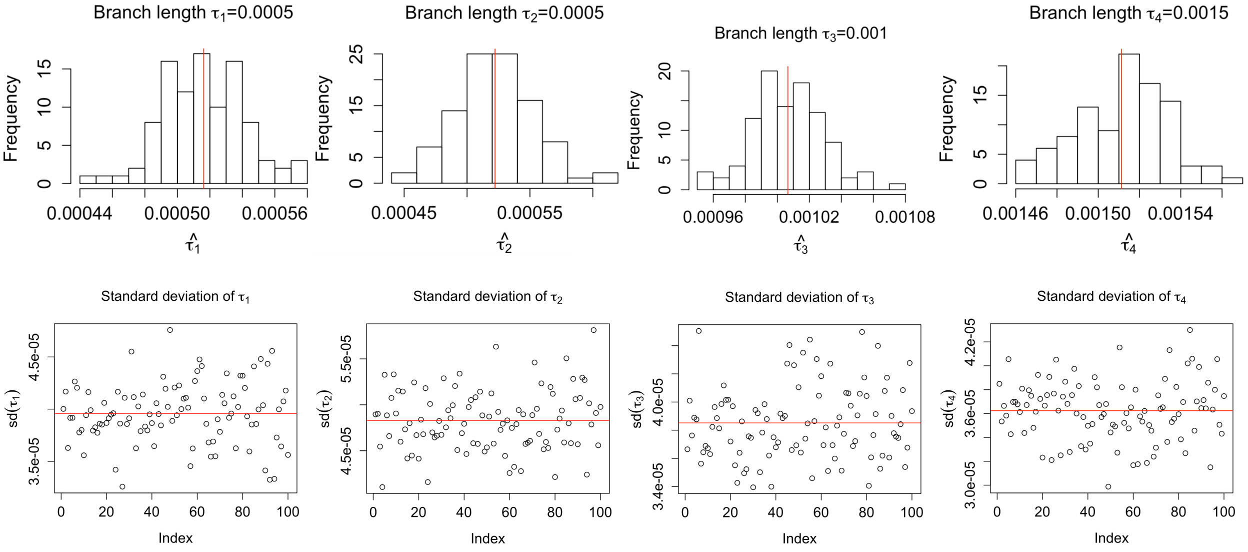


Figure S13: Plots of 100 $MAP_{\mathrm{CL}}$ estimates and their variance estimates. The 5-taxon model tree with node ages ($\tau_1 = 0.5, \tau_2 = 0.5, \tau_3 = 1.0, \tau_4 = 1.5$) is prespecified to generate 1,000,000 multi-locus data (10,000 genes, each of length 100), but 2 lineages for species D and E in Figure 2(a). The scatter plots in the second row show the performance of bootstrap variance estimates.

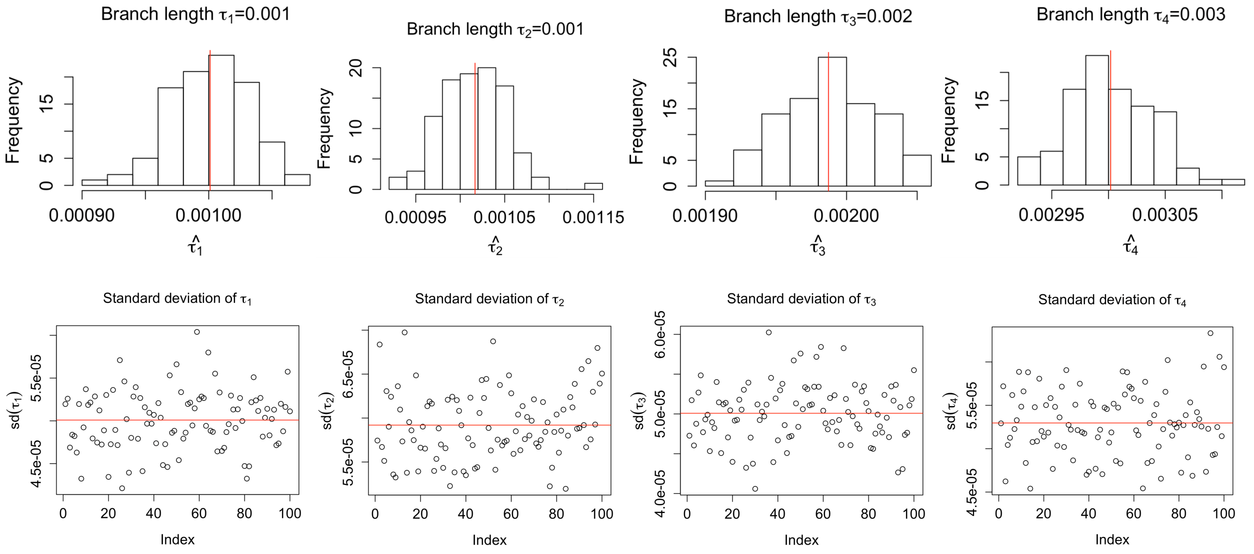


Figure S14: Plots of 100 $MAP_{\mathrm{CL}}$ estimates and their variance estimates. The 5-taxon model tree with node ages ($\tau_1 = 1.0, \tau_2 = 1.0, \tau_3 = 2.0, \tau_4 = 3.0$) is prespecified to generate 1,000,000 multi-locus data (10,000 genes, each of length 100), but 2 lineages for species D and E in Figure 2(a).

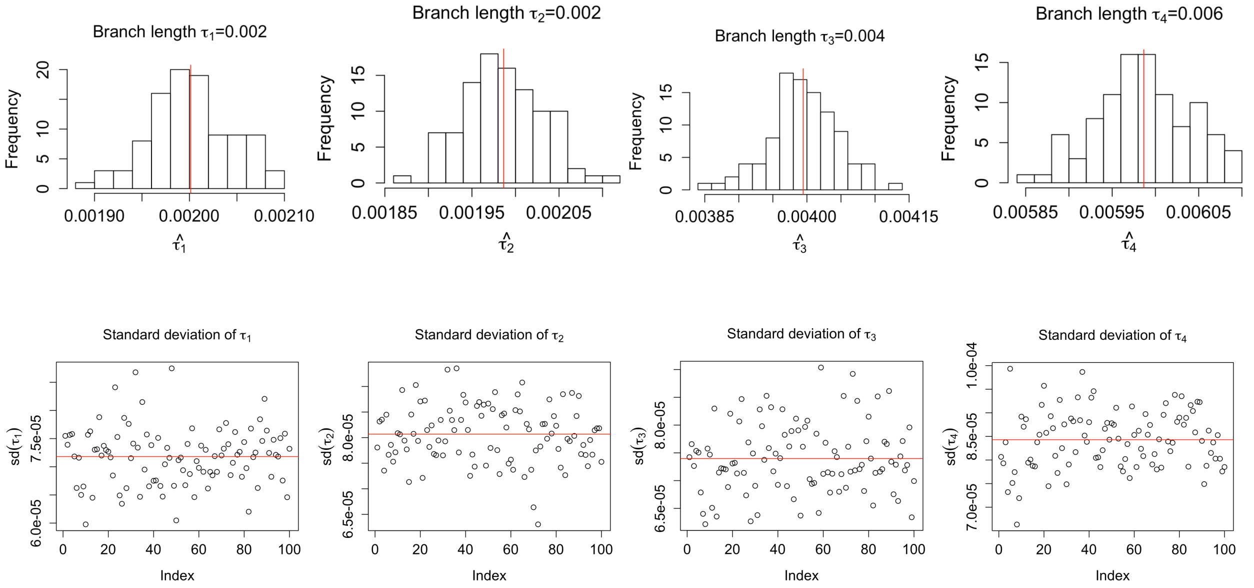


Figure S15: Plots of 100 $MAP_{\mathrm{CL}}$ estimates and their variance estimates. The 5-taxon model tree with node ages ($\tau_1 = 2.0, \tau_2 = 2.0, \tau_3 = 4.0, \tau_4 = 6.0$) is prespecified to generate 1,000,000 multi-locus data (10,000 genes, each of length 100), but 2 lineages for species D and E in Figure 2(a).

### S3. Simulation Study 2: Comparison with BPP

Table S1: The parameter configuration for running MCMC in BPP analysis. Running time for BPP and $MAP_{\mathrm{CL}}$ is shown below.

| number of tips | number of samples for summarization | sample frequency | burnin steps | Running time BPP | Running time $MAP_{\mathrm{CL}}$ |
|---|---|---|---|---|---|
| 7 | 500 | 50 | 1000 | 00:49:44 | 00:00:01 |
| 8 | 500 | 50 | 1000 | 01:18:47 | 00:00:01 |
| 9 | 500 | 50 | 1400 | 01:19:07 | 00:00:02 |
| 10 | 500 | 50 | 2400 | 01:59:10 | 00:00:03 |
| 11 | 500 | 54 | 3000 | 01:59:23 | 00:00:06 |
| 12 | 500 | 64 | 3600 | 02:32:29 | 00:00:08 |
| 13 | 500 | 75 | 4200 | 03:26:32 | 00:00:13 |
| 14 | 500 | 130 | 7200 | 06:17:18 | 00:00:21 |
| 15 | 500 | 170 | 9600 | 09:01:09 | 00:00:29 |
| 20 | 500 | 432 | 24000 | 27:10:50 | 00:01:31 |

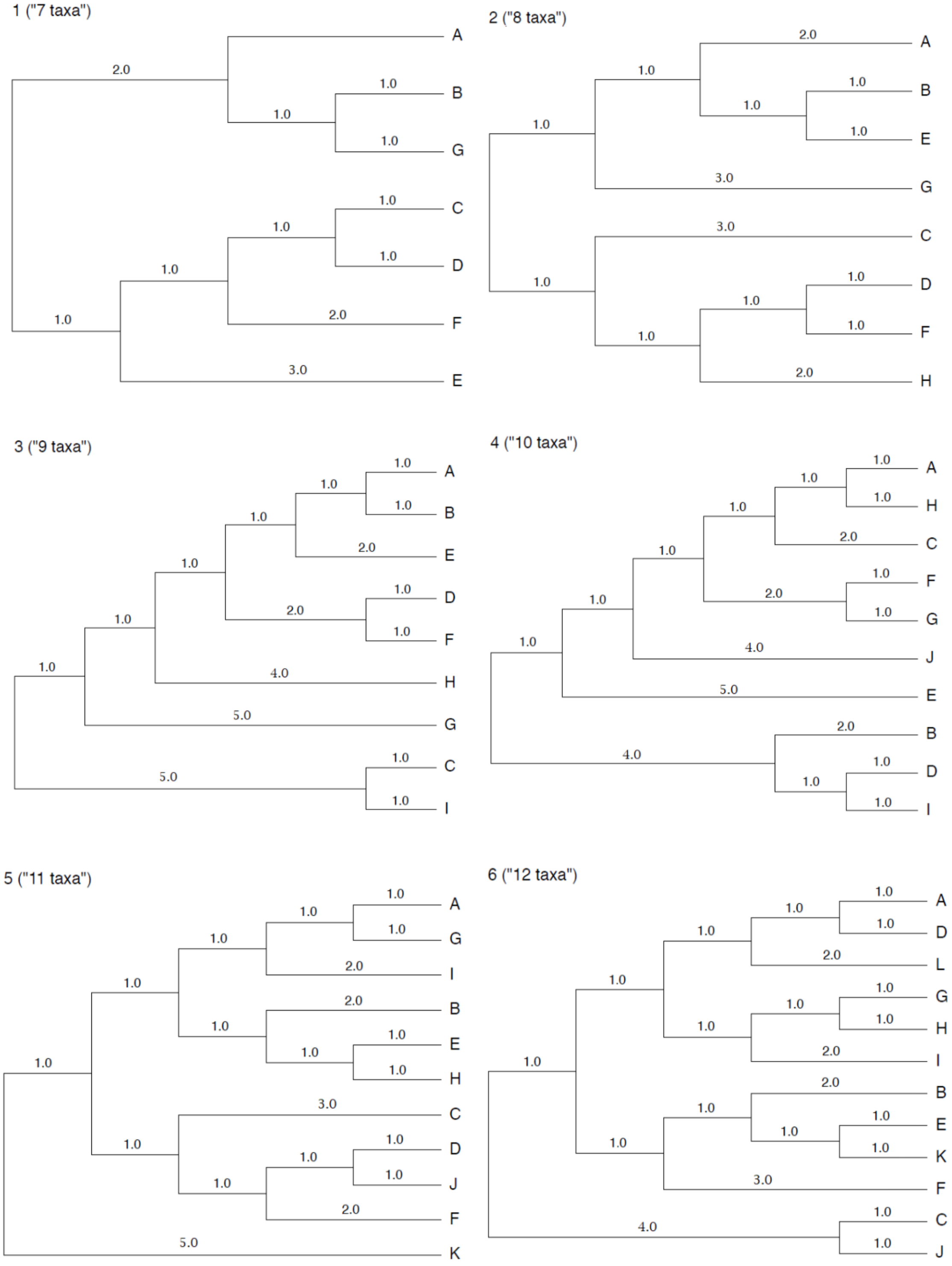


Figure S16: Trees used to carry out the simulation-based comparison with BPP for 7 - 12 taxa.

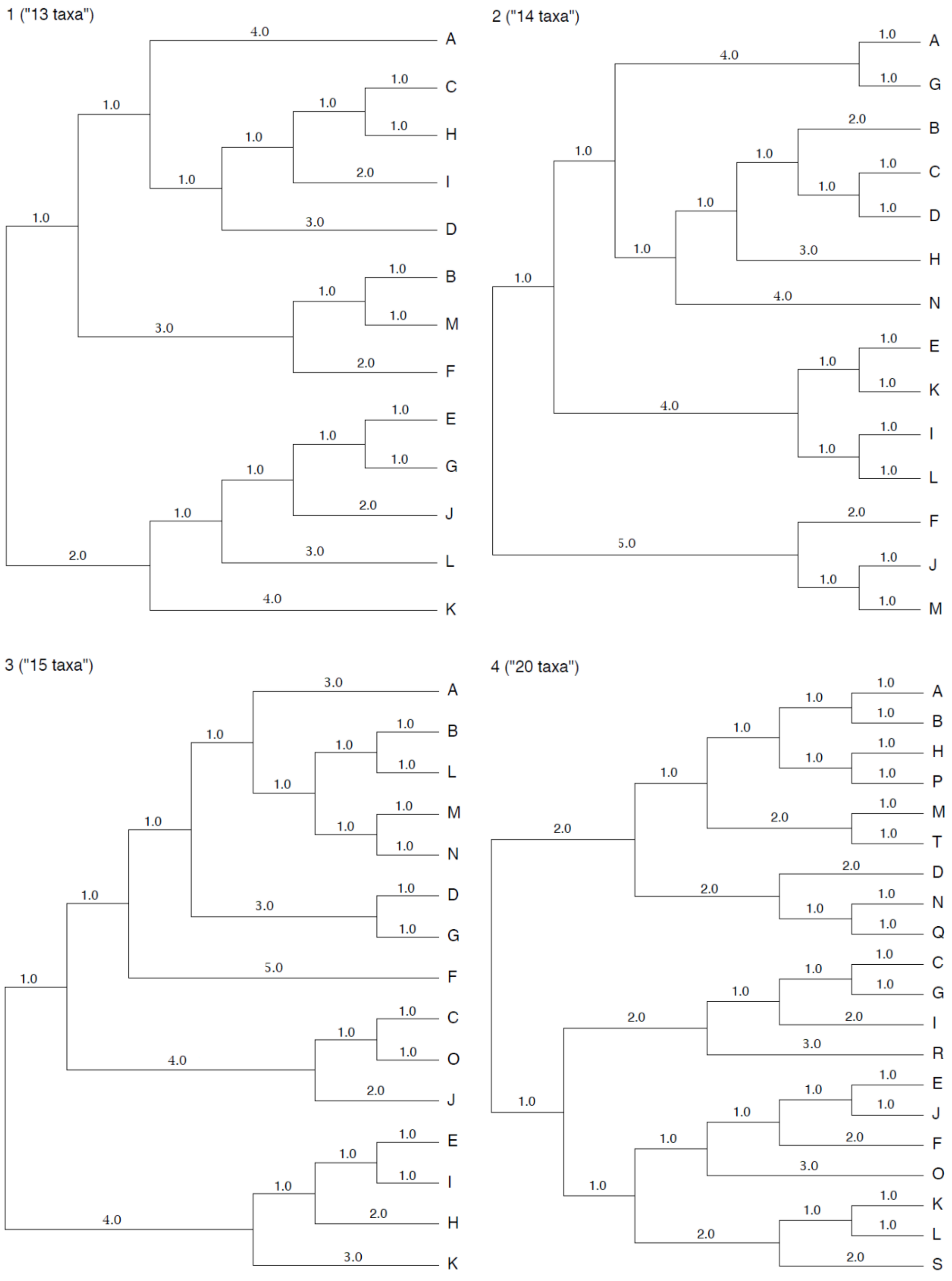


Figure S17: Trees used to carry out the simulation-based comparison with BPP for 13, 14, 15, and 20 taxa.

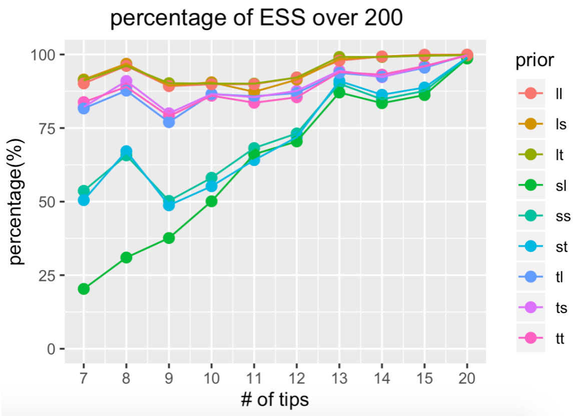


Figure S18: Percentage of ESS over 200 for BPP runs with different prior choices. X axis represents tree size with different number of tips. Prior ll corresponds to $\theta \sim IG(3, 0.02)$ and $\tau_{root} \sim$ IG(3,5*tree height); Prior ls corresponds to $\theta \sim IG(3, 0.02)$ and $\tau_{root} \sim$ IG(3,tree height/5); Prior lt corresponds to $\theta \sim IG(3, 0.02)$ and $\tau_{root} \sim$ IG(3,tree height); Prior sl corresponds to $\theta \sim IG(3, 0.0008)$ and $\tau_{root} \sim$ IG(3,5*tree height); Prior ss corresponds to $\theta \sim IG(3, 0.0008)$ and $\tau_{root} \sim$ IG(3,tree height/5); Prior st corresponds to $\theta \sim IG(3, 0.0008)$ and $\tau_{root} \sim$ IG(3,tree height); Prior tl corresponds to $\theta \sim IG(3, 0.004)$ and $\tau_{root} \sim$ IG(3,5*tree height); Prior ts corresponds to $\theta \sim IG(3, 0.004)$ and $\tau_{root} \sim$ IG(3,tree height/5); Prior tt corresponds to $\theta \sim IG(3, 0.004)$ and $\tau_{root} \sim$ IG(3,tree height);

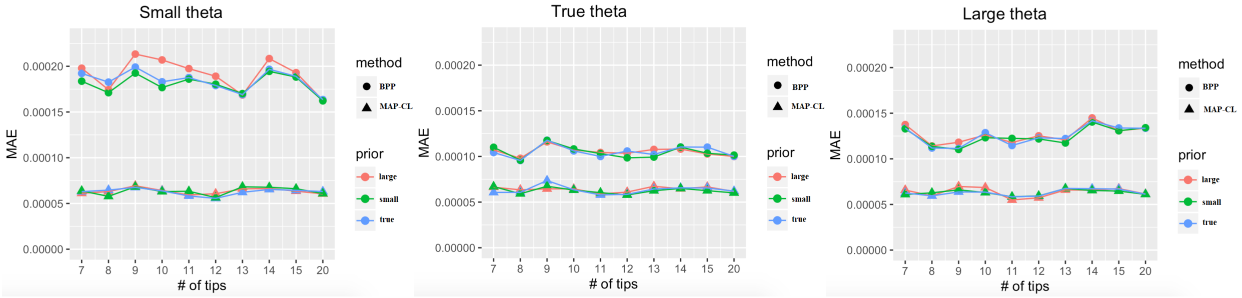


(a) $\theta \sim IG(3, 0.0008)$ (b) $\theta \sim IG(3, 0.004)$ (c) $\theta \sim IG(3, 0.02)$

Figure S19: Plots of MAE of node age estimates on trees with different sizes. X axis represents tree size with different number of tips. Analysis based on two methods (circle points – BPP, trangle points $-MAP_{\mathrm{CL}}$) is conducted with different priors. In each plot, large (in red): $\tau_{root} \sim$ IG(3, 5* tree height $h$); small (in green): $\tau_{root} \sim$ IG(3, tree height $h/5$); true (in blue): $\tau_{root} \sim$ IG(3, tree height $h$). Plots from (a) to (c) represents analysis under different priors for $\theta$: (a) $\theta \sim$ IG(3,0.0008); (b) $\theta \sim$ IG(3,0.004); (c) $\theta \sim$ IG(3,0.02).

### S4. Application to Gibbon Data

In this section, we provide information to support the analysis of the gibbon data described in the main text.

Table S2: To match the analysis done by Shi and Yang (2018) for the gibbon data set, we suggest the priors $\tau_{root} \sim IG(0.01, 1.0)$ (this corresponds to $\alpha = 3, \beta = 0.02$ in the parameterization used by BPP) and $\theta \sim IG(0.001, 1.0)$ (this corresponds to $\alpha = 3, \beta = 0.002$ in the parameterization used by BPP). We also considered priors with mean 5 times smaller and 5 times larger. All combinations of these priors were considered, and are labeled by a setting number, as shown in the table below.

| | $\tau_{root} \sim IG(0.002, 1.0)$ | $\tau_{root} \sim IG(0.01, 1.0)$ | $\tau_{root} \sim IG(0.05, 1.0)$ |
|---|---|---|---|
| $\theta \sim IG(0.0002, 1.0)$ | Setting 1 | Setting 2 | Setting 3 |
| $\theta \sim IG(0.001, 1.0)$ | Setting 4 | Setting 5 | Setting 6 |
| $\theta \sim IG(0.002, 1.0)$ | Setting 7 | Setting 8 | Setting 9 |

Table S3: Comparison of the node age estimates for BPP and $\text{MAP}_{CL}$ in mutation units.

| Age of | BPP | | $MAP_{CL}$ | |
|---|---|---|---|---|
| MRCA of ... | mean | 95% HPD interval | mean | 95% CI |
| HmHp | 0.00088 | (0.00084, 0.00093) | 0.00085 | (0.00081, 0.00090) |
| BS | 0.00164 | (0.00154, 0.00175) | 0.00274 | (0.00270, 0.00279) |
| NBS | 0.00264 | (0.00248, 0.00280) | 0.00279 | (0.00275, 0.00284) |
| NBSHmHp | 0.00306 | (0.00302, 0.00311) | 0.00290 | (0.00286, 0.00294) |
| ONBSHmHp | 0.01148 | (0.01127, 0.01169) | 0.01428 | (0.01417, 0.01438) |

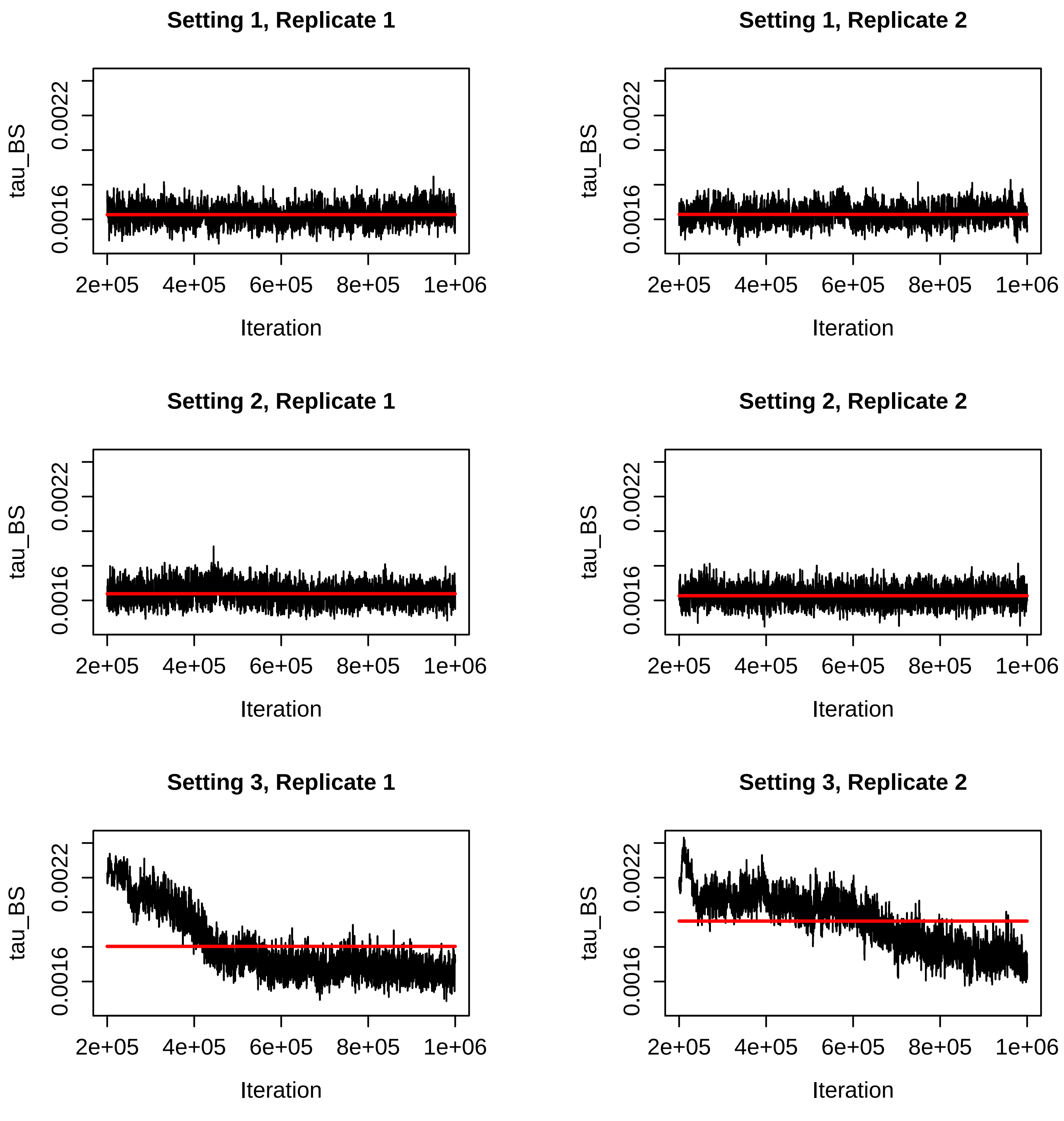


Figure S20: Trace plots for the $\tau_{BS}$ parameter for the BPP runs for Settings 1-3 in Table S2. The red line is the mean of the sampled values for that run.

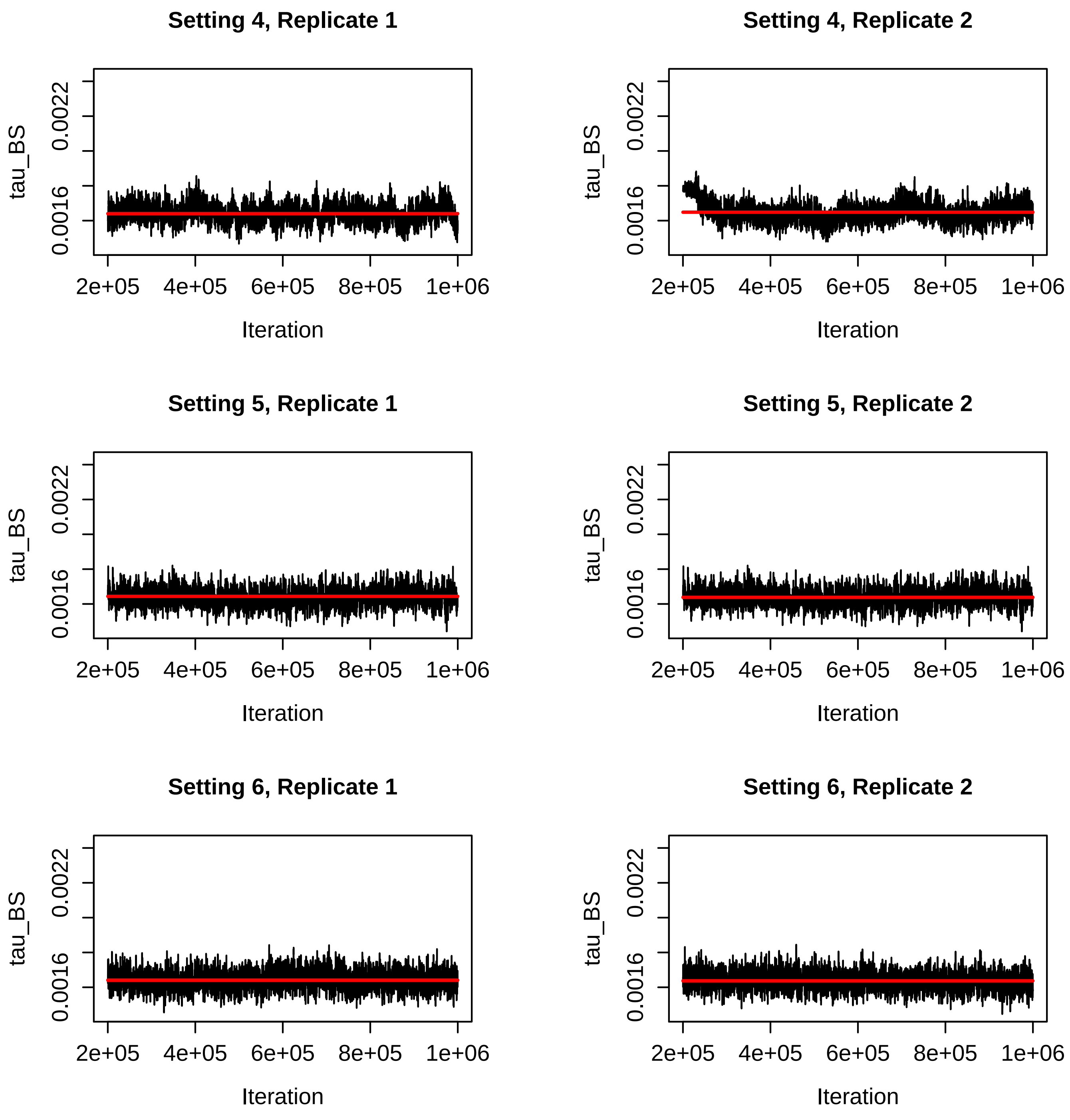


Figure S21: Trace plots for the $\tau_{BS}$ parameter for the BPP runs for Settings 4-6 in Table S2. The red line is the mean of the sampled values for that run.

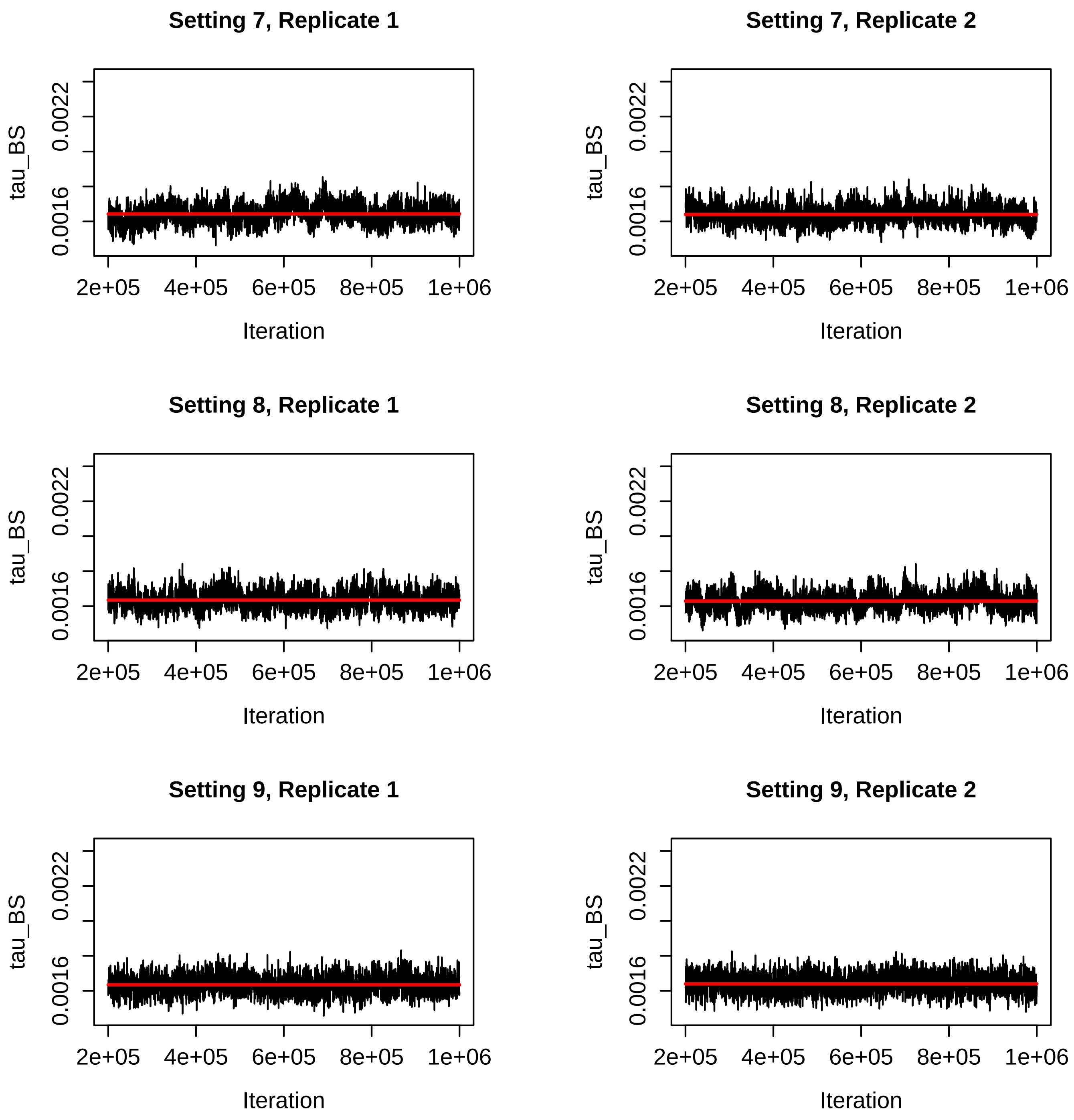


Figure S22: Trace plots for the $\tau_{BS}$ parameter for the BPP runs for Settings 7-9 in Table S2. The red line is the mean of the sampled values for that run.

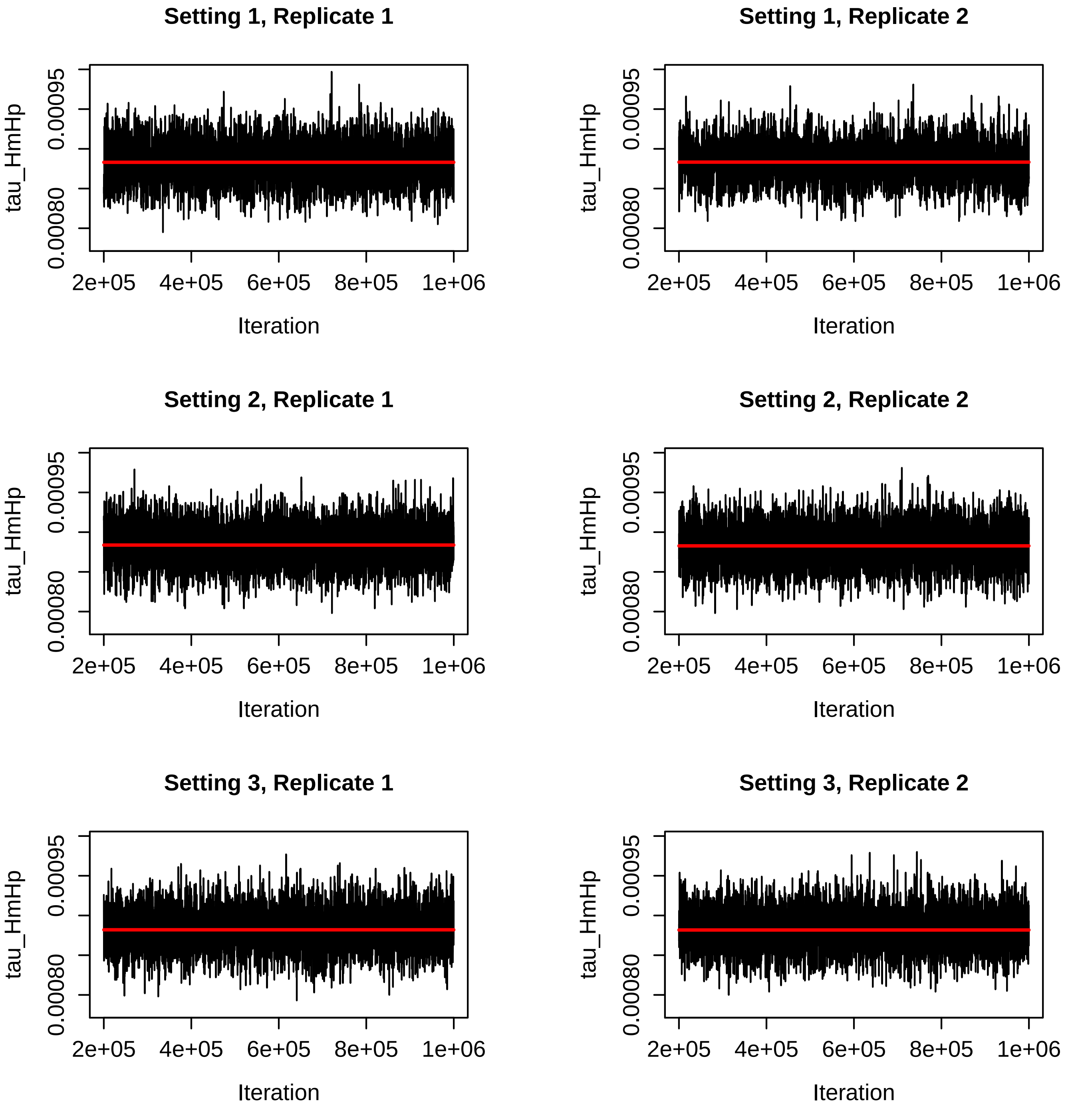


Figure S23: Trace plots for the $\tau_{HmHp}$ parameter for the BPP runs for Settings 1-3 in Table S2. The red line is the mean of the sampled values for that run.

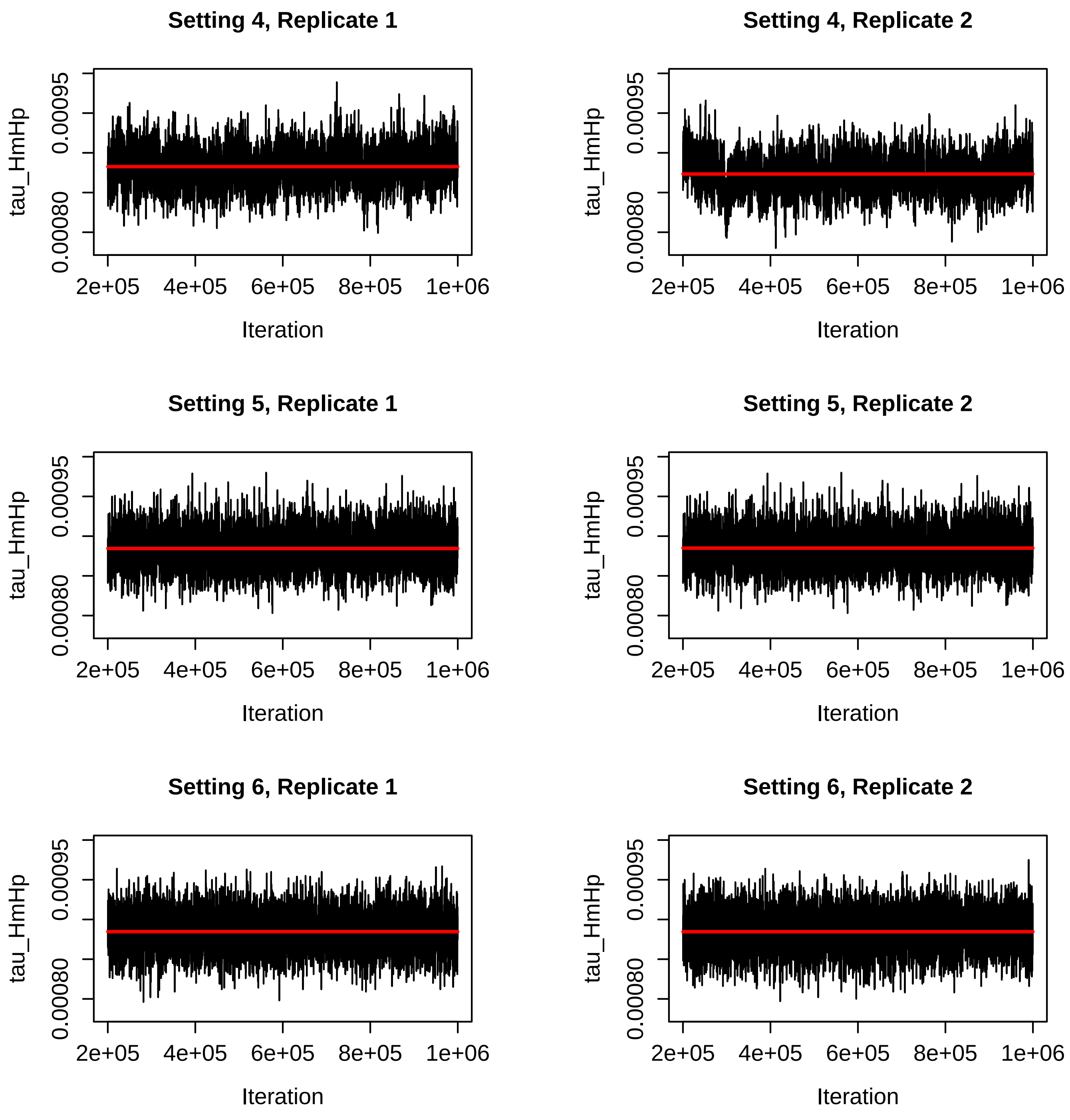


Figure S24: Trace plots for the $\tau_{HmHp}$ parameter for the BPP runs for Settings 4-6 in Table S2. The red line is the mean of the sampled values for that run.

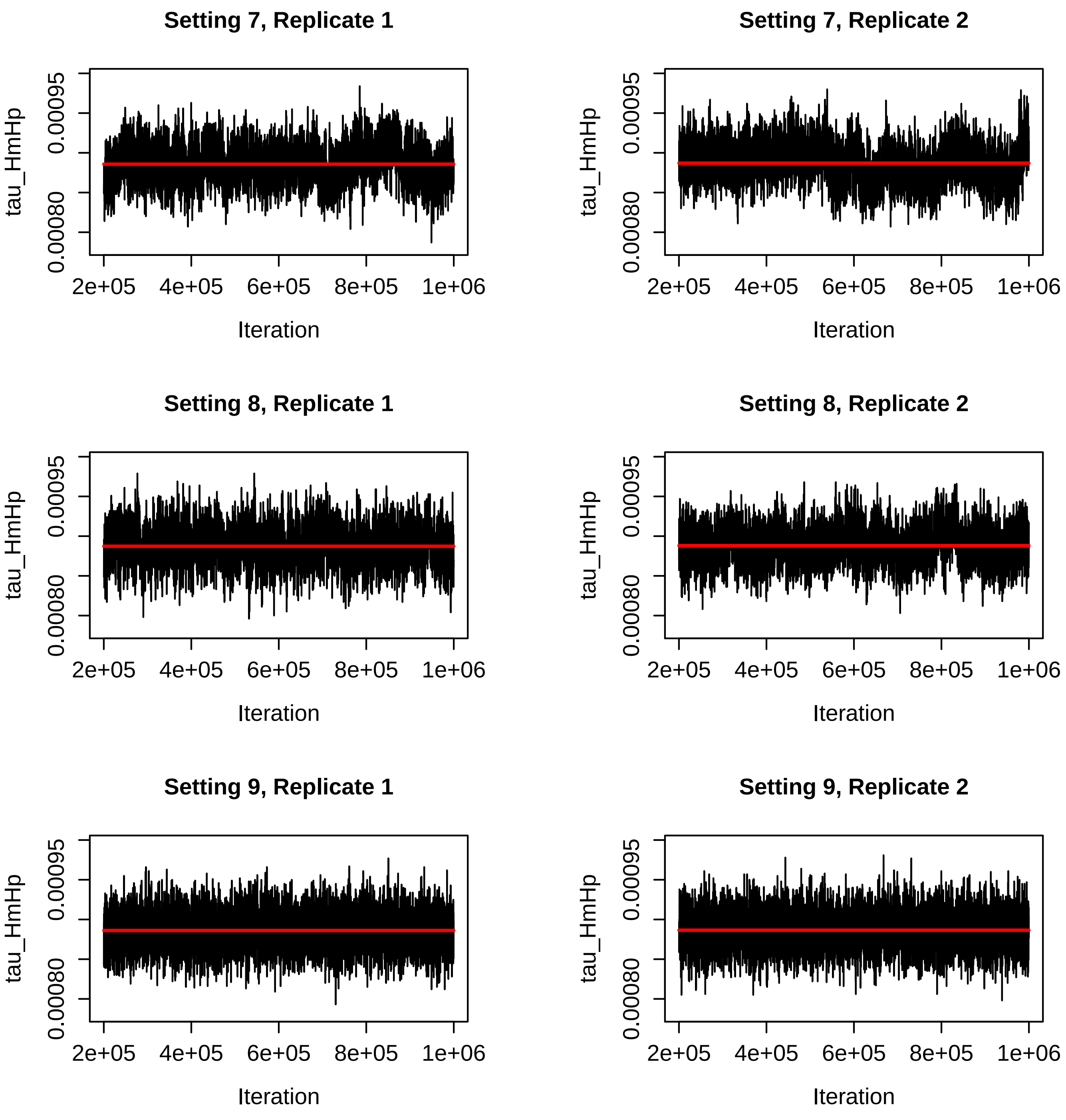


Figure S25: Trace plots for the $\tau_{HmHp}$ parameter for the BPP runs for Settings 7-9 in Table S2. The red line is the mean of the sampled values for that run.

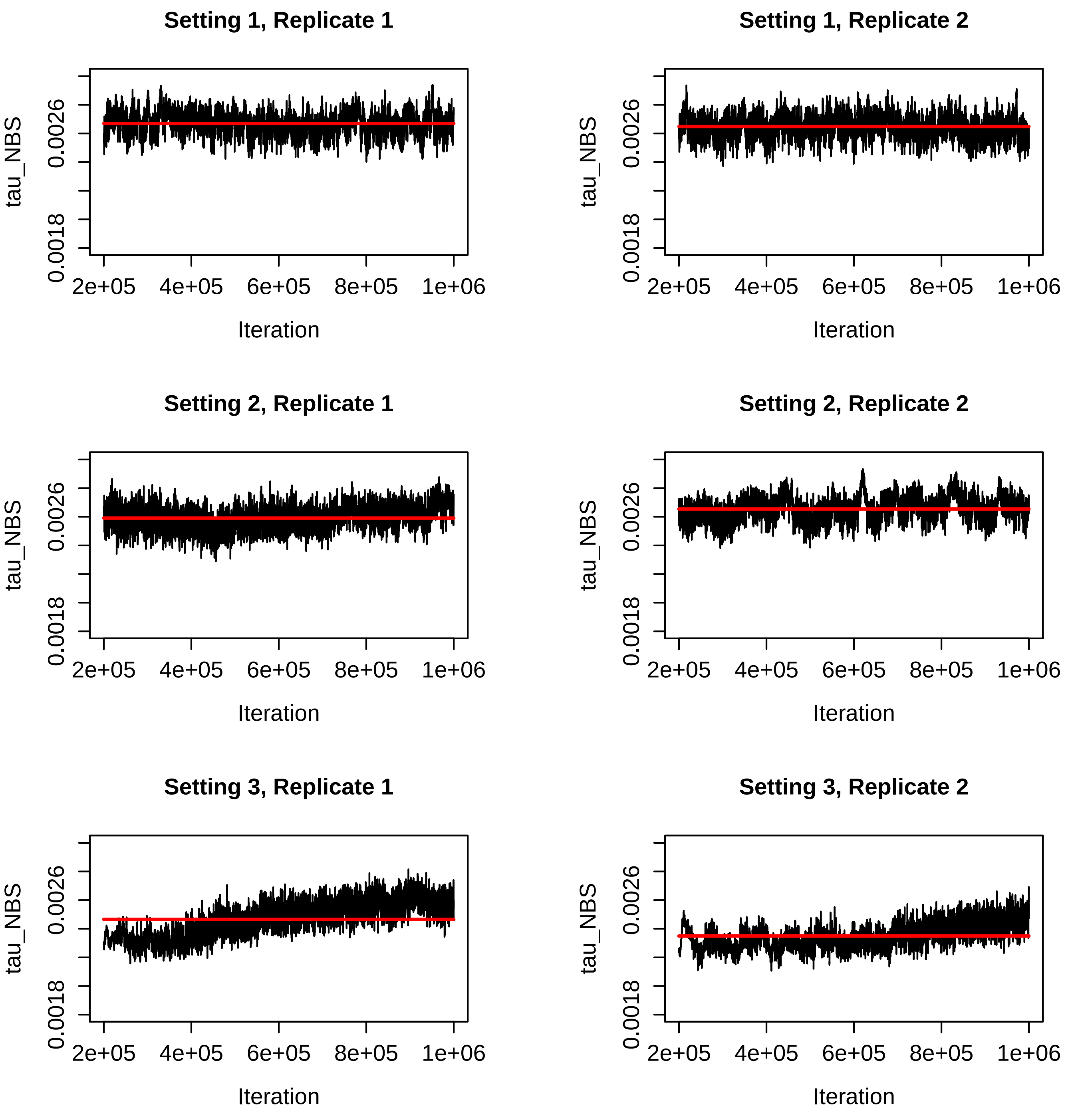


Figure S26: Trace plots for the $\tau_{NBS}$ parameter for the BPP runs for Settings 1-3 in Table S2. The red line is the mean of the sampled values for that run.

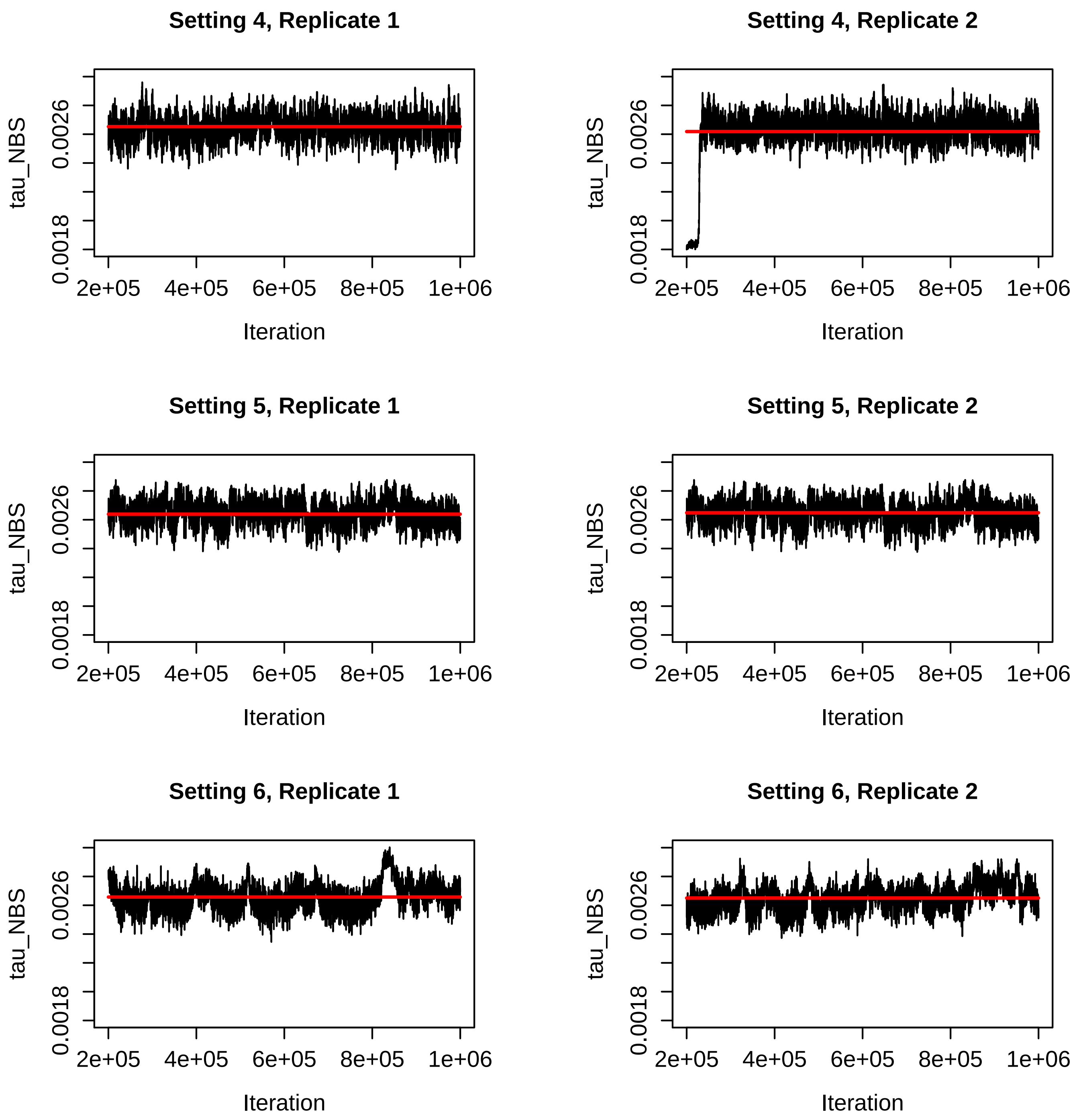


Figure S27: Trace plots for the $\tau_{NBS}$ parameter for the BPP runs for Settings 4-6 in Table S2. The red line is the mean of the sampled values for that run.

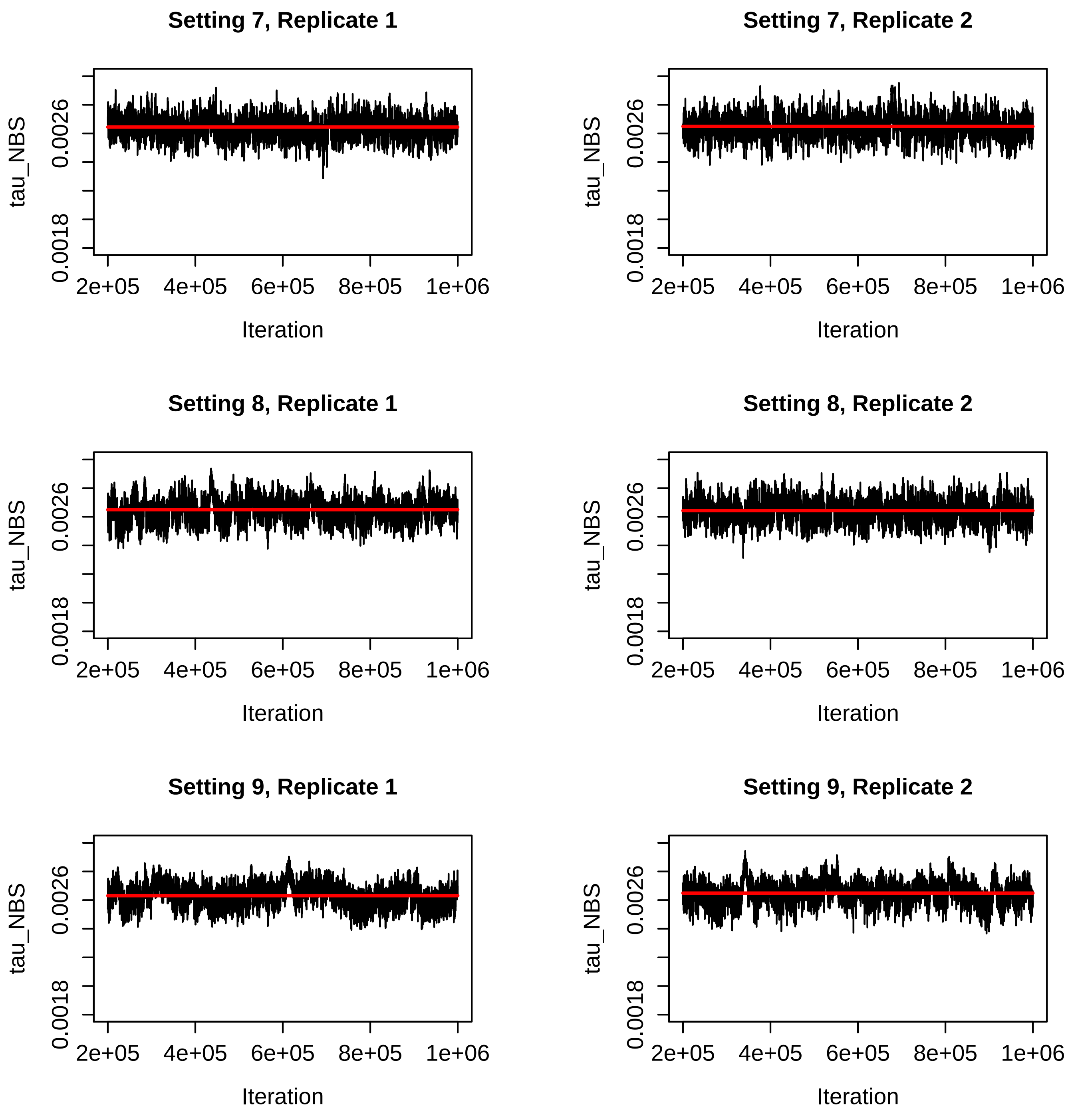


Figure S28: Trace plots for the $\tau_{NBS}$ parameter for the BPP runs for Settings 7-9 in Table S2. The red line is the mean of the sampled values for that run.

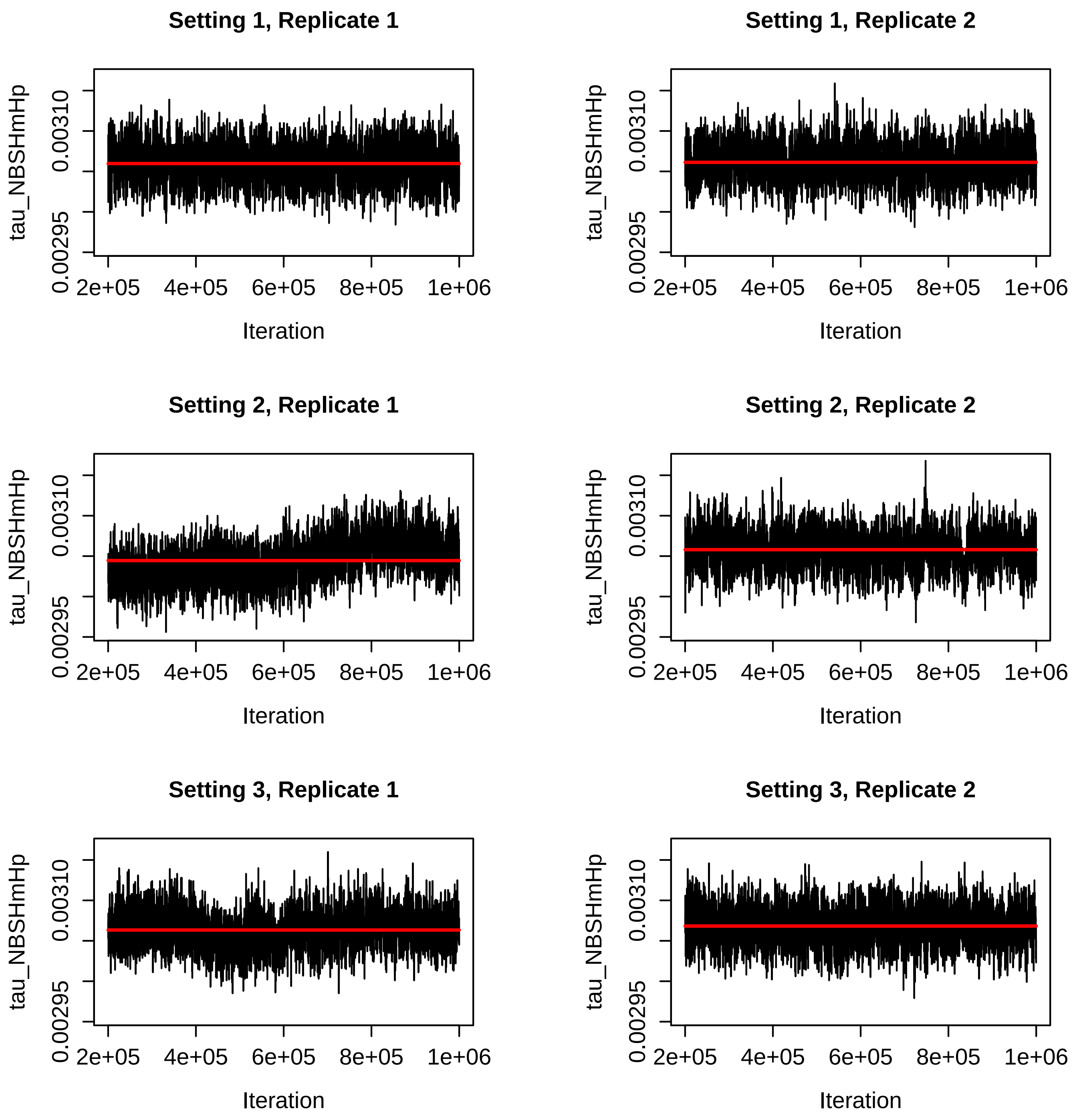


Figure S29: Trace plots for the $\tau_{NBSHmHp}$ parameter for the BPP runs for Settings 1-3 in Table S2. The red line is the mean of the sampled values for that run.

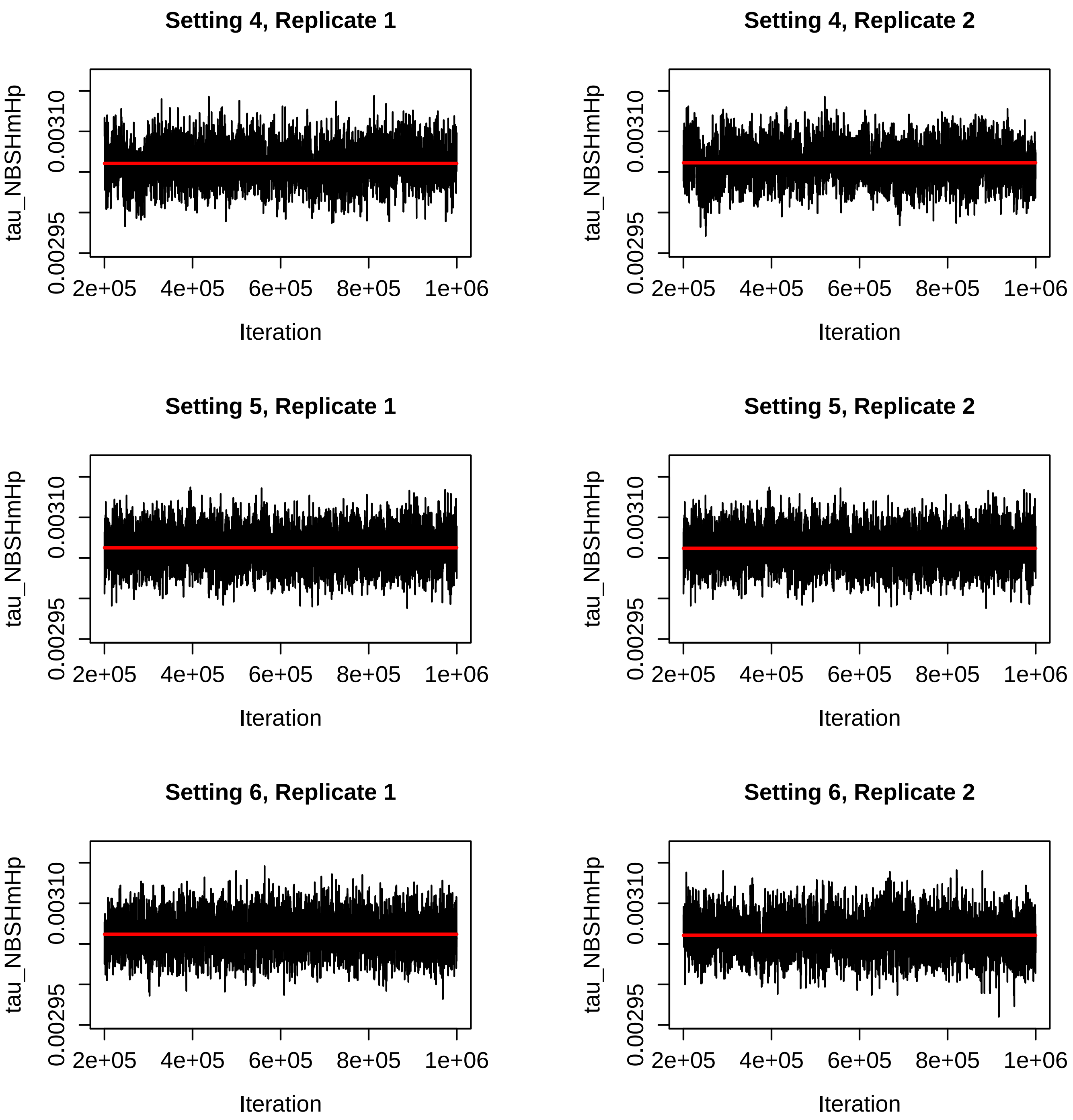


Figure S30: Trace plots for the $\tau_{NBSHmHp}$ parameter for the BPP runs for Settings 4-6 in Table S2. The red line is the mean of the sampled values for that run.

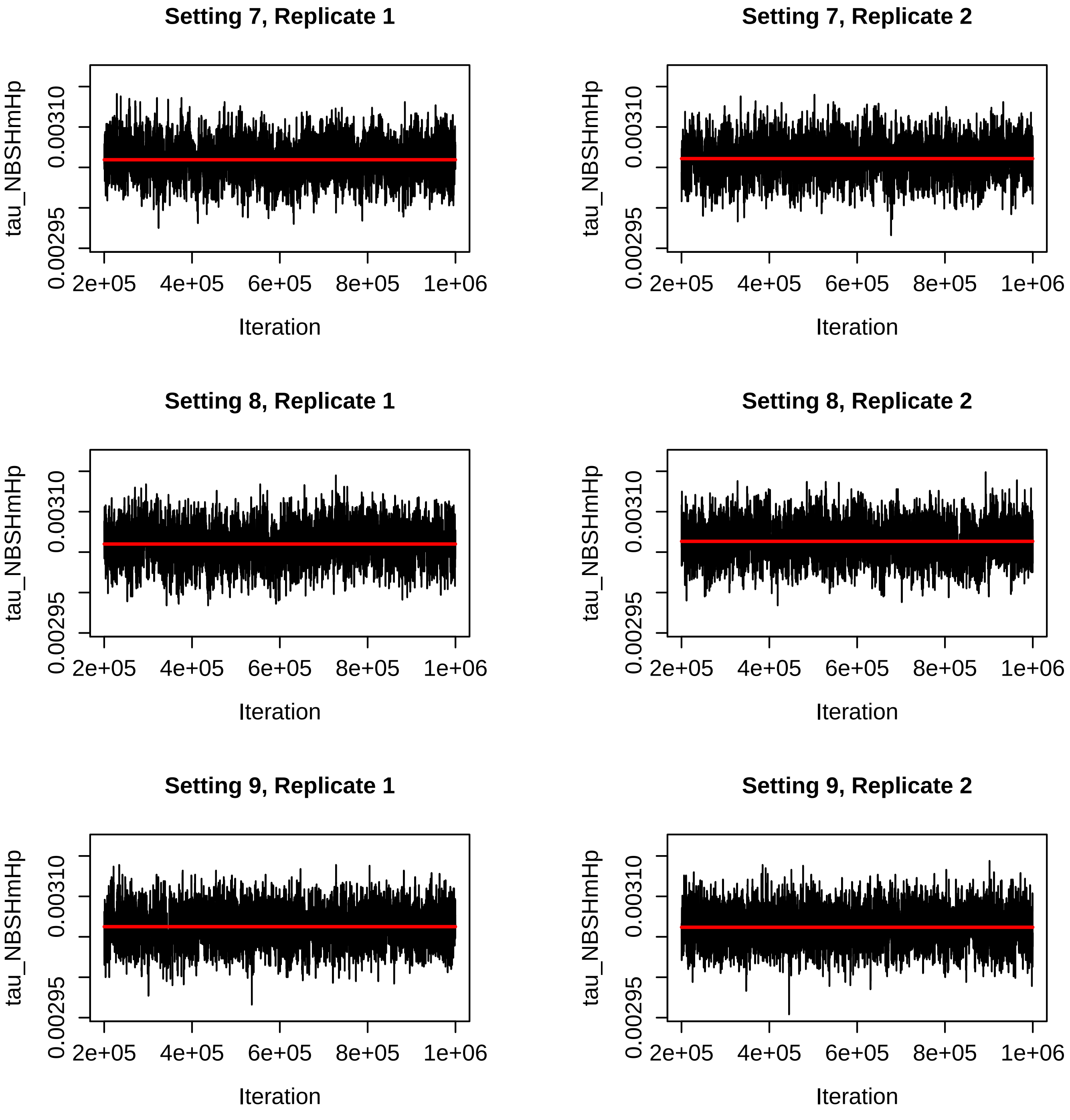


Figure S31: Trace plots for the $\tau_{NBSHmHp}$ parameter for the BPP runs for Settings 7-9 in Table S2. The red line is the mean of the sampled values for that run.

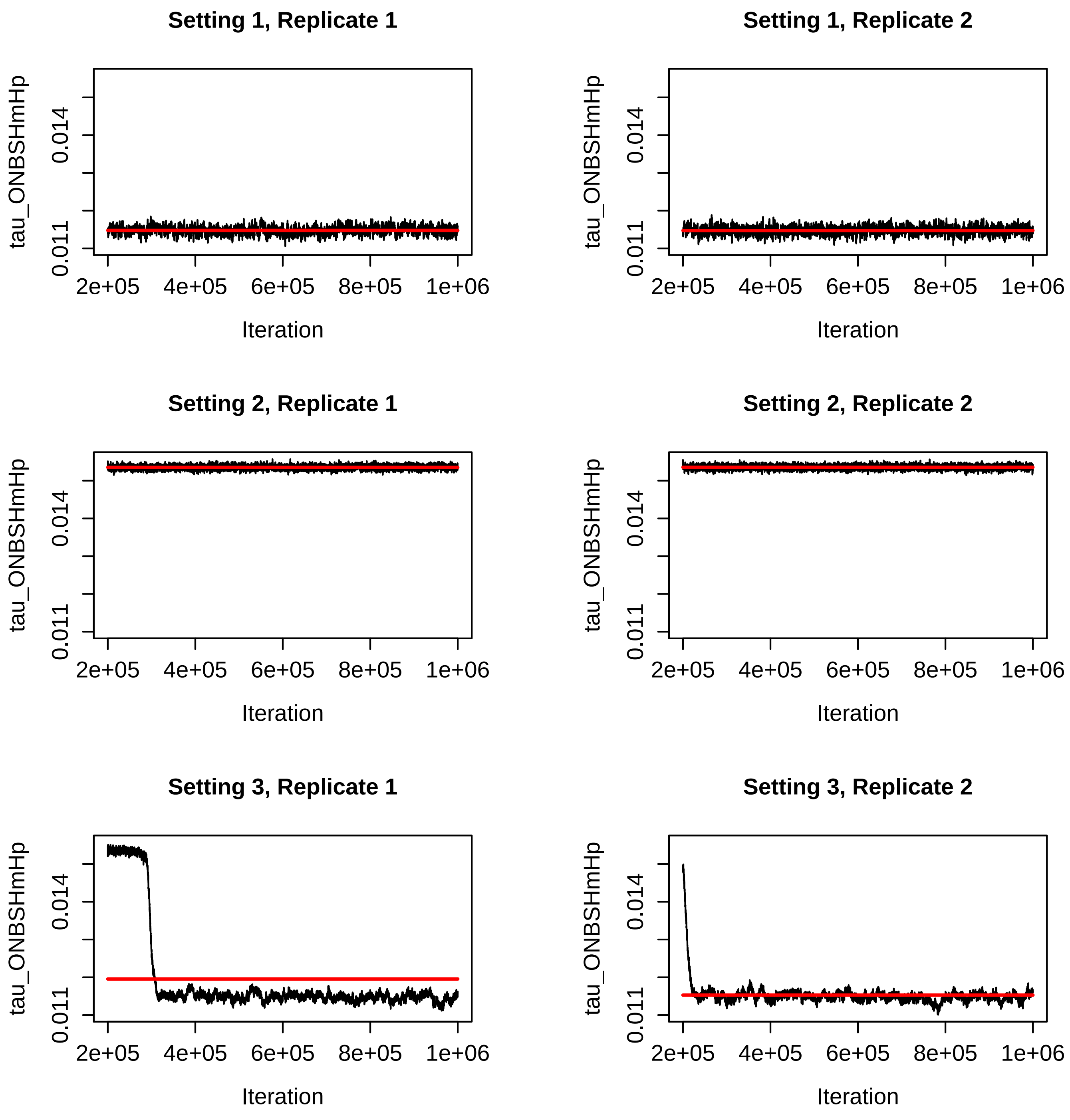


Figure S32: Trace plots for the $\tau_{ONBSHmHp}$ parameter for the BPP runs for Settings 1-3 in Table S2. The red line is the mean of the sampled values for that run.

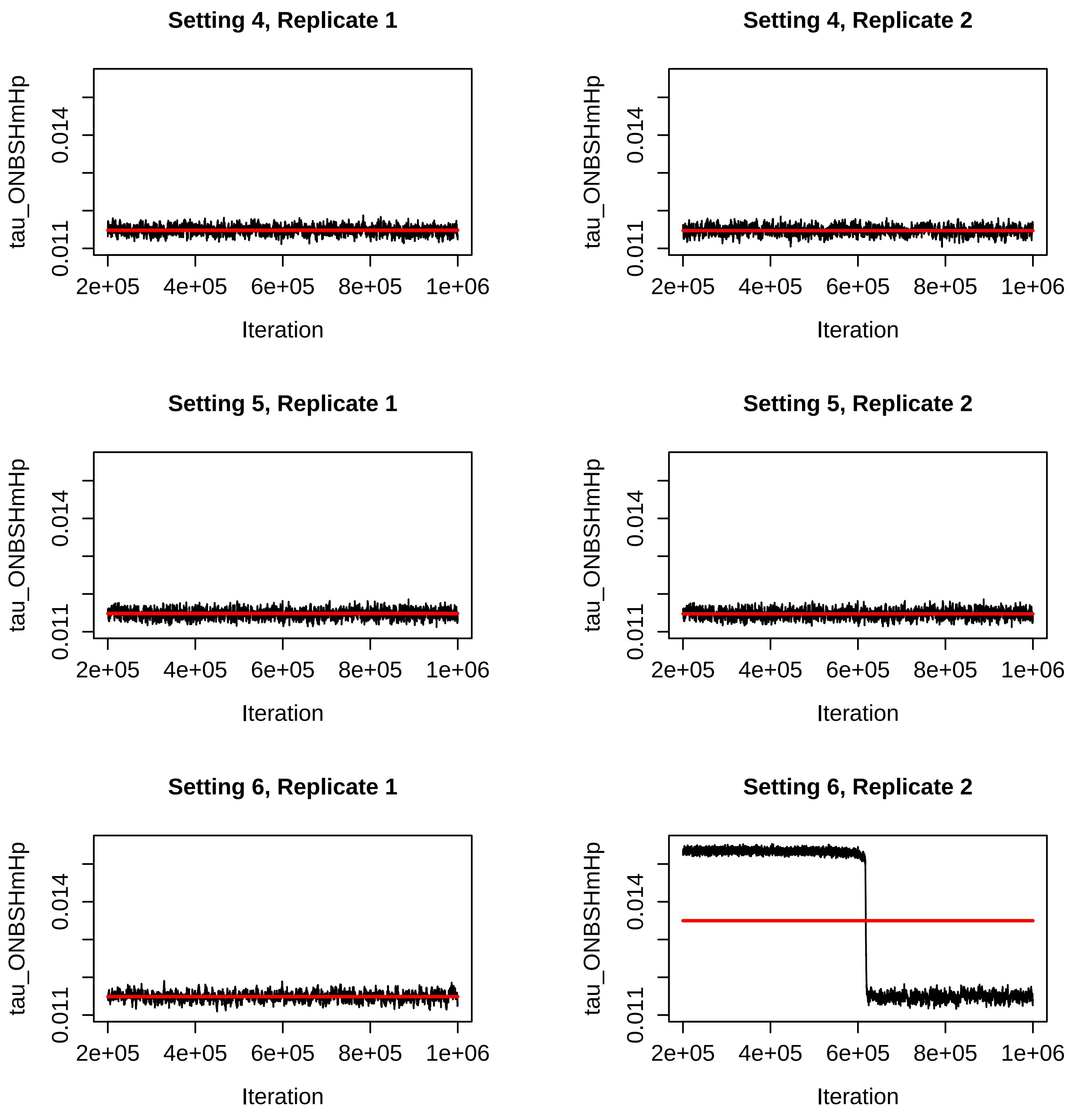


Figure S33: Trace plots for the $\tau_{ONBSHmHp}$ parameter for the BPP runs for Settings 4-6 in Table S2. The red line is the mean of the sampled values for that run.

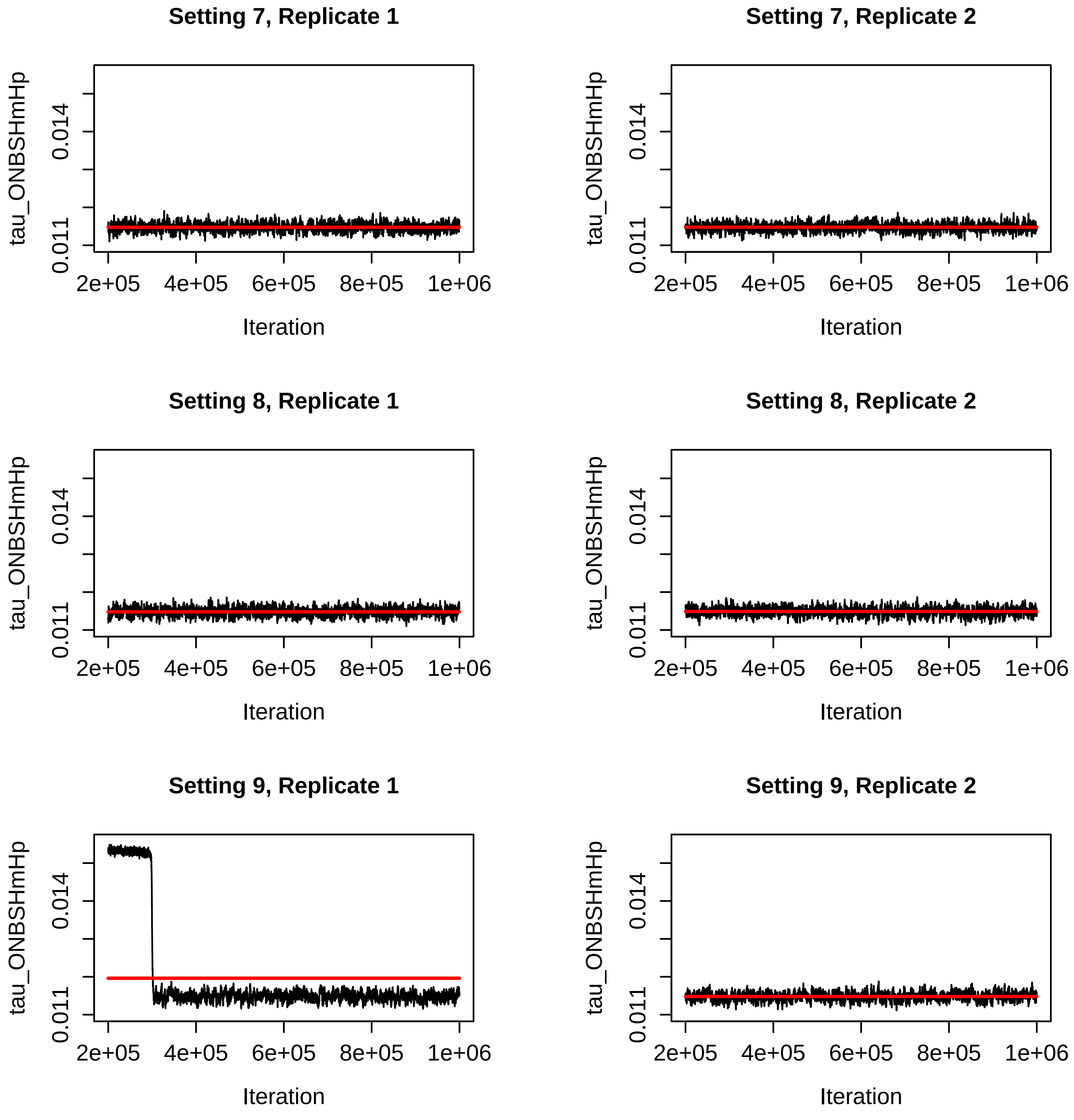


Figure S34: Trace plots for the $\tau_{ONBSHmHp}$ parameter for the BPP runs for Settings 7-9 in Table S2. The red line is the mean of the sampled values for that run.

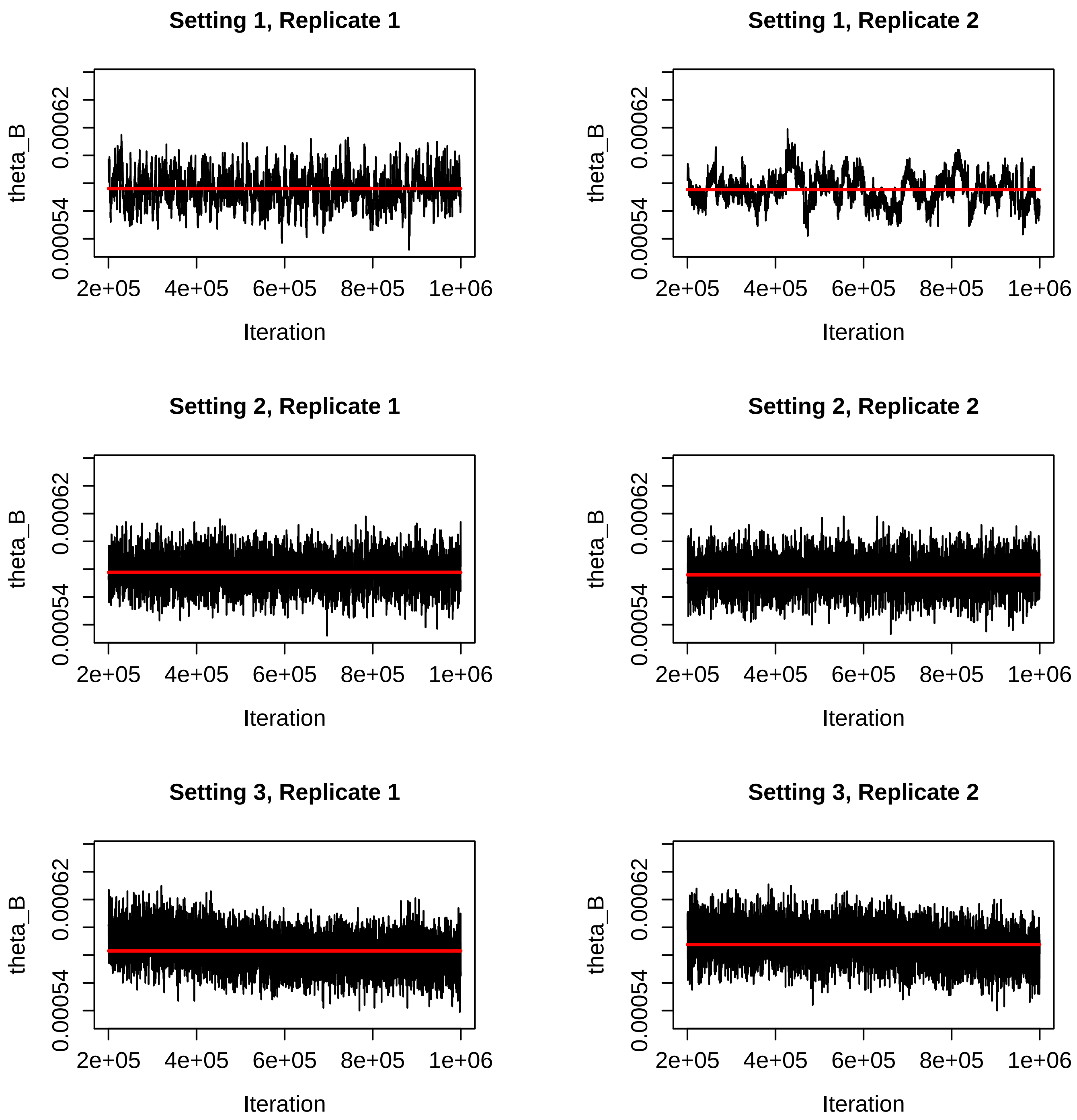


Figure S35: Trace plots for the $\theta_B$ parameter for the BPP runs for Settings 1-3 in Table S2. The red line is the mean of the sampled values for that run.

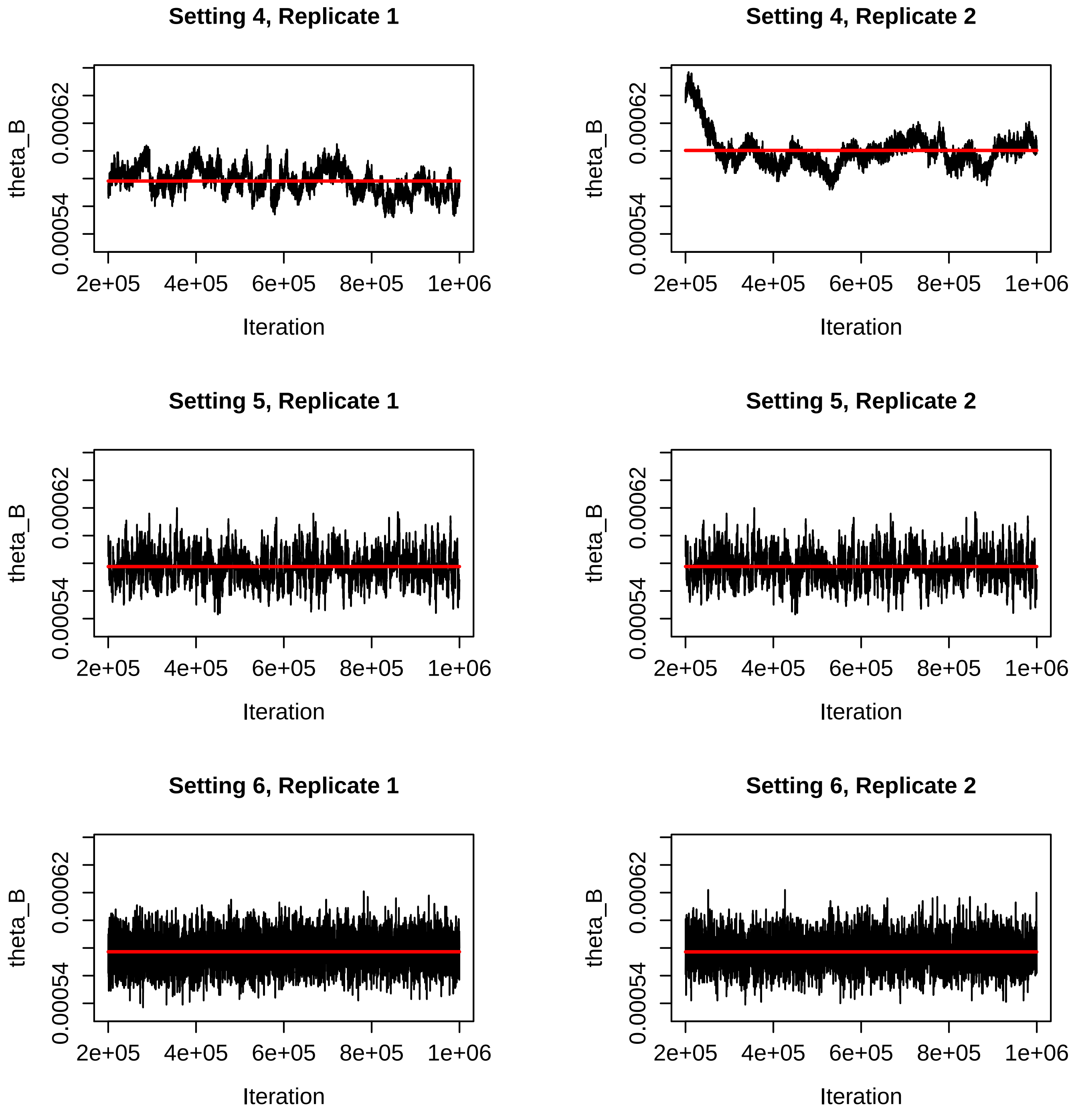


Figure S36: Trace plots for the $\theta_B$ parameter for the BPP runs for Settings 4-6 in Table S2. The red line is the mean of the sampled values for that run.

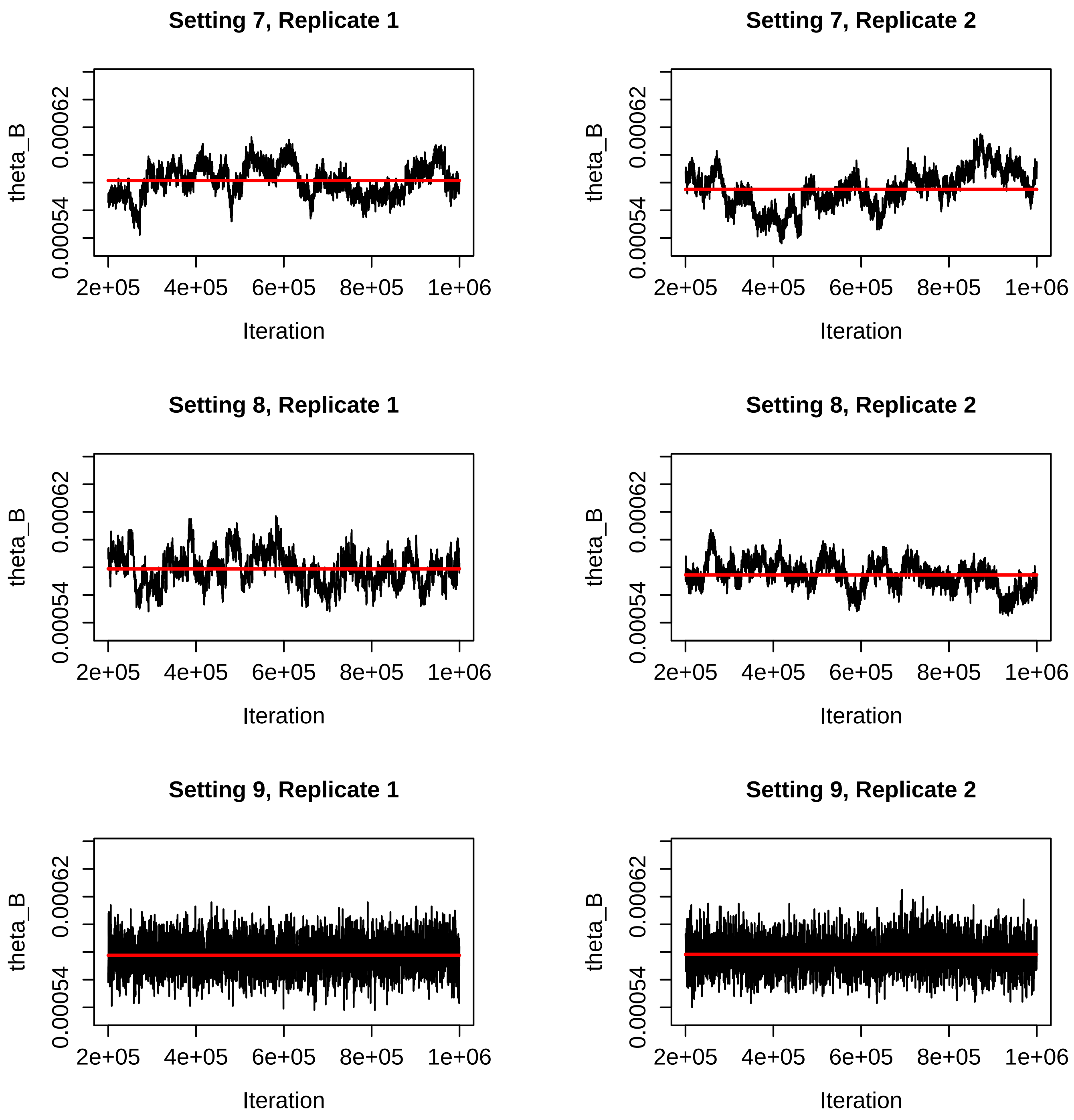


Figure S37: Trace plots for the $\theta_B$ parameter for the BPP runs for Settings 7-9 in Table S2. The red line is the mean of the sampled values for that run.

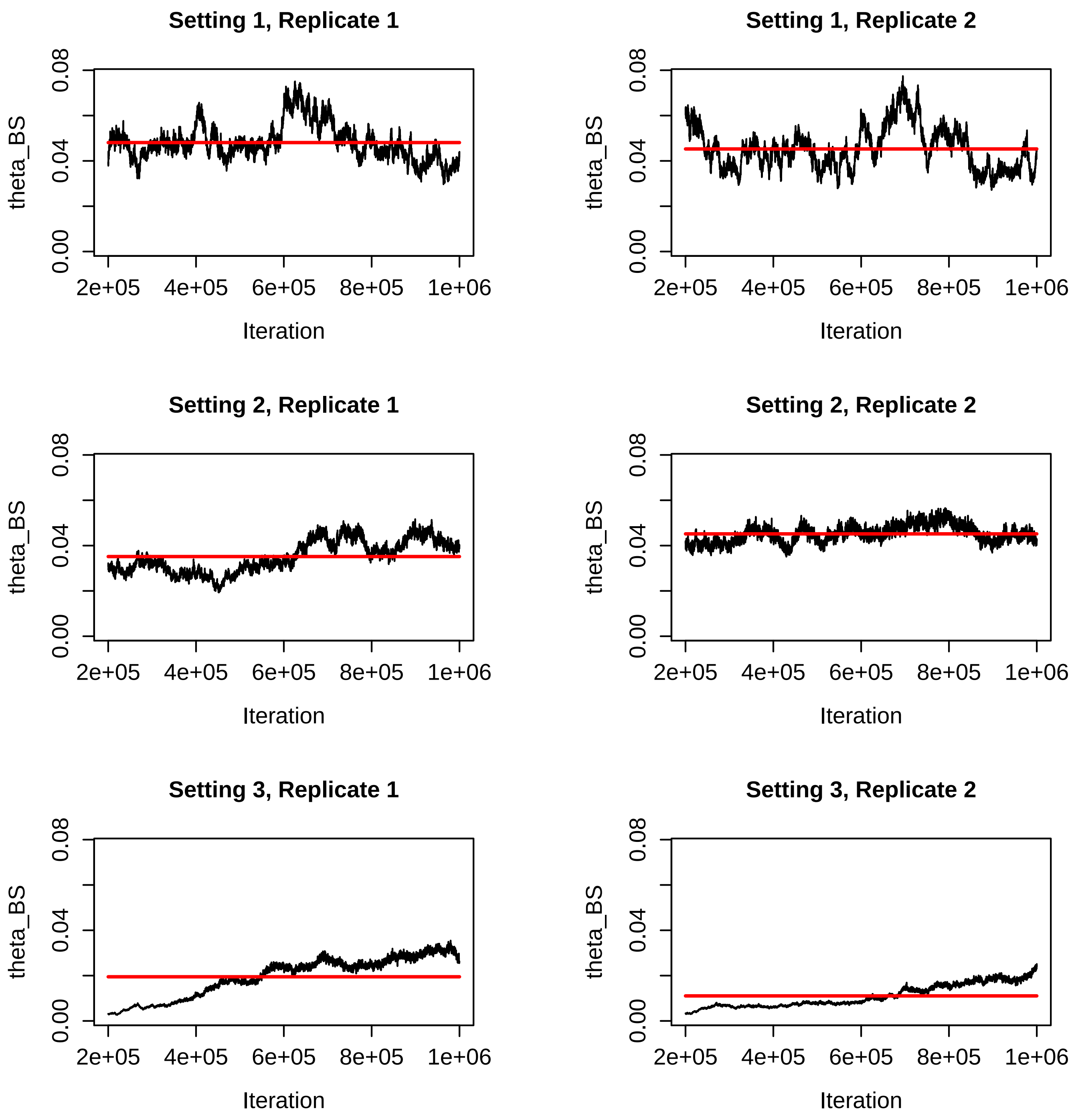


Figure S38: Trace plots for the $\theta_{BS}$ parameter for the BPP runs for Settings 1-3 in Table S2. The red line is the mean of the sampled values for that run. Note the difference in the y-axis values for Setting 3, Replicate 1.

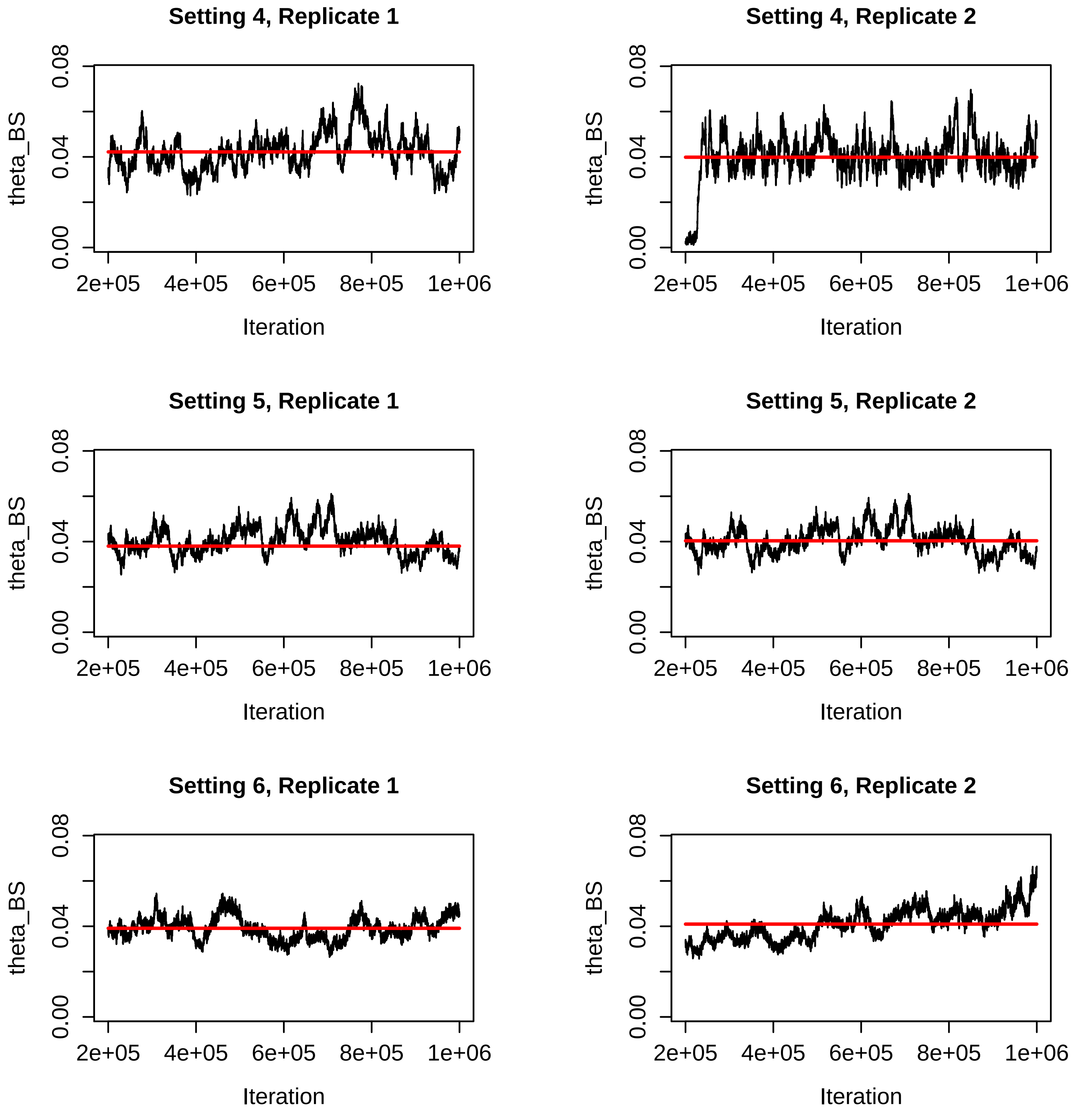


Figure S39: Trace plots for the $\theta_{BS}$ parameter for the BPP runs for Settings 4-6 in Table S2. The red line is the mean of the sampled values for that run.

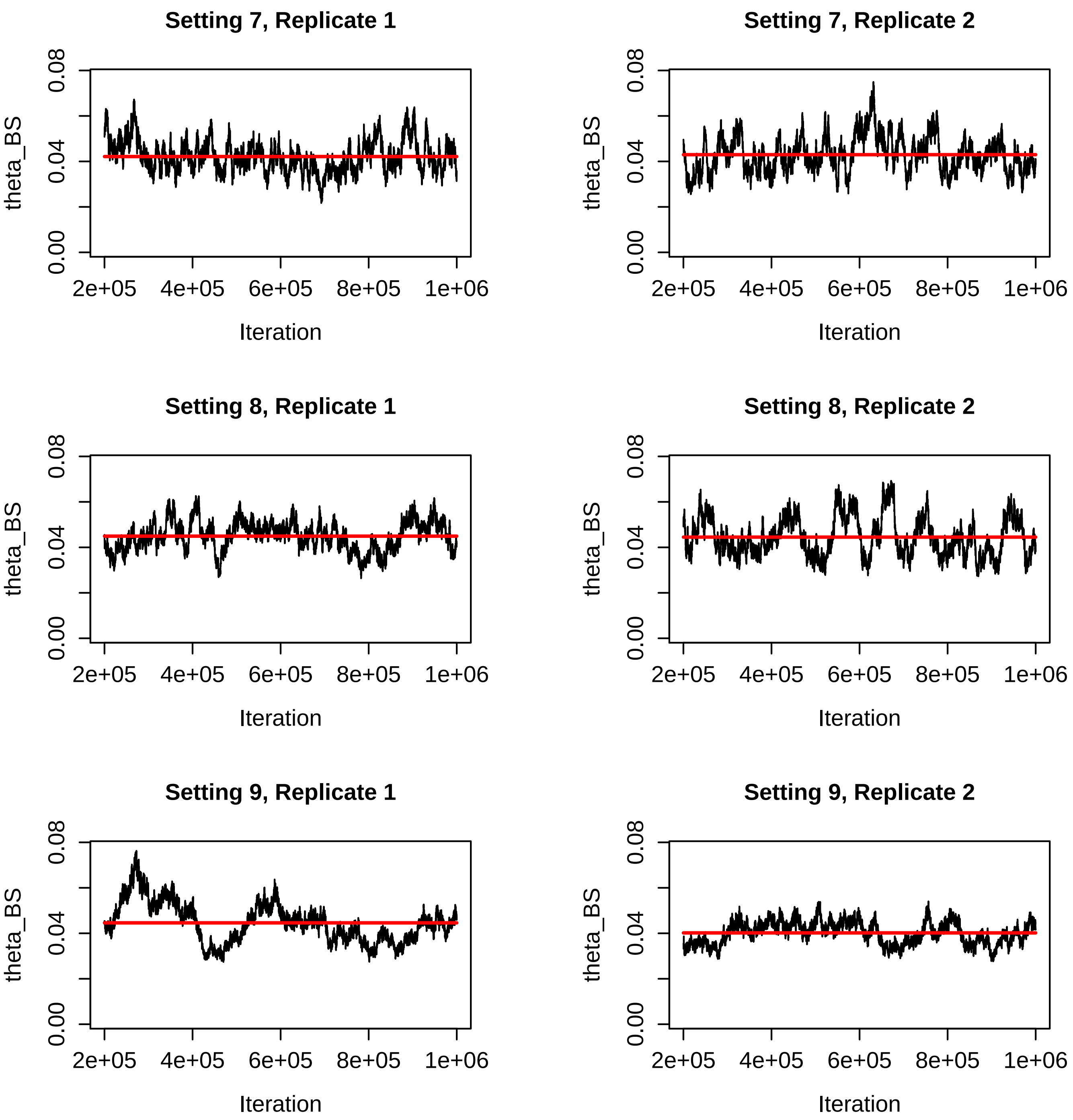


Figure S40: Trace plots for the $\theta_{BS}$ parameter for the BPP runs for Settings 7-9 in Table S2. The red line is the mean of the sampled values for that run.

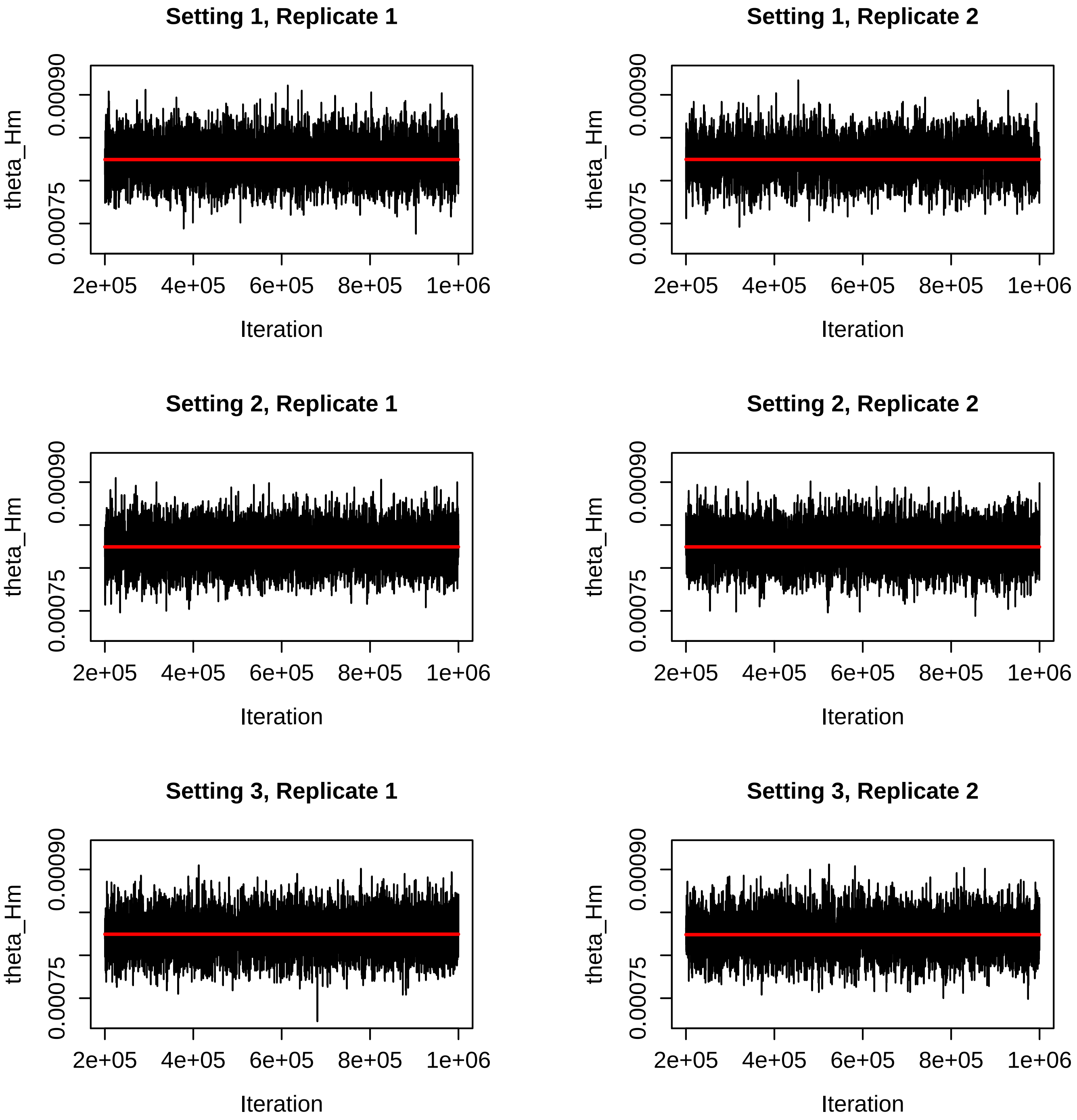


Figure S41: Trace plots for the $\theta_{Hm}$ parameter for the BPP runs for Settings 1-3 in Table S2. The red line is the mean of the sampled values for that run.

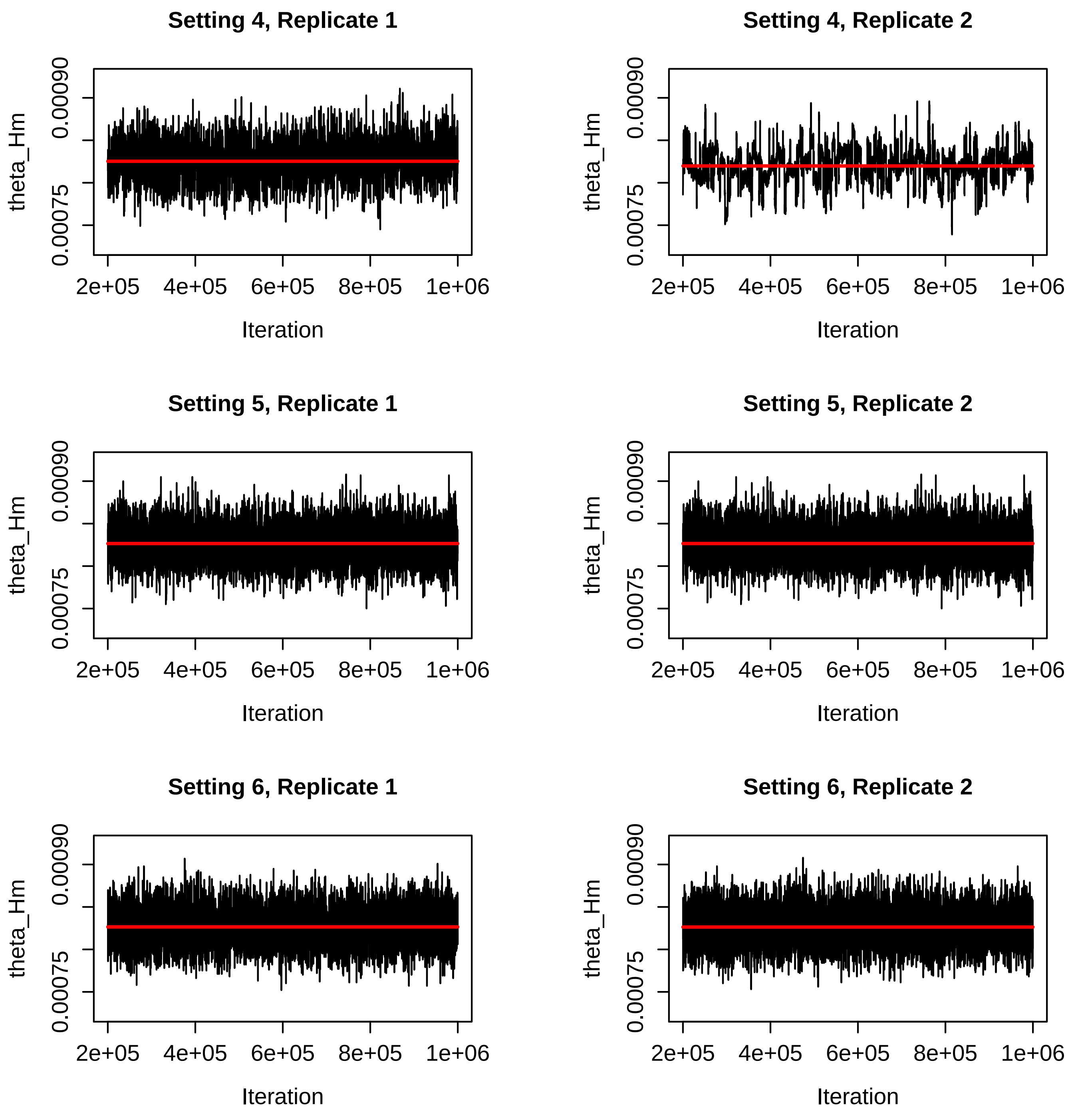


Figure S42: Trace plots for the $\theta_{Hm}$ parameter for the BPP runs for Settings 4-6 in Table S2. The red line is the mean of the sampled values for that run.

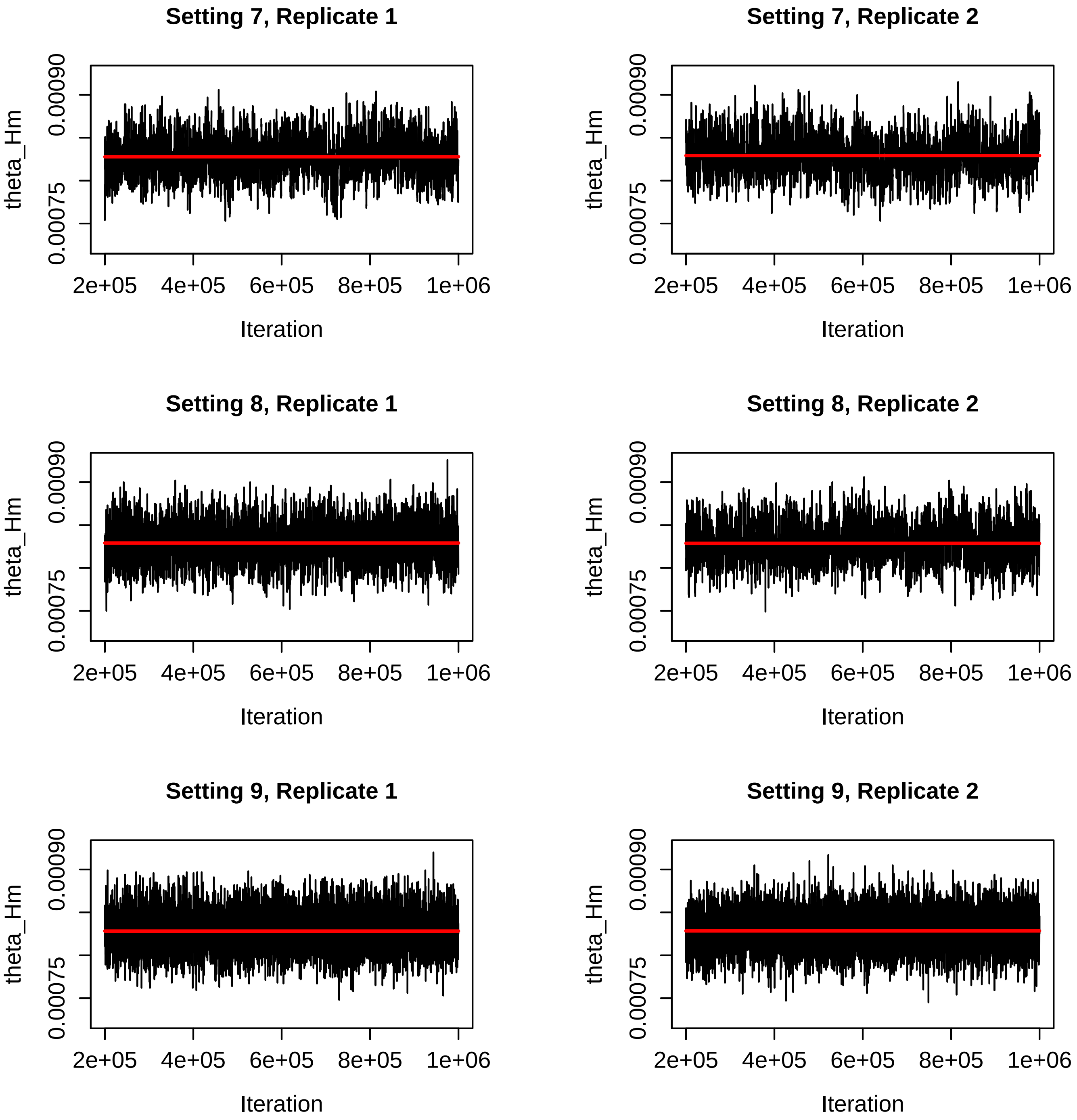


Figure S43: Trace plots for the $\theta_{Hm}$ parameter for the BPP runs for Settings 7-9 in Table S2. The red line is the mean of the sampled values for that run.

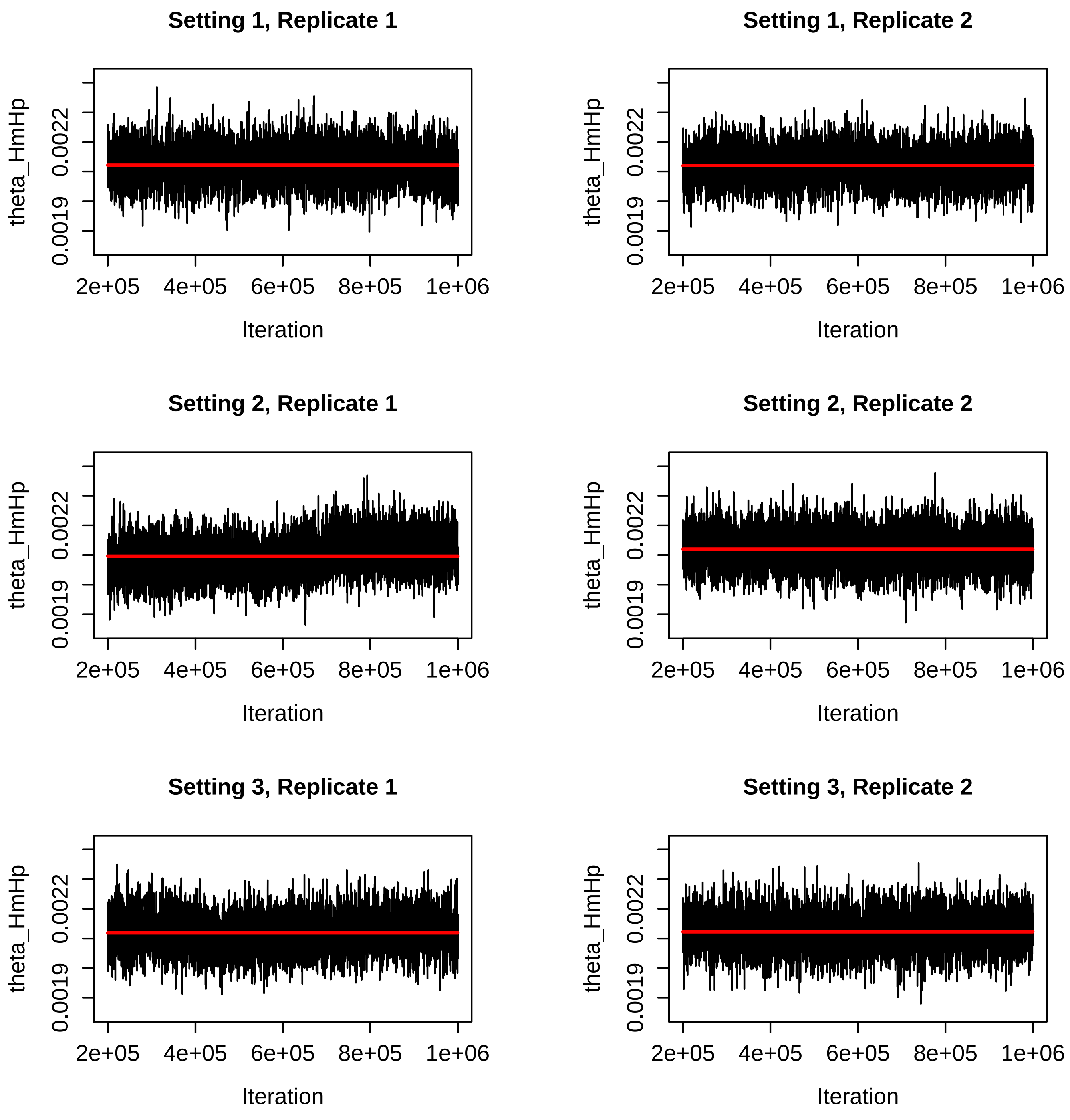


Figure S44: Trace plots for the $\theta_{HmHp}$ parameter for the BPP runs for Settings 1-3 in Table S2. The red line is the mean of the sampled values for that run.

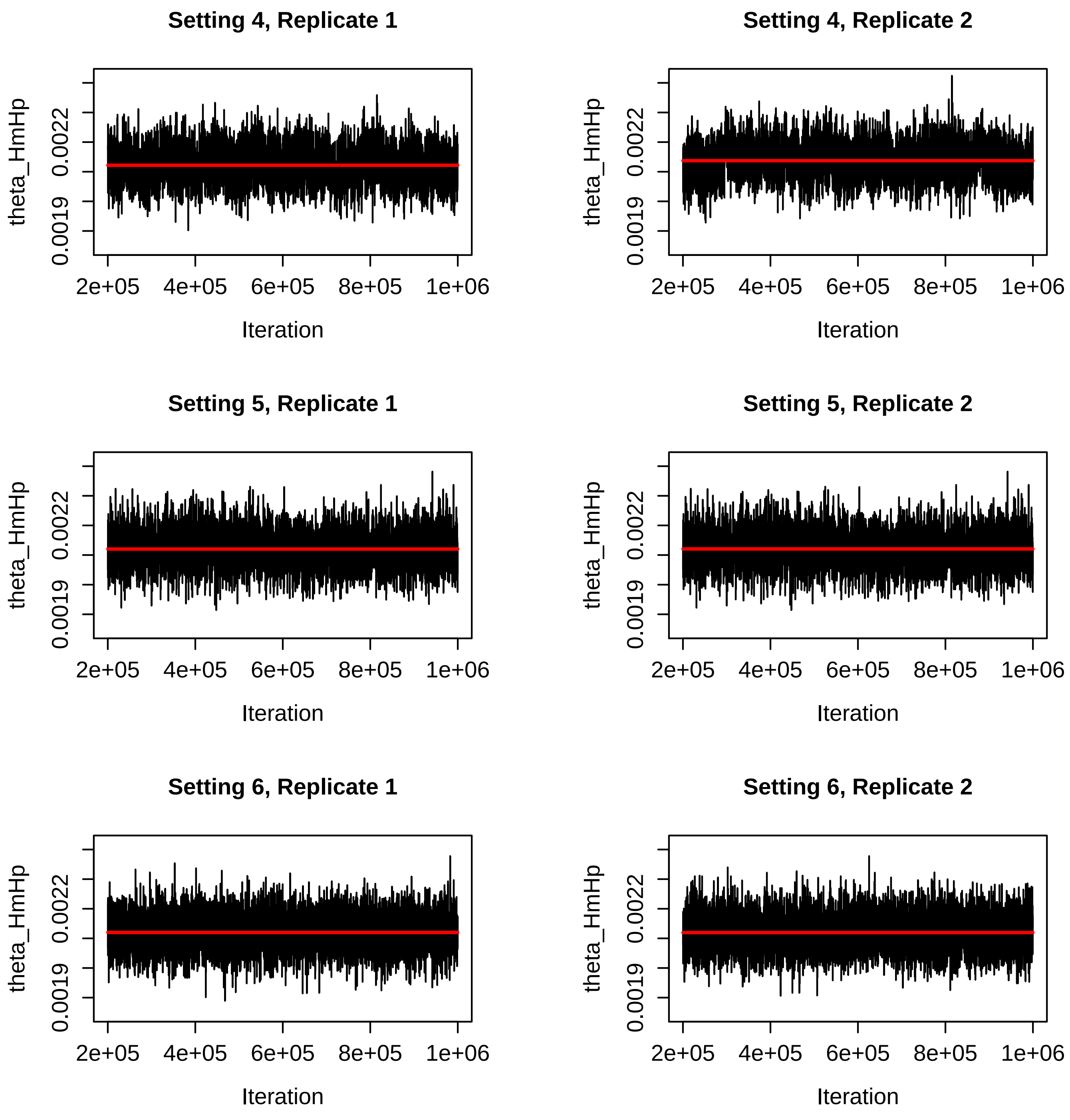


Figure S45: Trace plots for the $\theta_{HmHp}$ parameter for the BPP runs for Settings 4-6 in Table S2. The red line is the mean of the sampled values for that run.

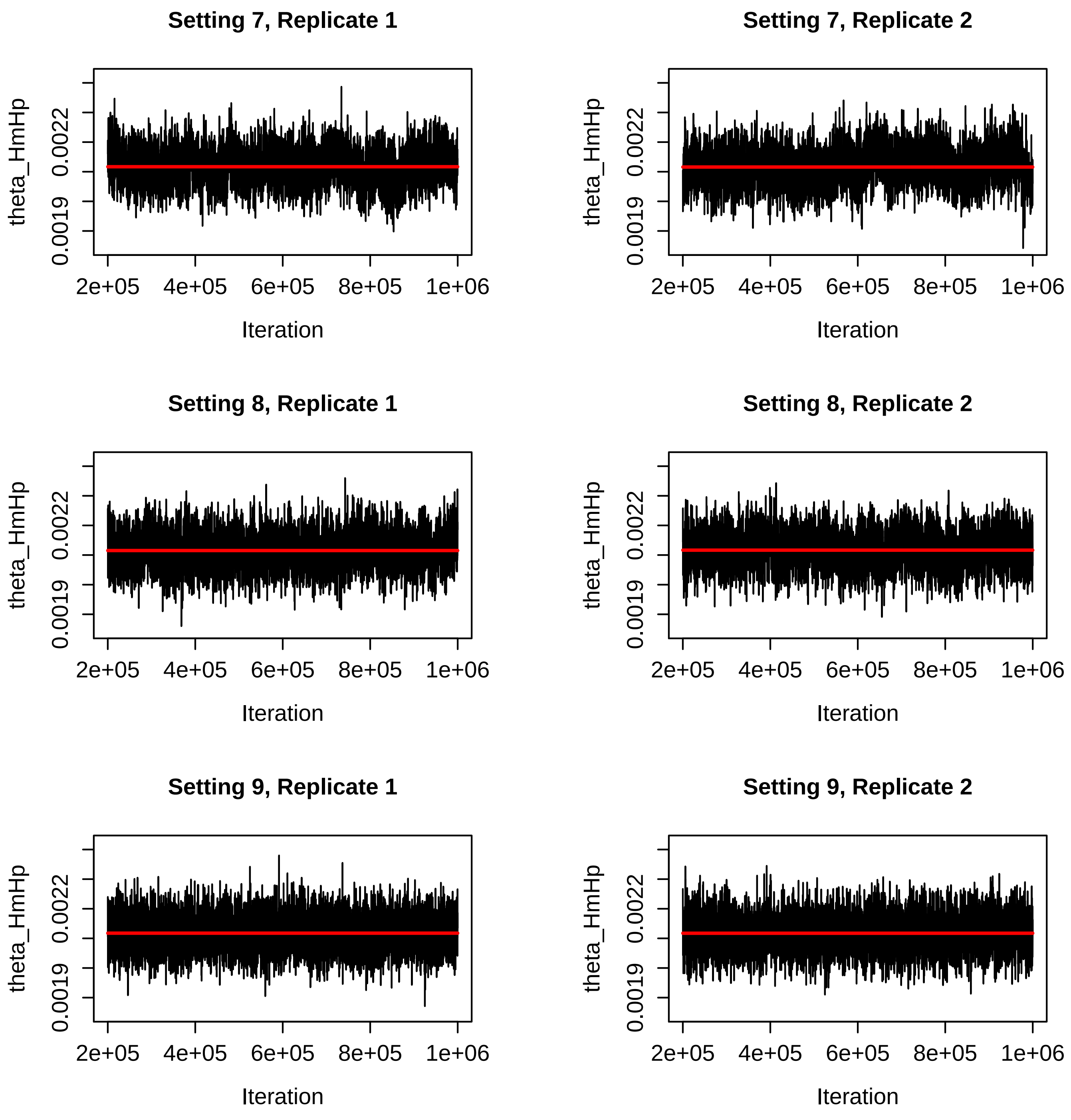


Figure S46: Trace plots for the $\theta_{HmHp}$ parameter for the BPP runs for Settings 7-9 in Table S2. The red line is the mean of the sampled values for that run.

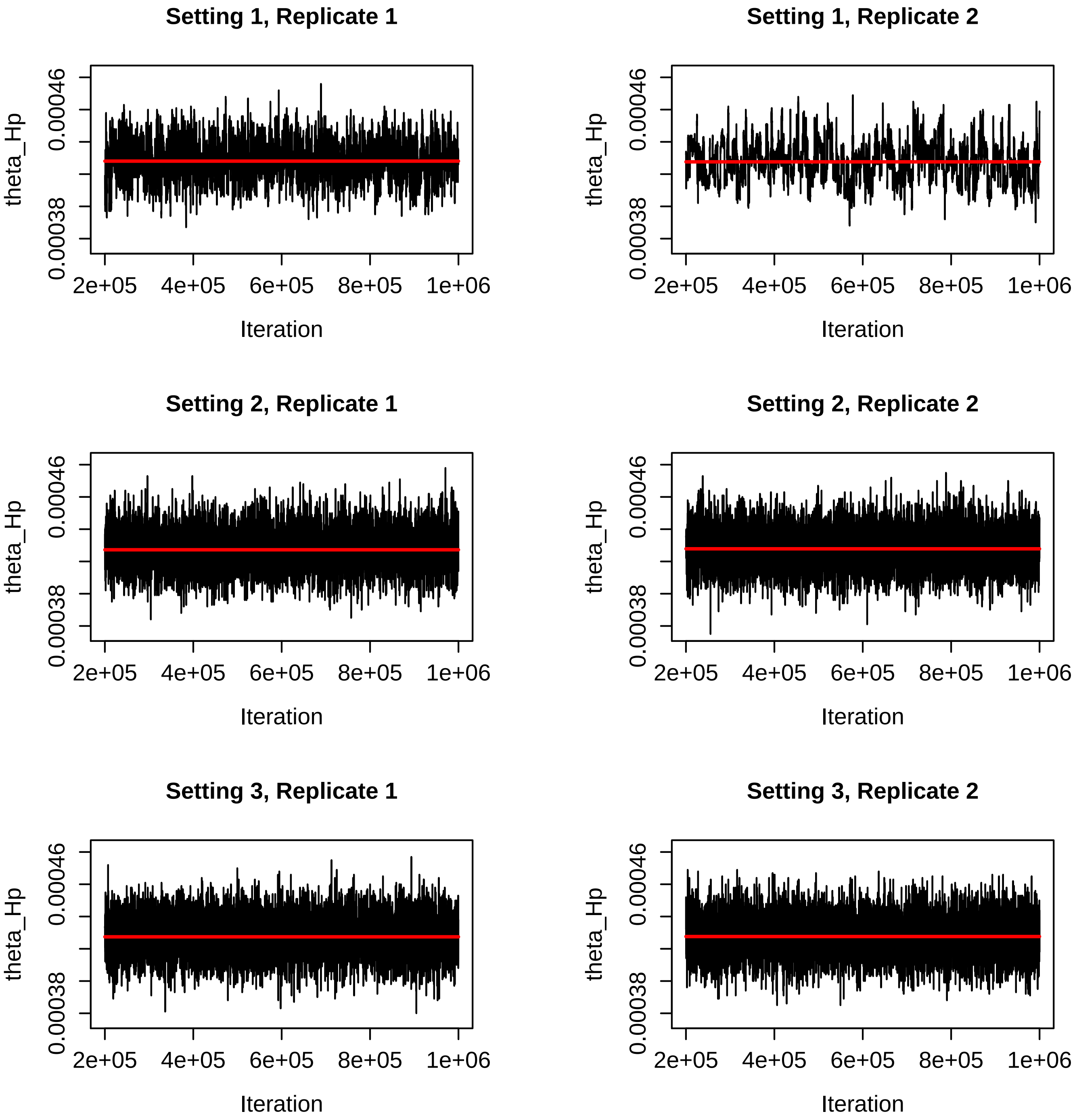


Figure S47: Trace plots for the $\theta_{Hp}$ parameter for the BPP runs for Settings 1-3 in Table S2. The red line is the mean of the sampled values for that run.

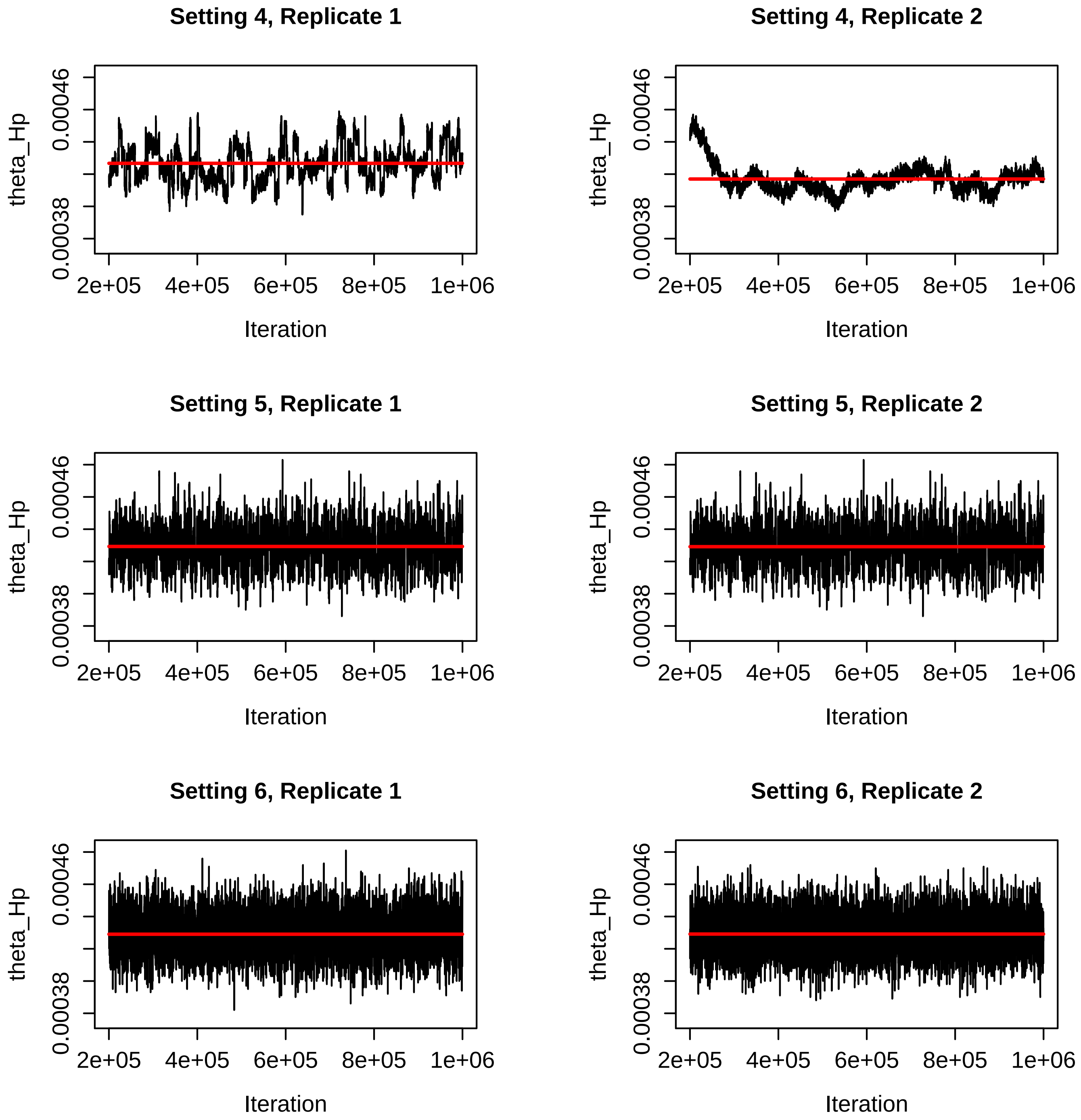


Figure S48: Trace plots for the $\theta_{Hp}$ parameter for the BPP runs for Settings 4-6 in Table S2. The red line is the mean of the sampled values for that run.

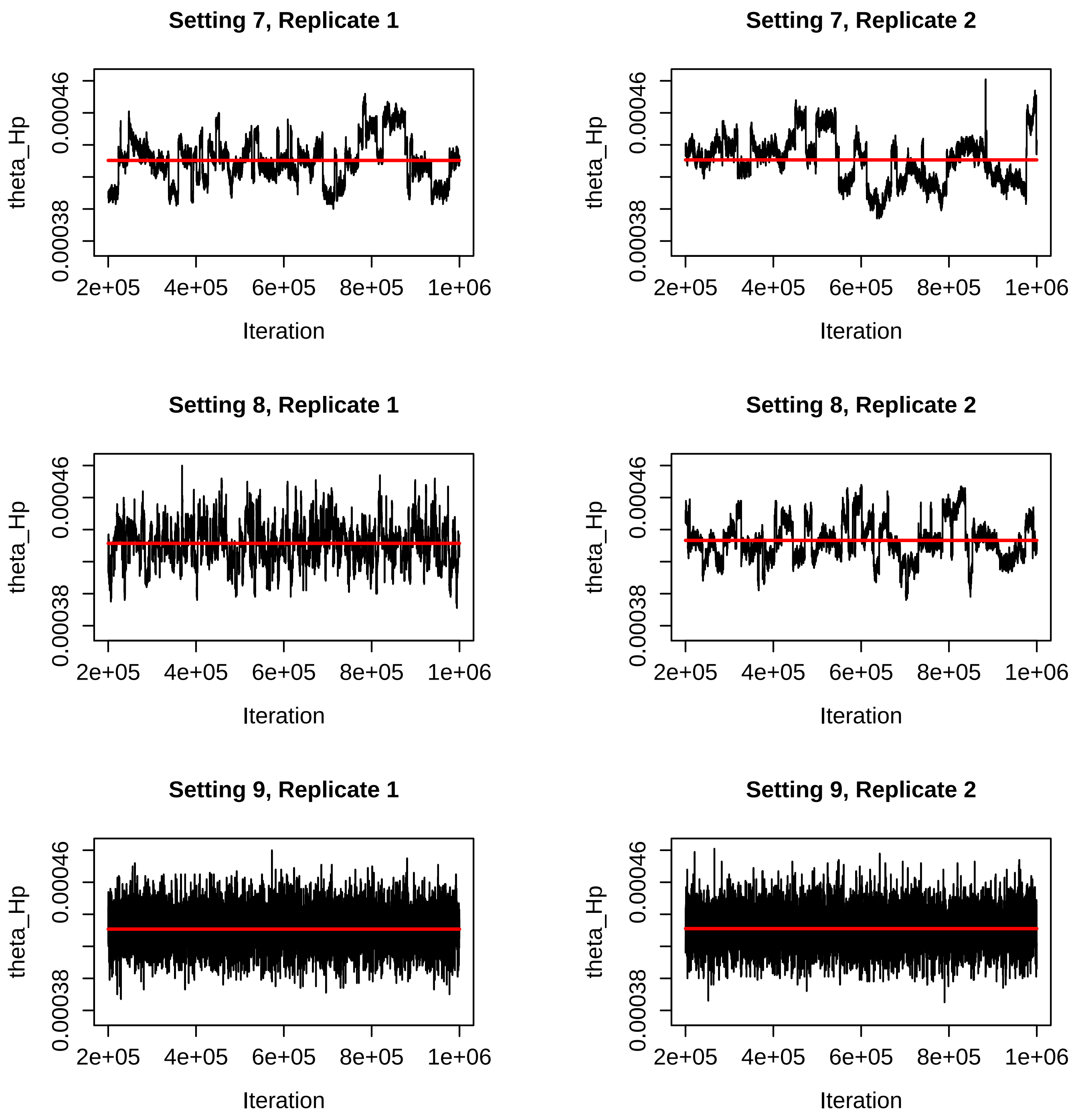


Figure S49: Trace plots for the $\theta_{Hp}$ parameter for the BPP runs for Settings 7-9 in Table S2. The red line is the mean of the sampled values for that run.

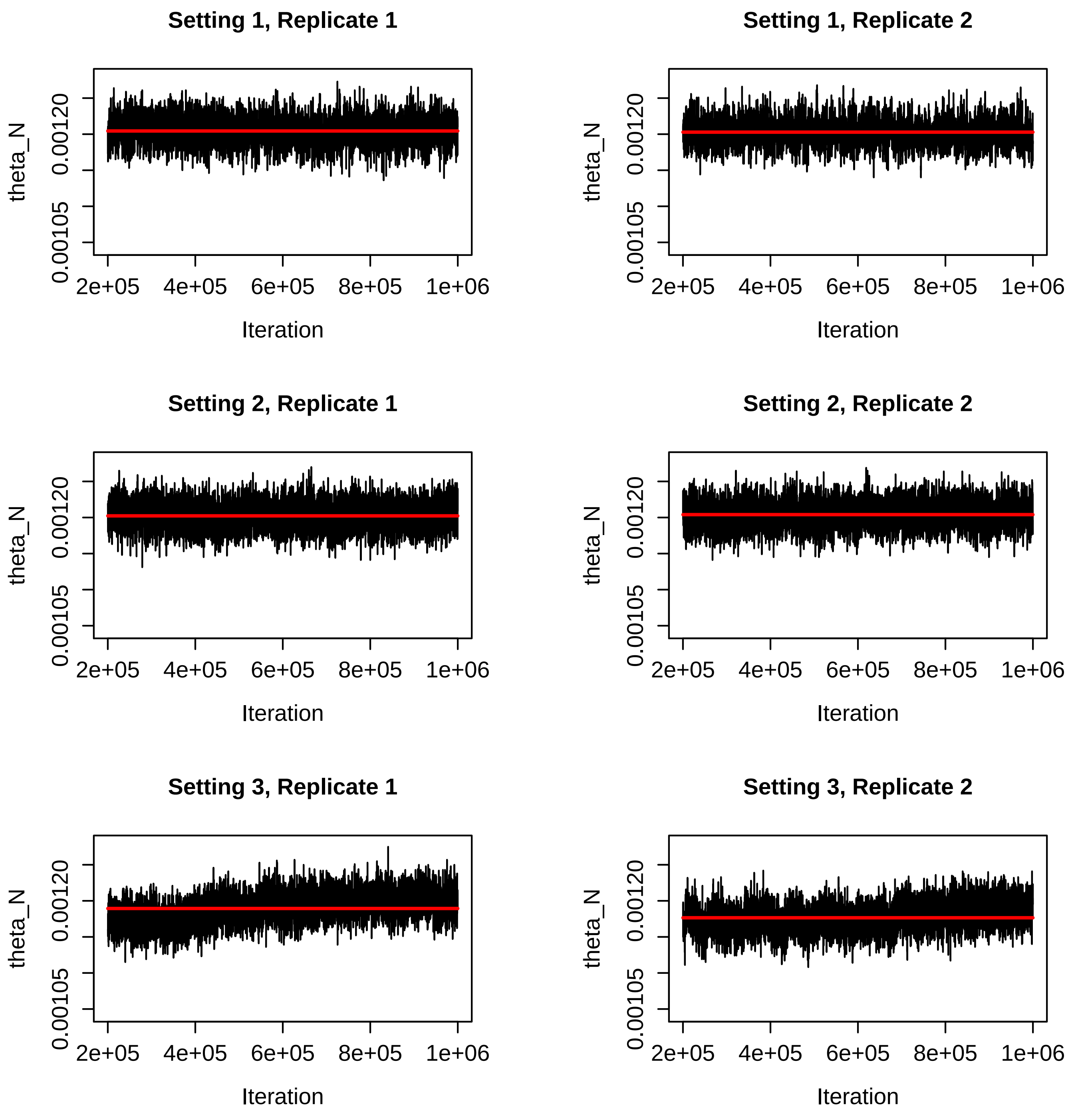


Figure S50: Trace plots for the $\theta_N$ parameter for the BPP runs for Settings 1-3 in Table S2. The red line is the mean of the sampled values for that run.

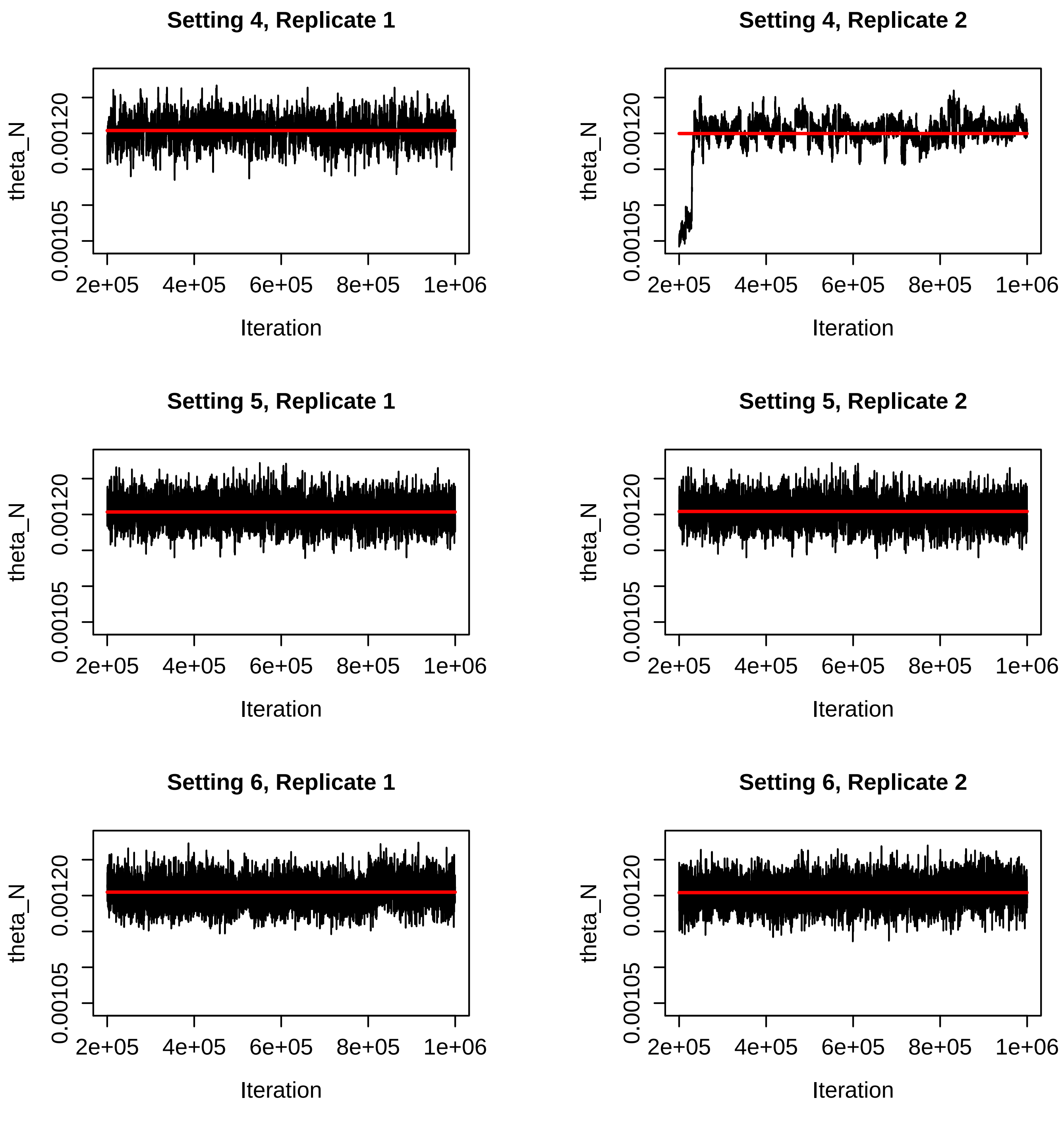


Figure S51: Trace plots for the $\theta_N$ parameter for the BPP runs for Settings 4-6 in Table S2. The red line is the mean of the sampled values for that run.

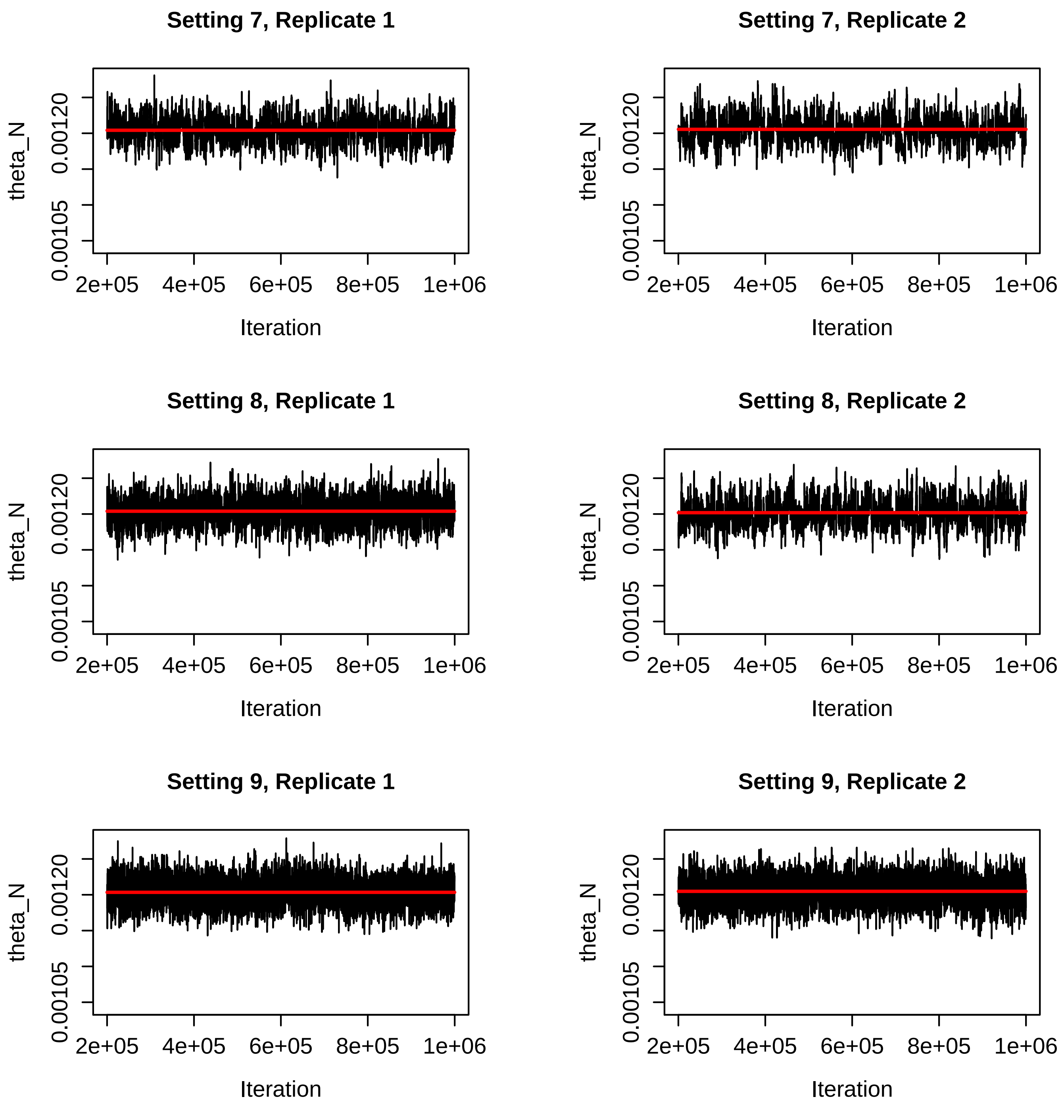


Figure S52: Trace plots for the $\theta_N$ parameter for the BPP runs for Settings 7-9 in Table S2. The red line is the mean of the sampled values for that run.

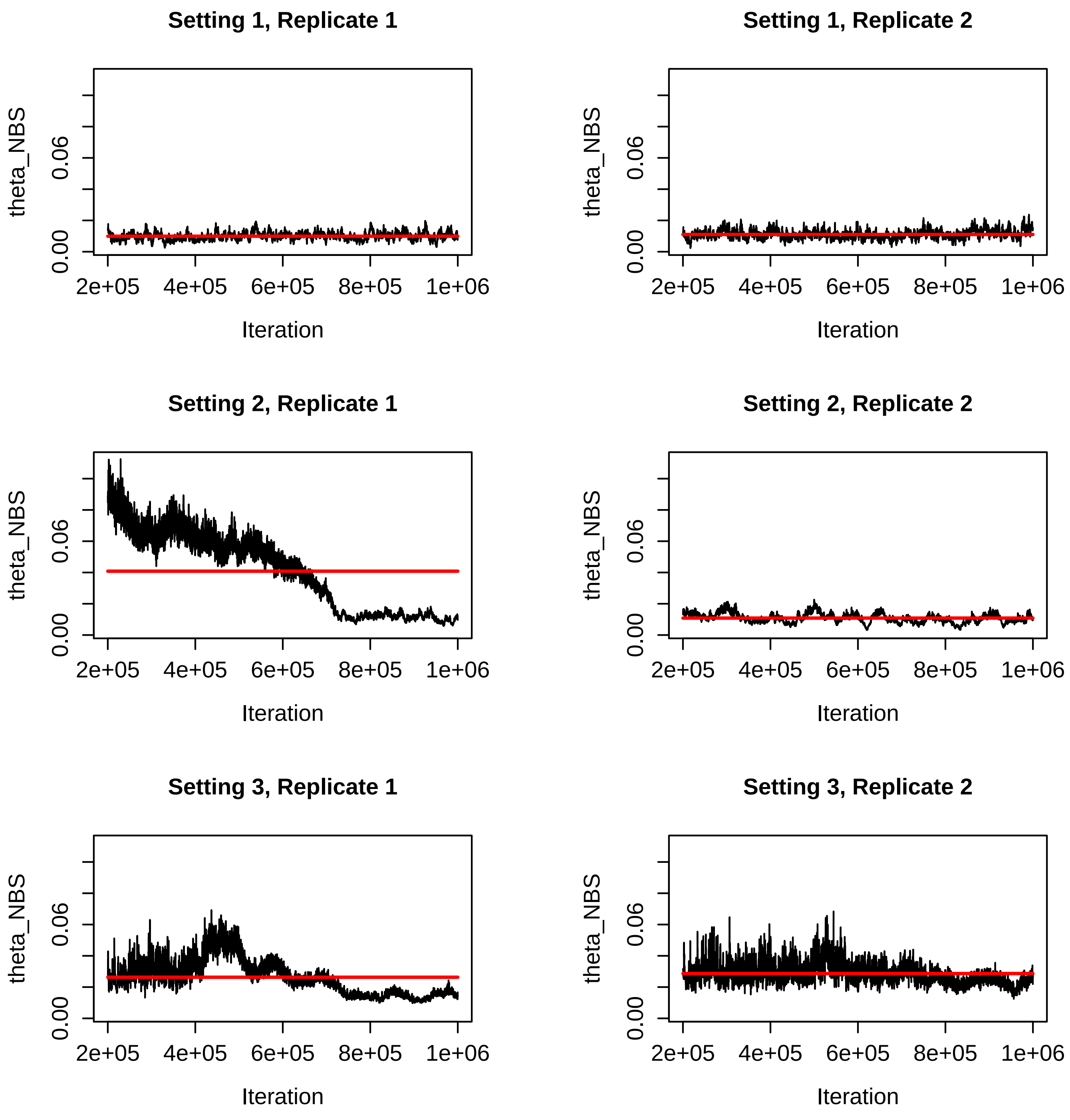


Figure S53: Trace plots for the $\theta_{NBS}$ parameter for the BPP runs for Settings 1-3 in Table S2. The red line is the mean of the sampled values for that run.

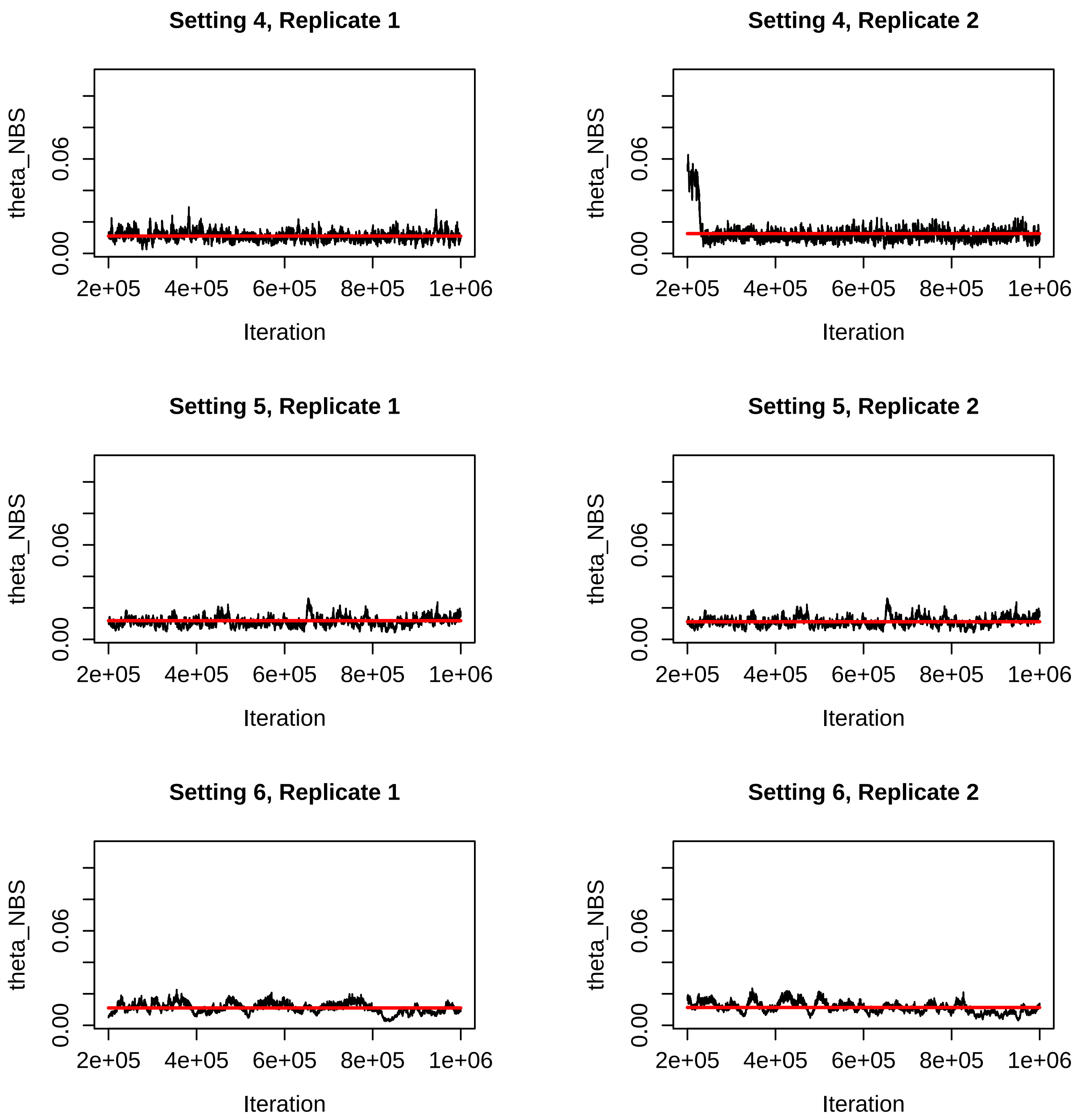


Figure S54: Trace plots for the $\theta_{NBS}$ parameter for the BPP runs for Settings 4-6 in Table S2. The red line is the mean of the sampled values for that run. Note the difference in the y-axis scale for Setting 6, Replicate 1.

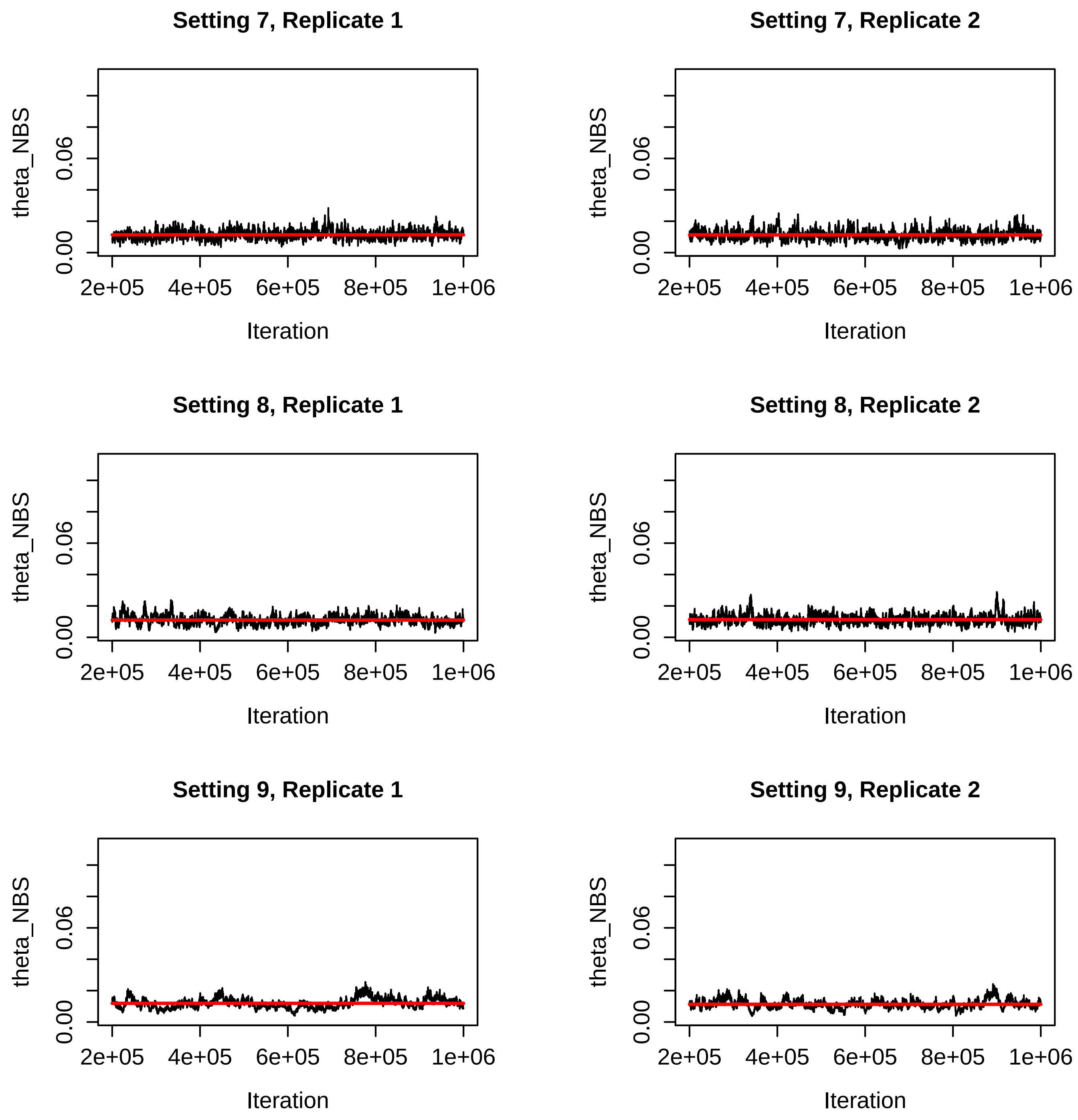


Figure S55: Trace plots for the $\theta_{NBS}$ parameter for the BPP runs for Settings 7-9 in Table S2. The red line is the mean of the sampled values for that run.

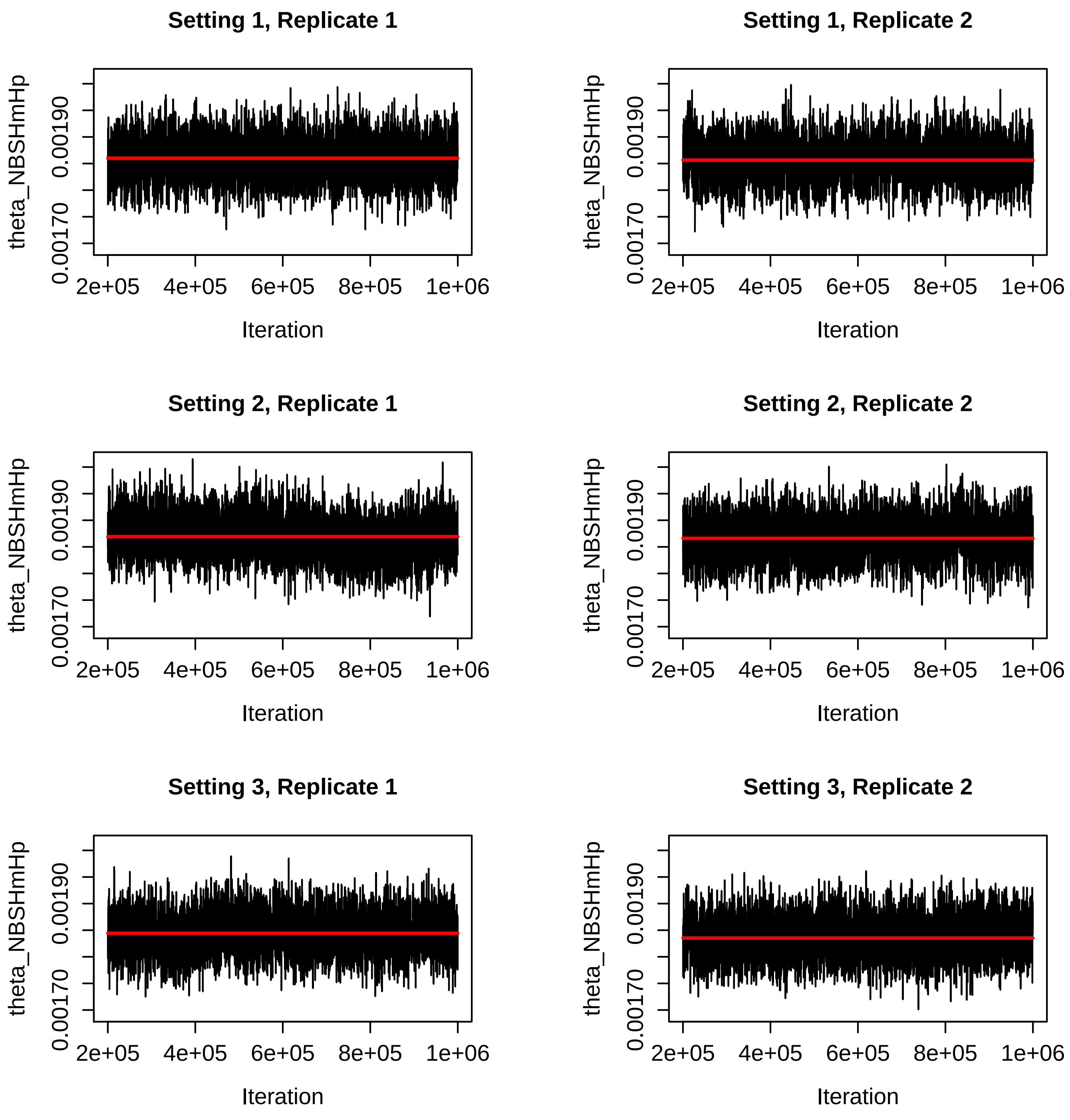


Figure S56: Trace plots for the $\theta_{NBSHmHp}$ parameter for the BPP runs for Settings 1-3 in Table S2. The red line is the mean of the sampled values for that run.

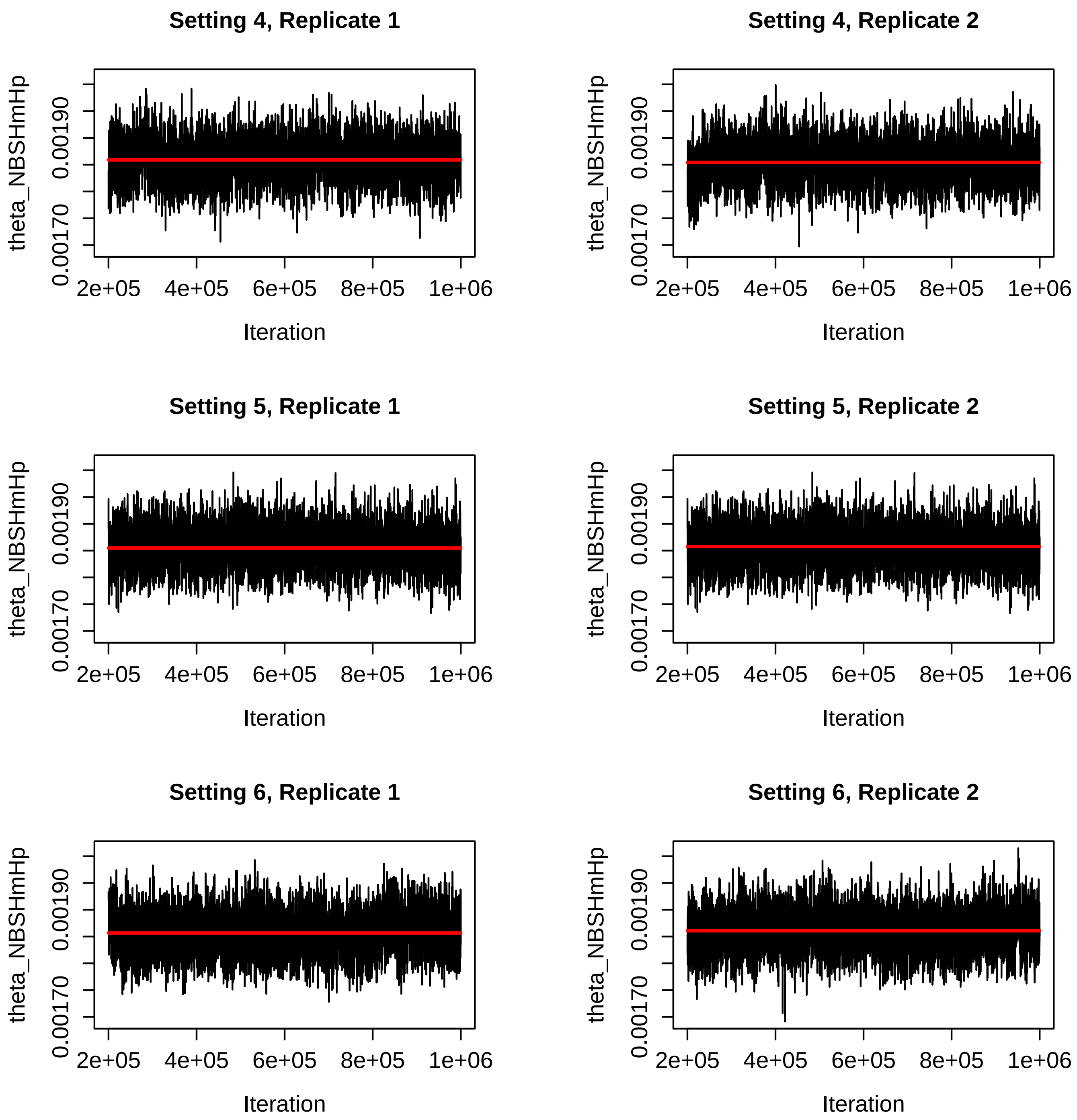


Figure S57: Trace plots for the $\theta_{NBSHmHp}$ parameter for the BPP runs for Settings 4-6 in Table S2. The red line is the mean of the sampled values for that run.

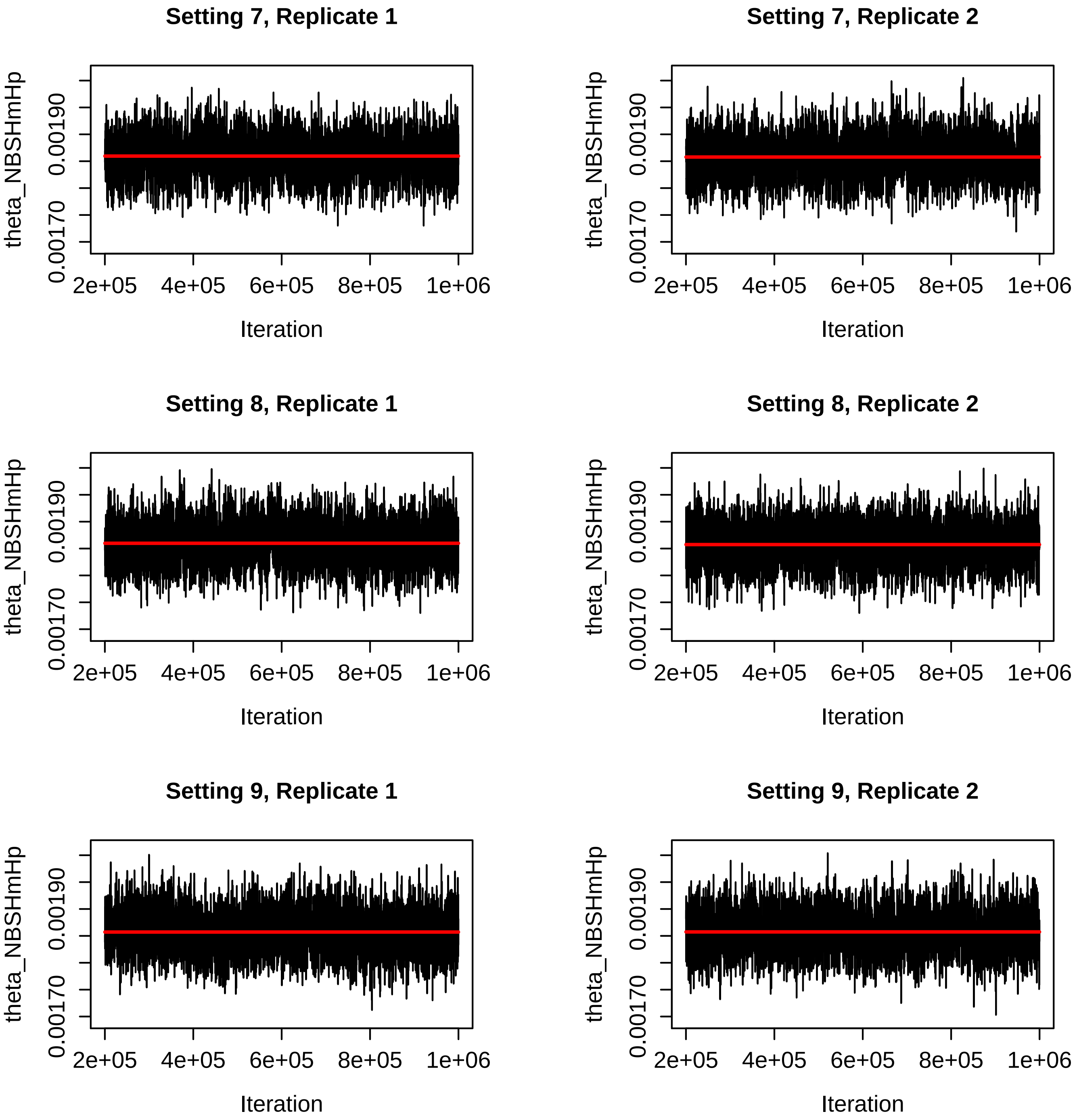


Figure S58: Trace plots for the $\theta_{NBSHmHp}$ parameter for the BPP runs for Settings 7-9 in Table S2. The red line is the mean of the sampled values for that run.

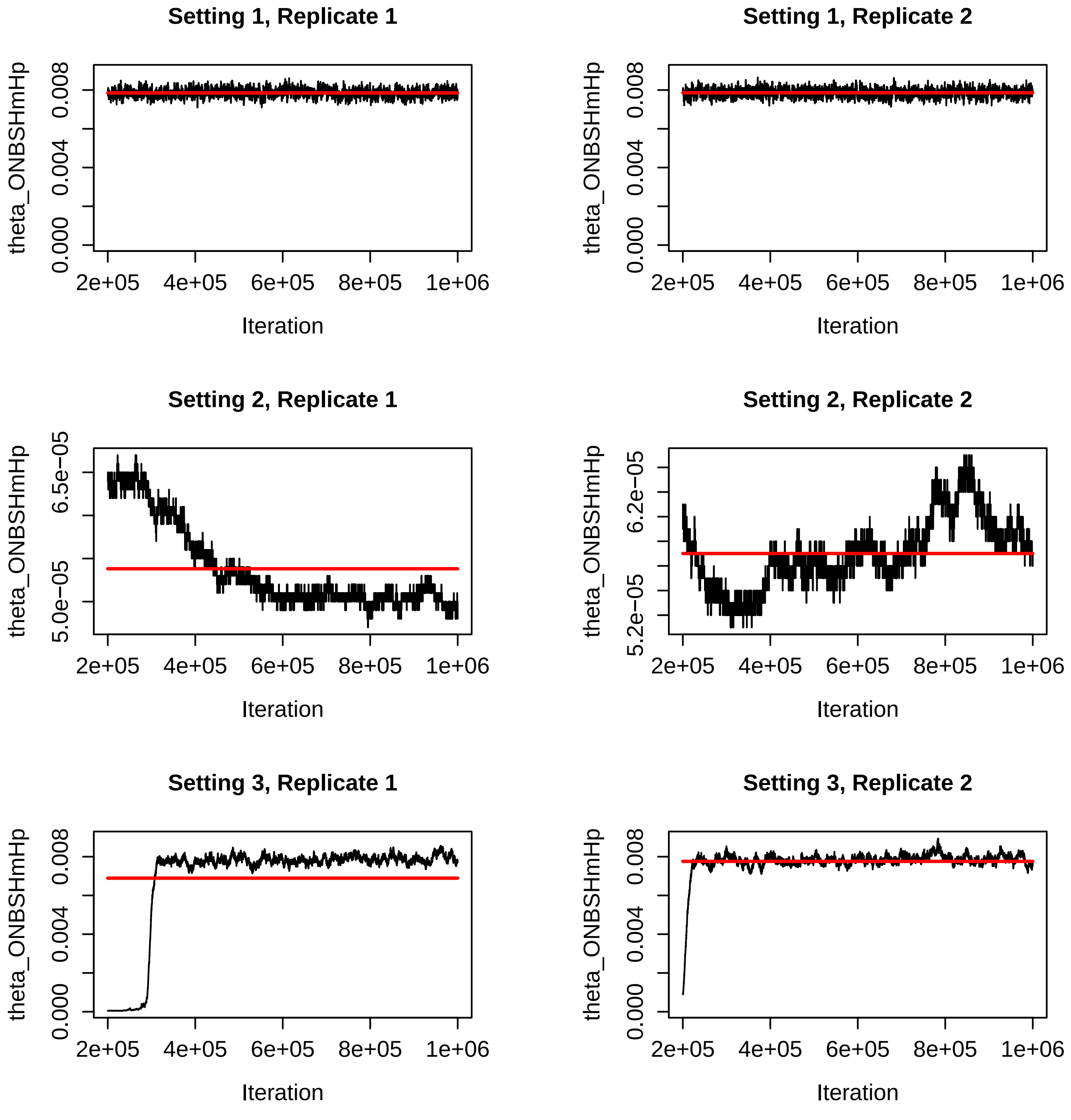


Figure S59: Trace plots for the $\theta_{ONBSHmHp}$ parameter for the BPP runs for Settings 1-3 in Table S2. The red line is the mean of the sampled values for that run. Note the difference in the y-axis values for Setting 2, Replicate 1.

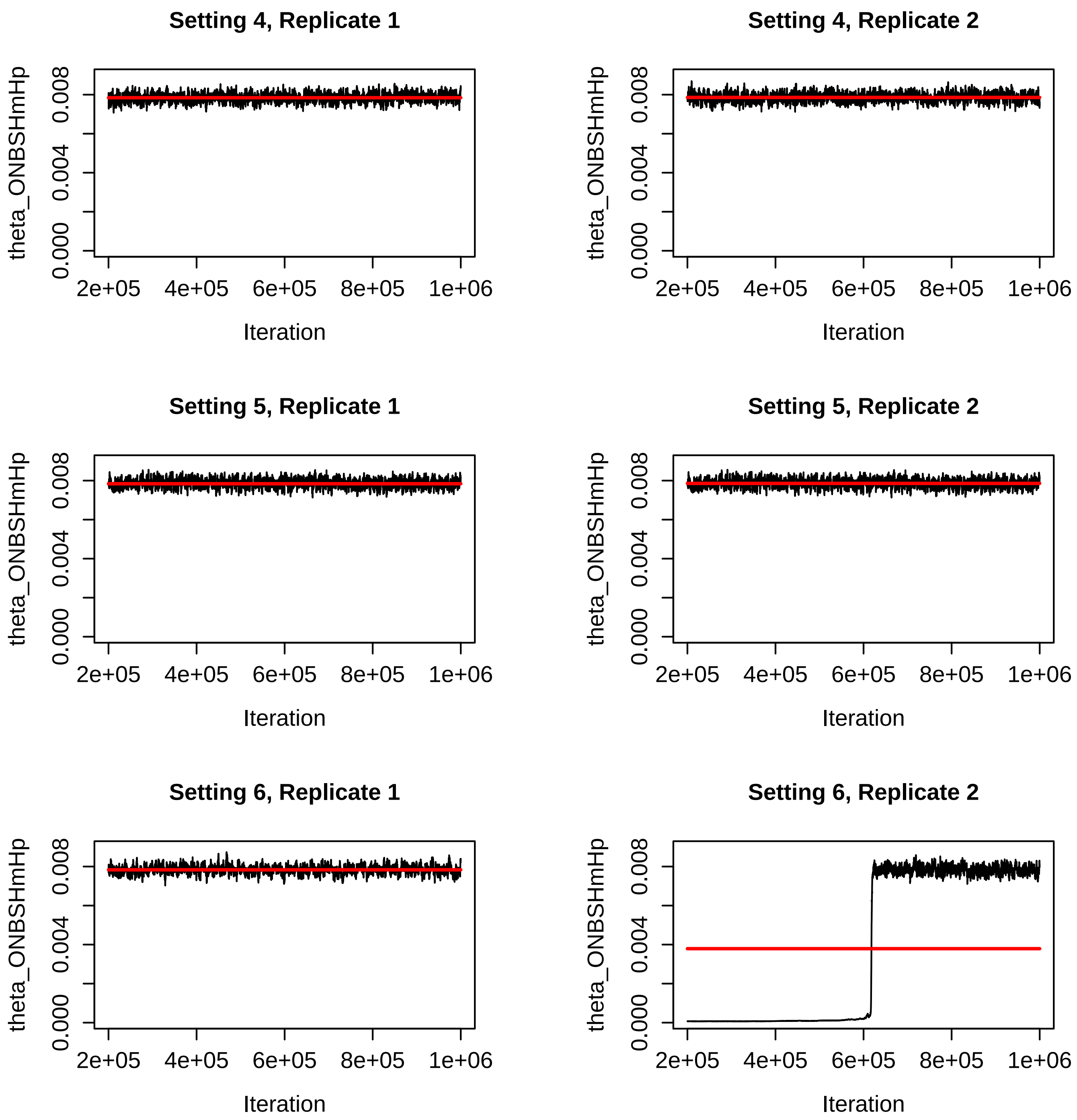


Figure S60: Trace plots for the $\theta_{ONBSHmHp}$ parameter for the BPP runs for Settings 4-6 in Table S2. The red line is the mean of the sampled values for that run.

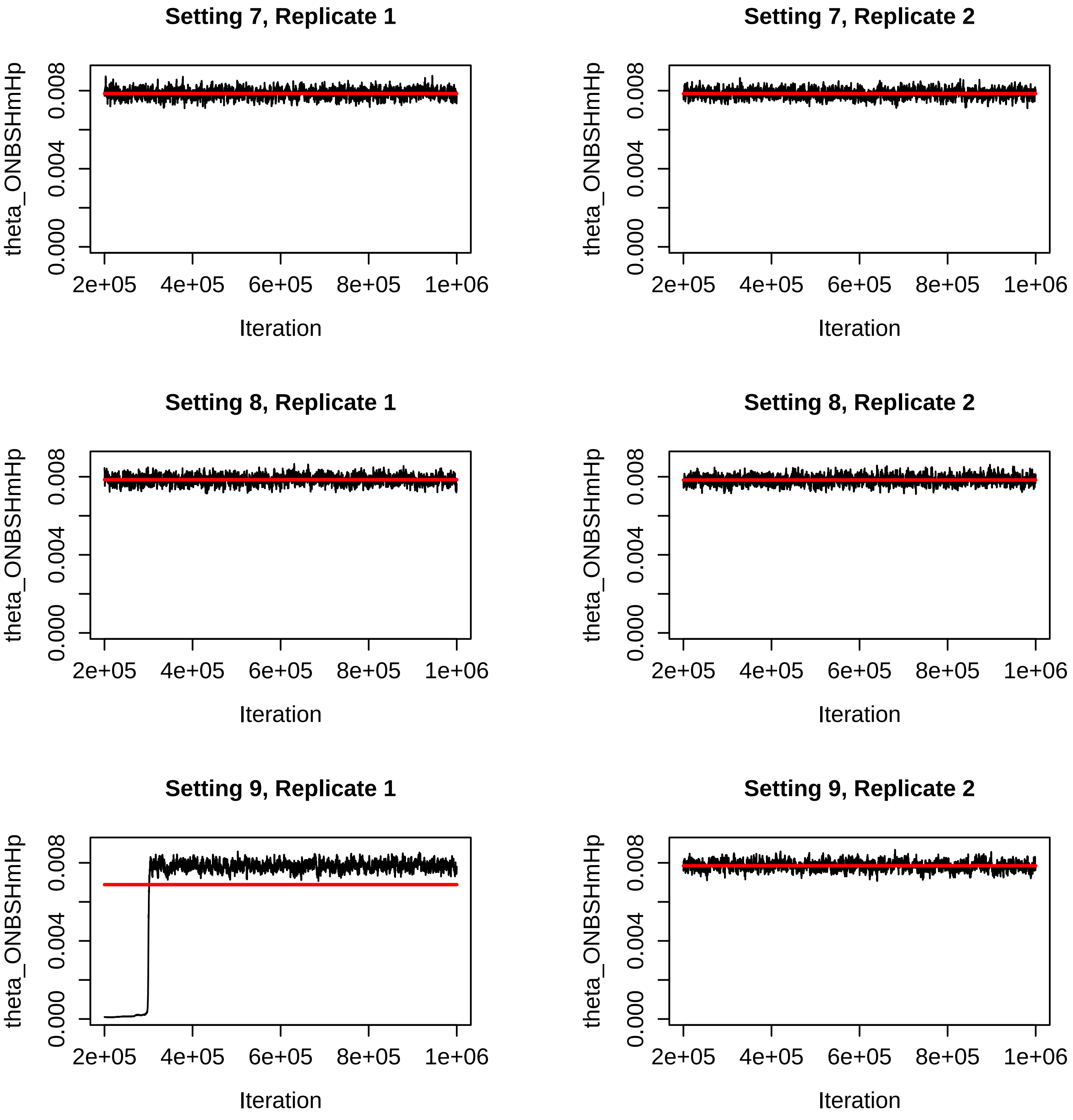


Figure S61: Trace plots for the $\theta_{ONBSHmHp}$ parameter for the BPP runs for Settings 7-9 in Table S2. The red line is the mean of the sampled values for that run.

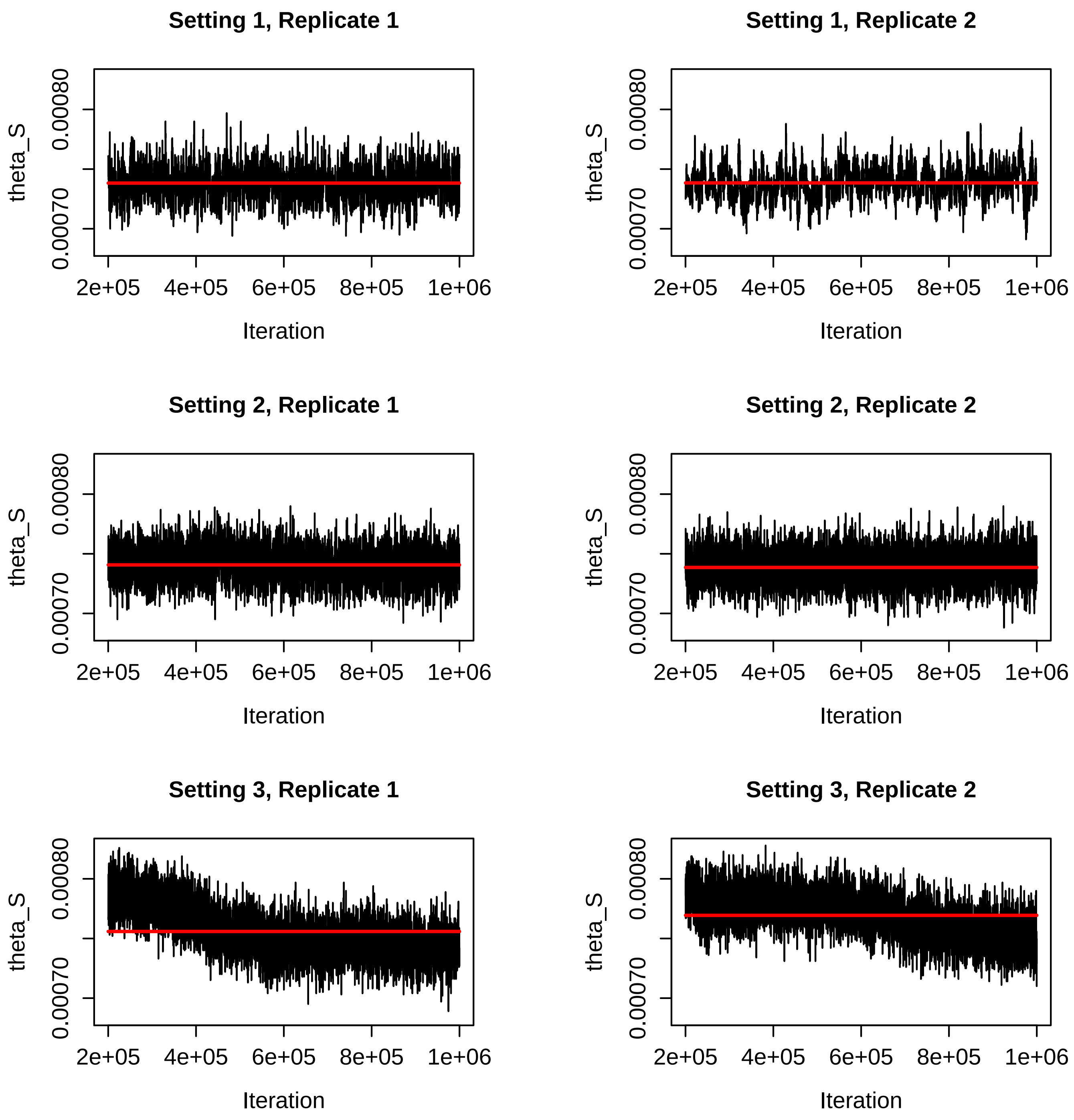


Figure S62: Trace plots for the $\theta_S$ parameter for the BPP runs for Settings 1-3 in Table S2. The red line is the mean of the sampled values for that run.

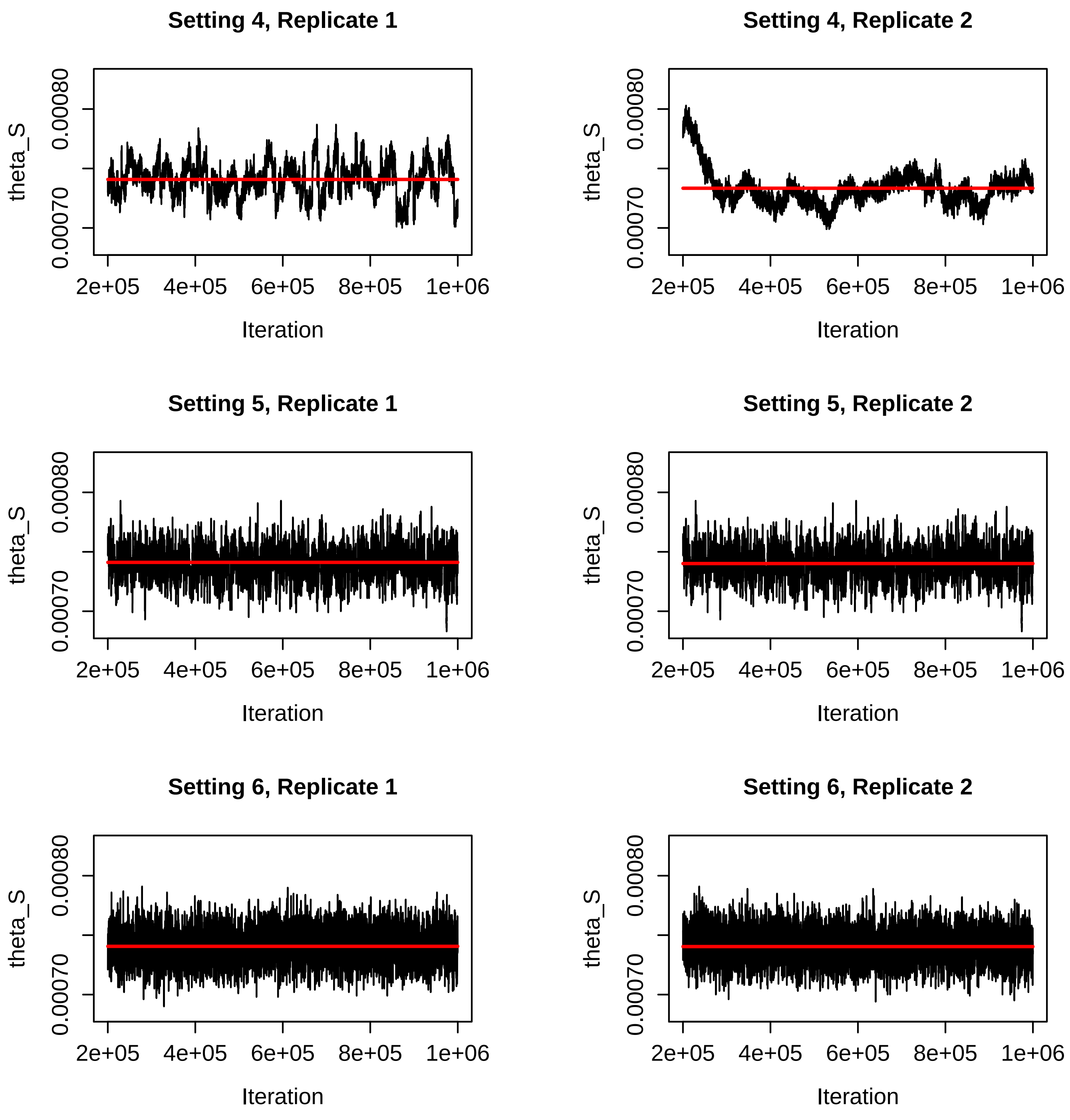


Figure S63: Trace plots for the $\theta_S$ parameter for the BPP runs for Settings 4-6 in Table S2. The red line is the mean of the sampled values for that run.

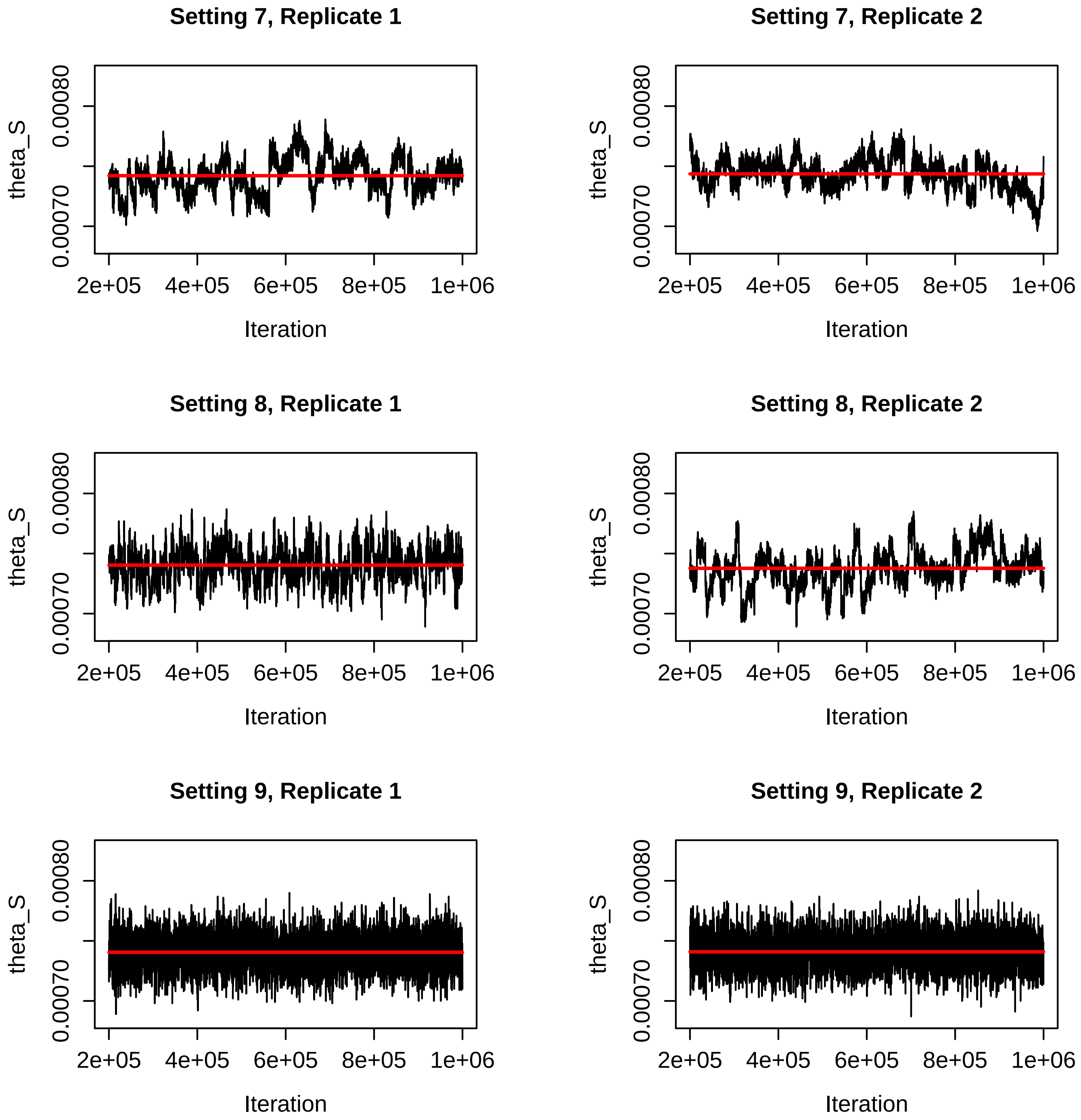


Figure S64: Trace plots for the $\theta_S$ parameter for the BPP runs for Settings 7-9 in Table S2. The red line is the mean of the sampled values for that run.

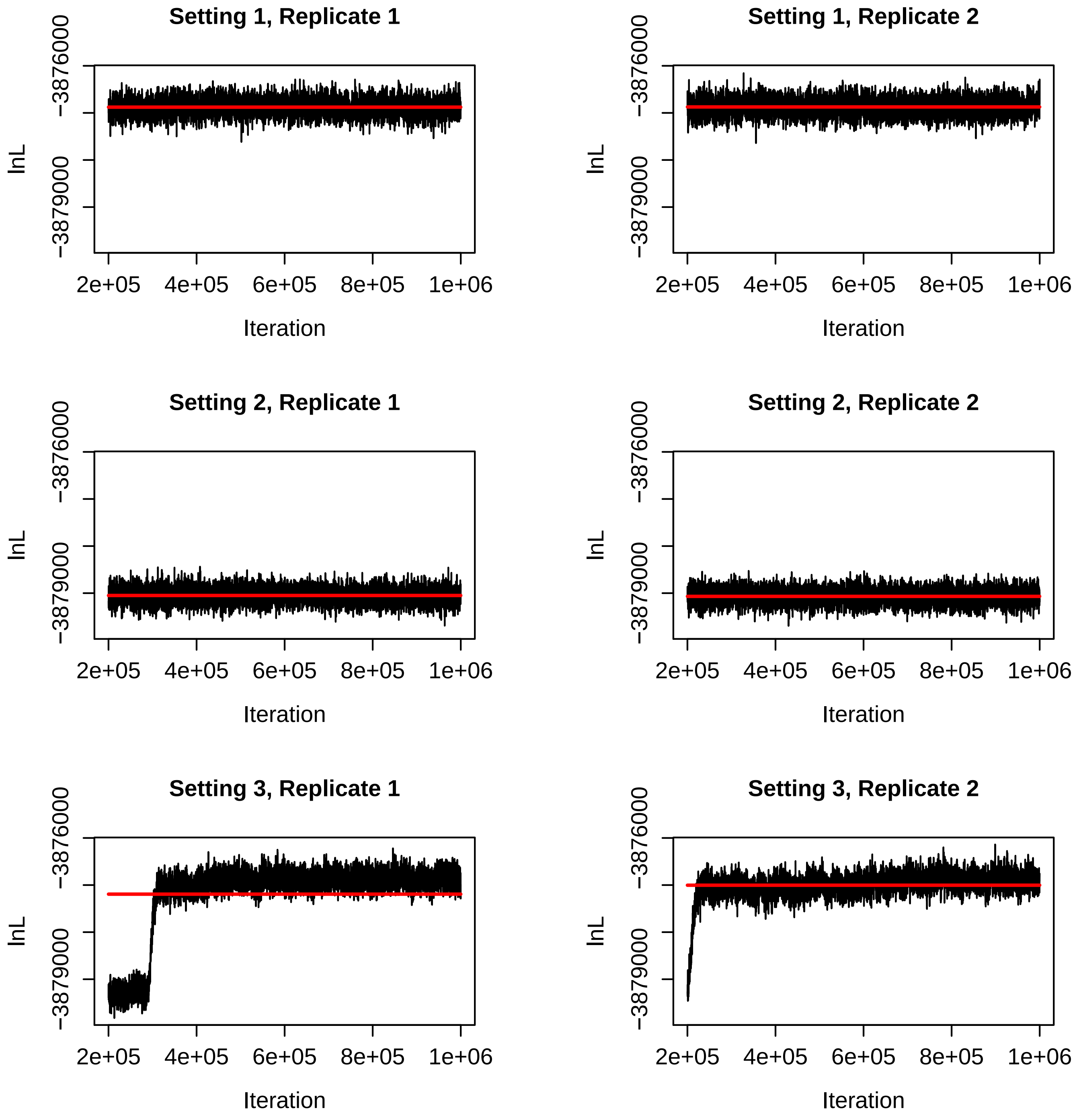


Figure S65: Trace plots for the log likelihood for the BPP runs for Settings 1-3 in Table S2. The red line is the mean of the sampled values for that run.

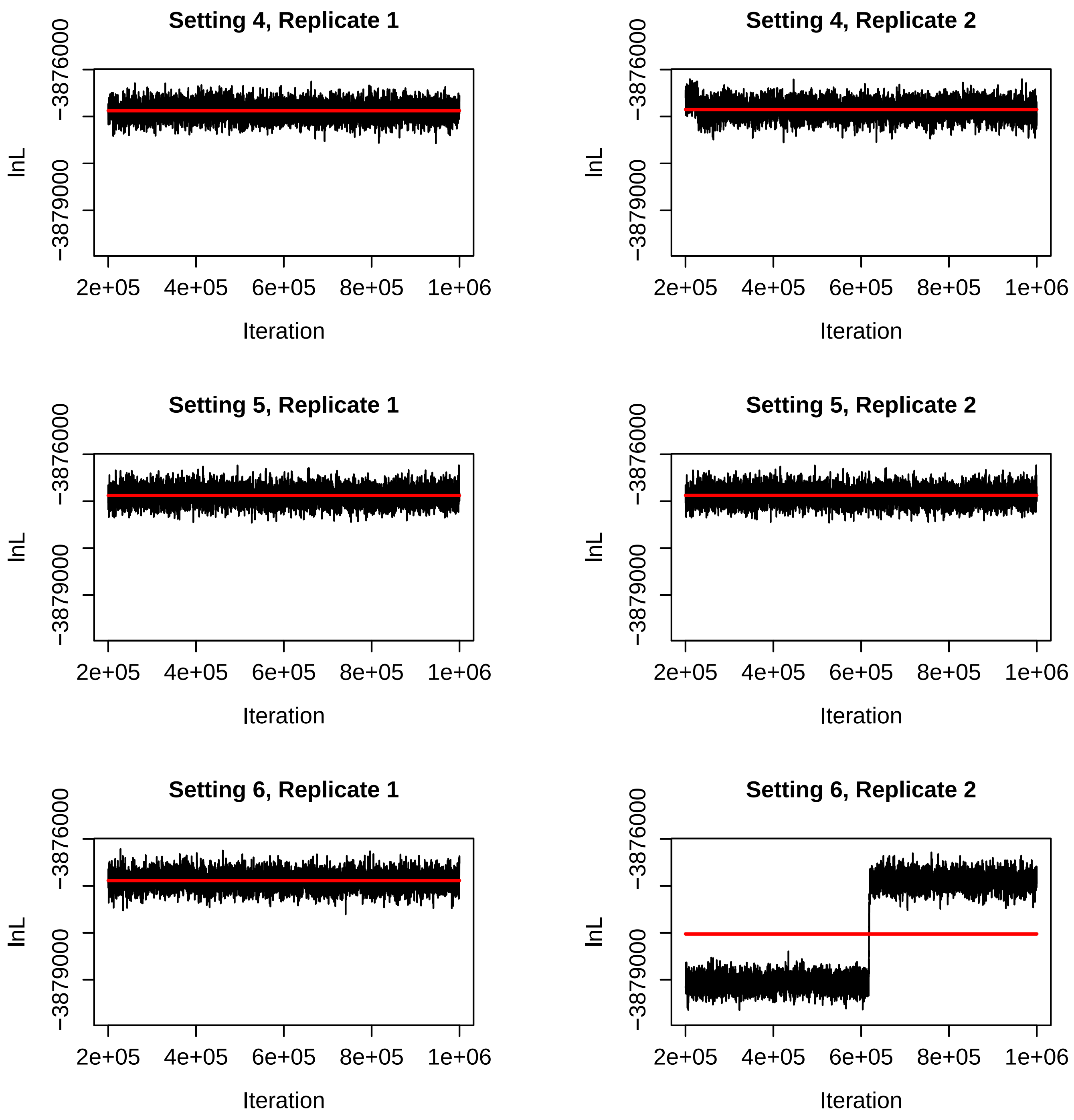


Figure S66: Trace plots for the log likelihood for the BPP runs for Settings 4-6 in Table S2. The red line is the mean of the sampled values for that run.

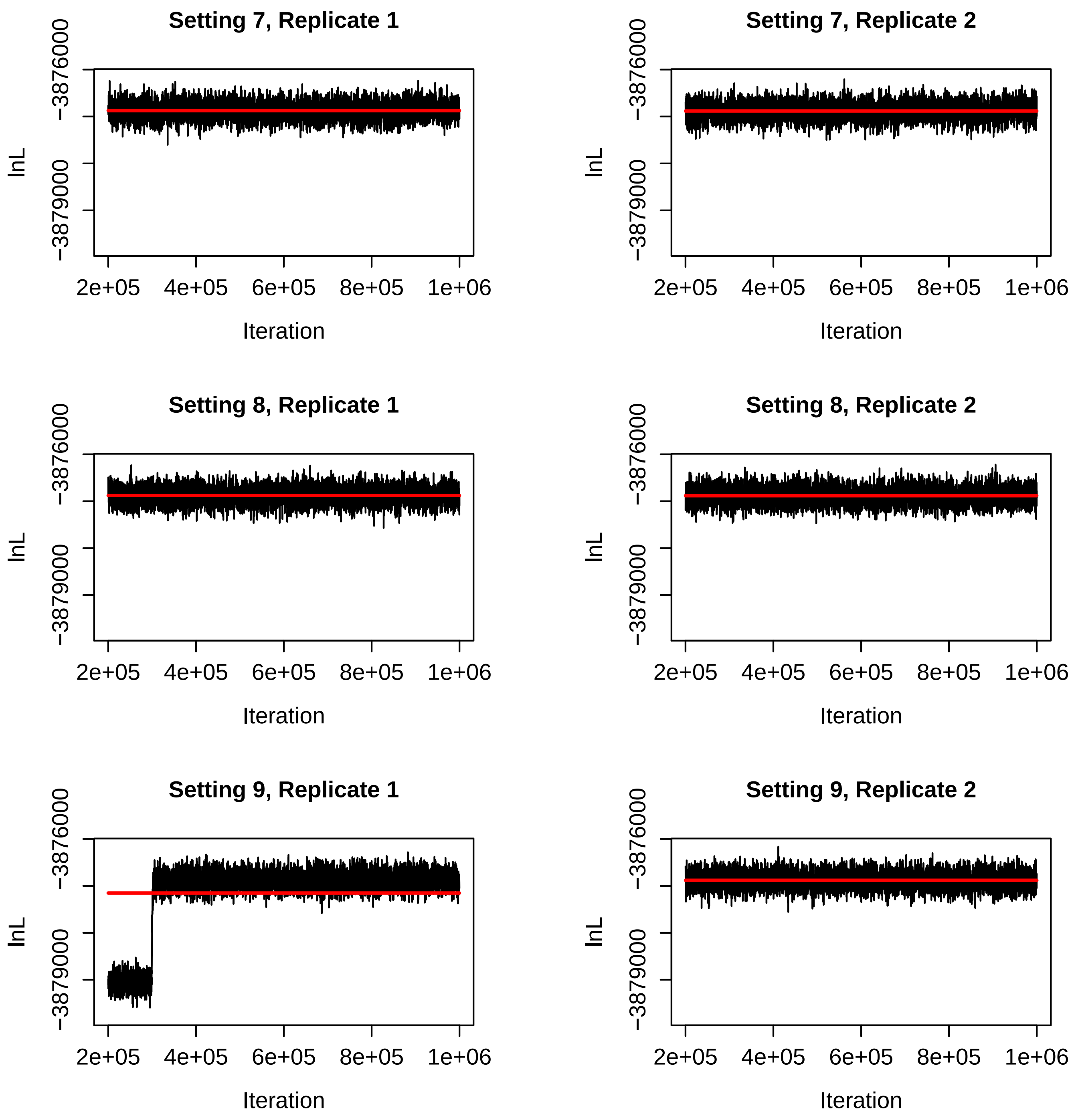


Figure S67: Trace plots for the log likelihood for the BPP runs for Settings 7-9 in Table S2. The red line is the mean of the sampled values for that run.
